# Rewarding capacity of optogenetically activating a giant GABAergic central-brain interneuron in larval *Drosophila*

**DOI:** 10.1101/2022.12.19.521052

**Authors:** Nino Mancini, Juliane Thoener, Esmeralda Tafani, Dennis Pauls, Oded Mayseless, Martin Strauch, Katharina Eichler, Andrew Champion, Oliver Kobler, Denise Weber, Edanur Sen, Aliće Weiglein, Volker Hartenstein, Andreas S. Thum, Astrid Rohwedder, Michael Schleyer, Bertram Gerber

**Affiliations:** Leibniz Institute for Neurobiology (LIN), Department Genetics of Learning and Memory, 39118 Magdeburg, Germany; Department of Animal Physiology, Institute of Biology, Leipzig University, 04103 Leipzig, Germany; Department of Genetics, Institute of Biology, Leipzig University, 04103 Leipzig, Germany; Department of Molecular Cell Biology, Weizmann Institute of Science, Rehovot 7610001 Israel; RWTH Aachen University, Institute of Imaging and Computer Vision, 52074 Aachen, Germany; Institute of Neurobiology, University of Puerto Rico Medical Science Campus, Old San Juan, Puerto Rico 00901; Department of Physiology, Development and Neuroscience, Cambridge University, Cambridge, CB2 3EL, United Kingdom; Janelia Research Campus, Howard Hughes Medical Institute, Ashburn, Virginia, USA; Leibniz Institute for Neurobiology (LIN), Combinatorial Neuroimaging Core Facility (CNI), 39118 Magdeburg, Germany; University of California, Department of Molecular, Cell and Developmental Biology, CA 90095-1606, Los Angeles, USA; Center for Behavioral Brain Sciences (CBBS), Magdeburg, Germany.; Institute for Biology, Otto von Guericke University, 39106 Magdeburg, Germany

**Keywords:** reward, dopamine, mushroom body, optogenetics, olfaction

## Abstract

Larvae of the fruit fly *Drosophila melanogaster* are a powerful study case for understanding the neural circuits underlying behavior. Indeed, the numerical simplicity of the larval brain has permitted the reconstruction of its synaptic connectome, and genetic tools for manipulating single, identified neurons allow neural circuit function to be investigated with relative ease and precision. We focus on one of the most complex neurons in the brain of the larva (of either sex), the GABAergic anterior paired lateral neuron (APL). Using behavioral and connectomic analyses, optogenetics, Ca^2+^ imaging and pharmacology, we study how APL affects associative olfactory memory. We first provide a detailed account of the structure, regional polarity, connectivity, and metamorphic development of APL, and further confirm that optogenetic activation of APL has an inhibiting effect on its main targets, the mushroom body Kenyon cells. All these findings are consistent with the previously identified function of APL in the sparsening of sensory representations. To our surprise, however, we found that optogenetically activating APL can also have a strong rewarding effect. Specifically, APL activation together with odor presentation establishes an odor-specific, appetitive, associative short-term memory, whereas naïve olfactory behavior remains unaffected. An acute, systemic inhibition of dopamine synthesis as well as an ablation of the dopaminergic pPAM neurons impair reward learning through APL activation. Our findings provide a study case of complex circuit function in a numerically simple brain, and suggest a previously unrecognized capacity of central-brain GABAergic neurons to engage in dopaminergic reinforcement.

**Significance statement:** The single, identified giant anterior paired lateral (APL) neuron is one of the most complex neurons in the insect brain. It is GABAergic and contributes to the sparsening of neuronal activity in the mushroom body, the memory center of insects. We provide the most detailed account yet of the structure of APL in larval *Drosophila* as a neurogenetically accessible study case. We further reveal that, contrary to expectations, the experimental activation of APL can exert a rewarding effect, likely via dopaminergic reward pathways. The present study both provides an example of unexpected circuit complexity in a numerically simple brain, and reports an unexpected effect of activity in central-brain GABAergic circuits.

## Introduction

Larvae of the fruit fly *Drosophila melanogaster*, which naturally live on overripe fruit, provide a powerful study case for investigating the neurogenetic bases of learning and memory (Gerber and Stocker, 2007; Widmann et al., 2018; Thum and Gerber, 2019; Eschbach and Zlatic, 2020). Their small size and low number of neurons have allowed their chemical synapse connectome to be reconstructed, revealing unexpected complexity. In the mushroom body, which is a higher brain structure for sensory integration and memory in insects (Heisenberg, 2003), more than half of the classes of synaptic connections had previously escaped attention (Figure 1D; Eichler et al., 2017; Eschbach et al., 2020, 2021; adults: Takemura et al., 2017; Li et al., 2020). For instance, dopaminergic mushroom body input neurons (DANs) not only relay ascending information to local compartments along the elongated axonal fibers of the mushroom body intrinsic Kenyon cells (KCs), but also integrate local information from the KCs and recurrent signals originating from mushroom body output neurons (MBONs; these likewise respect compartmental boundaries). Similar complexity is observed for octopaminergic input neurons (OANs) and for input neurons using as yet unidentified signaling (Eichler et al., 2017; Saumweber et al., 2018; Eschbach et al., 2020, 2021; Schleyer et al., 2020). Collectively, this should prepare us for more surprises regarding mushroom body function. Here we study one of the most complex mushroom body neurons of larvae, the anterior paired lateral (APL) neuron.

**Figure 1.**
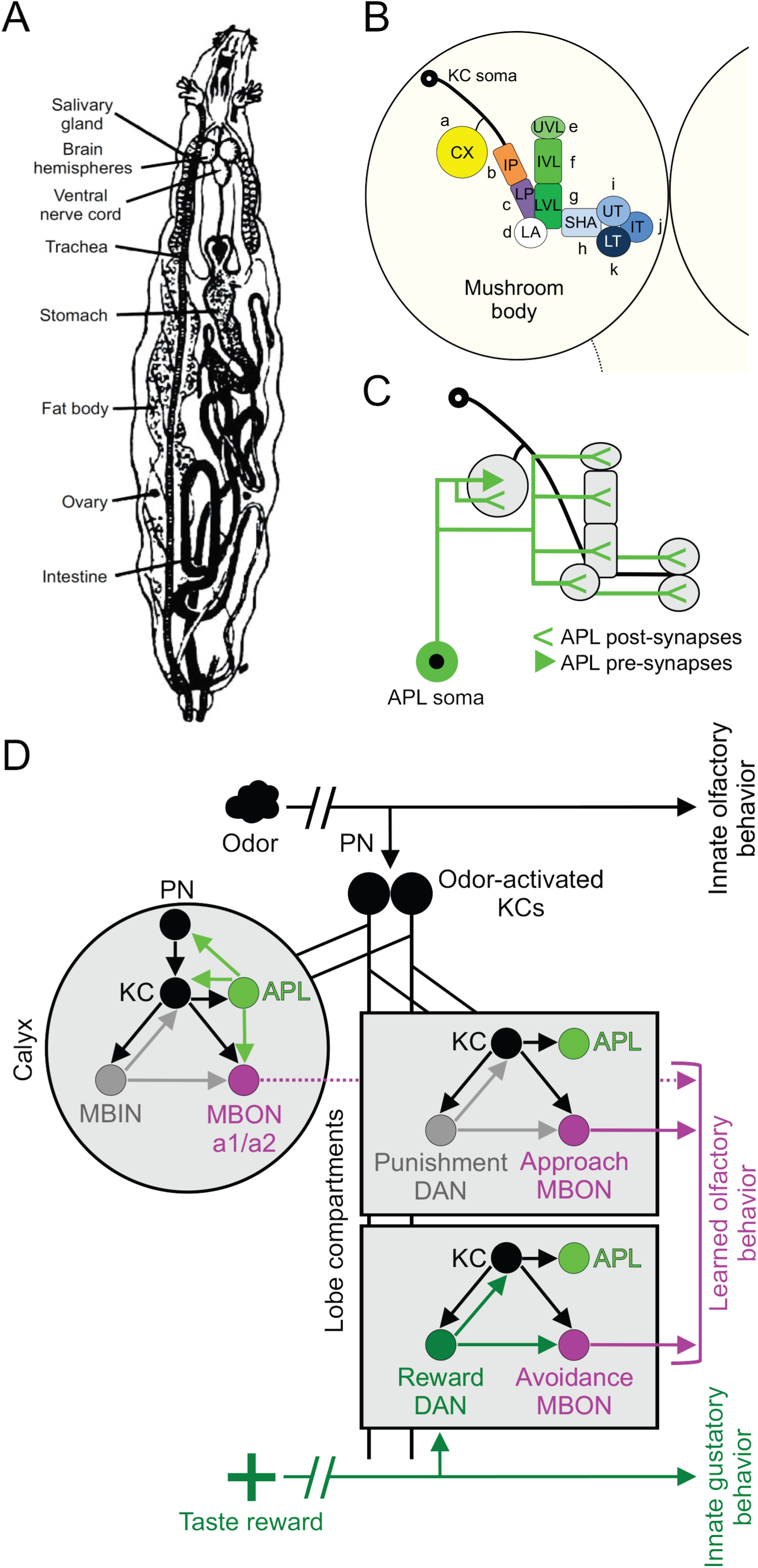
Overview of the larval body and brain and connectivity of the APL neuron in the mushroom body. **(A)** Schematic overview of the larval body, adapted from Demerec and Kaufmann (1940). **(B)** Sketch of one larval brain hemisphere with the mushroom body, highlighting its intrinsic Kenyon cells (KCs) and organization in 11 compartments (letters a-k refer to the suffixes used to indicate the compartments innervated by mushroom body extrinsic neurons): CX: calyx; IP and LP: intermediate and lower peduncle; LA: lateral appendix; UVL, IVL and LVL: upper, intermediate, and lower vertical lobe; SHA: shaft; UT, IT, LT: upper, intermediate, and lower toe of the medial lobe. Adapted from Saumweber et al. (2018). **(C)** The larval anterior paired lateral (APL) neuron collects input (<) mostly from the KCs both in the calyx and in a subset of the compartments in the lobes, and delivers output (arrowhead) mostly to KCs and almost exclusively in the calyx. Adapted from Saumweber et al. (2018). **(D)** Simplified diagram of the connectivity of APL and of circuits underlying associative odor-reward learning. Within the calyx (gray-filled circle), a given odor (cloud) leads to the activation of a sparse, odor-specific pattern of KCs established through input from the projection neurons (PN). Within the lobes (bottom gray-filled rectangle), modulatory dopaminergic neurons (reward DAN) carry taste reward signals to the KCs, which send their axonal projection to avoidance-promoting mushroom body output neurons (avoidance MBON). In addition to its connections with the KCs, the APL neuron establishes synaptic contacts with the calyx MBONs (MBON-a1 and - a2) as well as with a subset of PNs (for additional connections between APL and mushroom body extrinsic neurons that are omitted here see Figures 6-7). During odor-taste reward associative learning, the coincidence between the odor and the reward signal at the KCs is thought to lead to a pre-synaptic depression of the synapses between the odor-activated KCs and avoidance-promoting MBONs, whereas the synapses of these KCs with approach- promoting MBONs in other compartments remains unchanged (note that the contribution of MBON-a1/a2 to learned behavior is unclear, as indicated by the stippled lines). Future processing of the learned odor is thus biased in favor of approach. The same rationale is thought to apply for odor-punishment learning, occurring at the synapses between the KCs and approach MBONs (top gray-filled rectangle). The electron microscopy reconstruction of a first-instar larval nervous system additionally revealed unexpected connections from KCs towards mushroom body input neurons (MBINs) including DANs, as well as MBIN/DAN-to- MBON synapses; note that KC-to-KC and MBON-to-MBIN connections are not displayed. Arrows indicate synaptic contacts between neurons.

APL is a hemispherically unique local interneuron and can be identified from the earliest larval stage on, throughout metamorphosis and in adults (Eichler et al., 2017; Mayseless et al., 2018; Saumweber et al., 2018). It receives most of its input from, and provides GABAergic output to, the cholinergic KCs, suggesting a role in sparsening sensory representation within the mushroom body (Masuda-Nakagawa et al., 2014; adults: Honegger et al., 2011; Lin et al., 2014; Amin et al., 2020; Prisco et al., 2021; further insects: Homberg et al., 1987; Grünewald, 1999; Papadopoulou et al., 2011). In contrast to most other aspects of mushroom body connectivity, however, there are major differences in APL connectivity between larvae and adults.

In adults, APL innervates all 15 mushroom body compartments and the calyx, where the KCs receive input from sensory projection neurons (Tanaka et al., 2008; Aso et al., 2014). In larvae, APL also innervates the calyx, but only six of the 10 compartments (Figure 1B; Eichler et al., 2017; Saumweber et al., 2018).

In adults, APL connects reciprocally with the KCs in the calyx and in all the compartments (Wu et al., 2013; Takemura et al., 2017; Zheng et al., 2018; Scheffer et al., 2020), whereas in larvae such reciprocal connections exist only in the calyx, and only KC-to-APL synapses are found otherwise (Masuda-Nakagawa et al., 2014; Eichler et al., 2017; Saumweber et al., 2018).

In adults, APL is electrically coupled to the dorsal paired median neuron (DPM), a local interneuron that innervates all the compartments but not the calyx (Pitman et al., 2011; Wu et al., 2011). DPM is serotonergic, co-releases GABA, and can express the amnesiac peptide (Waddell et al., 2000; Lee et al., 2011; Haynes et al., 2015; Turrel et al., 2018). Strikingly, DPM is absent in larvae (Eichler et al., 2017; Saumweber et al., 2018).

These differences caution against extrapolating between findings on APL in larvae and adults, since functions of APL other than a sparsening of KC activity have been described in adults (Liu et al., 2007; Liu and Davis, 2009; Ren et al., 2012; Wu et al., 2012; Lin et al., 2014). In this context, we provide a comprehensive account of the structure of the larval APL neuron, the spatial arrangement of its synapses, its physiological effect on KC activity, and its metamorphic development. Investigating its role in Pavlovian conditioning, we discover that, surprisingly, optogenetic activation of APL exerts a rewarding effect. This effect is studied in detail and is shown to involve a dopamine-dependent process.

## Materials & Methods

### Drosophila strains

*Drosophila melanogaster* were kept and maintained on standard medium, in mass culture at 25°C, 60%–70% humidity, in a 12/12 h light/dark cycle. We used randomly chosen third-instar, feeding-stage larvae of both sexes, aged 5 days (120 h) after egg laying, unless mentioned otherwise. The strains used in this study and their genotypes are listed in Extended Data Table 2-1.

### Full-body fluorescence microscopy

To allow a full-body assessment of transgene expression from the APL-specific split-GAL4 SS01671 driver (henceforth APL-GAL4; Saumweber et al., 2018), it was crossed to w*; UAS- mCherryCAAX (abbreviated as UAS-mCherry-CAAX; Sens et al., 2010; Bloomington Stock Center no. 59021) to express the mCherry-CAAX reporter. Double-heterozygous third-instar progeny (abbreviated as APL>mCherry-CAAX) were analyzed for fluorescence signals under a light-sheet microscope (see next paragraph). Genetic controls were heterozygous for either the GAL4 element (APL>+) or the UAS element (+>mCherry-CAAX). To obtain the driver control, APL-GAL4 was crossed to y^1^w^1^ (Bloomington Stock Center no. 1495). As regards the effector control, a strain lacking the GAL4 domains but containing the two split-GAL4 landing sites (attP40/attP2; Pfeiffer et al., 2010) was crossed to UAS-mCherry-CAAX.

Experimental procedures follow Kobler et al. (2021). In brief, third-instar larvae were first bleached (4% sodium hypochlorite, Roth, order no. 9062.1) for 10 min at room temperature. After washing (3 x 5 min in distilled water, dH_2_O), they were fixed in 4% paraformaldehyde (PFA) in 0.1 M phosphate buffer (PB) at pH 9, with gentle shaking overnight at 4°C. Fixed samples were briefly rinsed 3 x with 0.1 M PB containing 0.2% Triton-X-100 (PBT), then washed 2 x 60 min and left overnight at 4°C in PBT at pH 9. Dehydration the following day used a graded ethanol series (60 min in 10% and 25% ethanol, followed by 30 min in 50%, 60%, 80% (all at pH 9), and 2 x 100% ethanol). Samples were then cleared by replacing ethanol with ethyl cinnamate (ECi; Ethyl 3-phenyl-2-propenoate; Sigma-Aldrich, order no. 112372-100G). After 30-60 min, the ECi was refreshed once, and the samples stored at room temperature in black boxes in a desiccator.

The samples were placed ventral side up in ECi-cleared phytagel blocks (1 x 1 x 1 cm, Sigma-Aldrich, P8169). These were placed in a sample holder, which in turn was fixed on a mounting suspension to fit into a high-precision quartz glass cuvette filled with ECi and optically accessible with an UltraMicroscope II light-sheet microscope (Miltenyi Biotec). The microscope was equipped with a Zyla 4.2 PLUS sCMOS camera (Oxford Instruments, Abingdon-on-Thames, UK) and a tube for infinity-corrected objective lenses. An EXW-12 laser (NKT Photonics, Birkerød, Denmark) was used for excitation through triple-sheet optics to illuminate samples from one side. Excitation and emission filters (AHF Analysentechnik AG) were used as indicated in Figure 2 and Movie 1.

**Figure 2:**
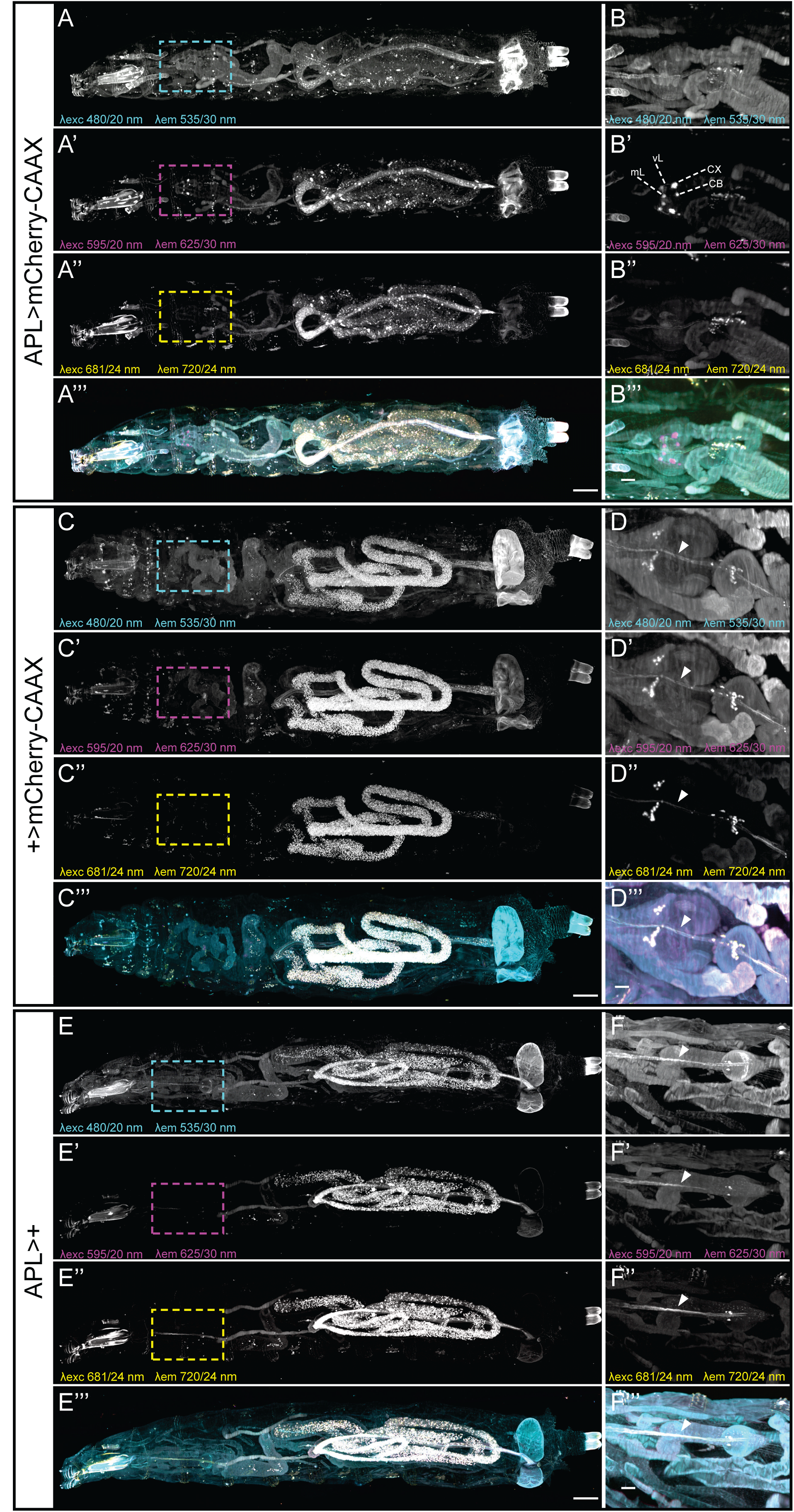
The APL-GAL4 driver does not express outside the brain. **(A-B’’’)** Maximum intensity projection of fluorescence signals from an entire larva of the genotype APL>mCherry-CAAX acquired on a light-sheet microscope using a 12x objective (top view, rostral to the left). Dashed boxes indicate the central nervous system, shown enlarged for a volume-restricted view in (B-B‘‘‘). λexc and λem indicate filter band passes used for excitation and emission, respectively (merged in A’’’ and B’’’). CX: mushroom body calyx; CB: cell body of the APL neuron; vL and mL: innervation of APL in the vertical and medial lobe of the mushroom body, respectively. Scale bars: 200 µm (A-A‘‘‘), 50 µm (B-B‘‘‘). See also Movie 2. **(C-F’’’)** As in A-B’’’, but for the effector control (+>mCherry-CAAX) (C-D’’’) and driver control (APL>+) (E-F’’’), respectively. The arrowhead points to autofluorescence signals from the pharynx (visible also in B-B’’’ but omitted for clarity). Fluorescence signals that can be observed across wavelengths and genotypes reflect autofluorescence (including from food particles in the gut) and allow bodily detail to be discerned. Fluorescence reflecting the expression of mCherry-CAAX was observed only in the brain and only the APL neuron (magenta signals in A’’’, B’’’, Movie 1) (compare B’, D’, F’).

Tiled image stacks were acquired with a LVMI-Fluor 12x objective (Miltenyi Biotec) with a format of 2048 x 2048. Using ImSpector software (version 7.1.4, Miltenyi Biotec), a 10% overlap of tiles was set for stitching.

Image files were processed with Imaris software (version 9.8, Bitplane), including file conversion, stitching, further processing, and rendering. The *ortho slicer* tool was used to restrict volumes in the z-direction to improve the representation of structures that were otherwise covered in the context of the whole body. All 3D images were generated using the *snapshot* function. 2D maximum-intensity projections were generated in Fiji (Schindelin et al., 2012).

Movie 1 was produced in Imaris with the *key frame animation* tool; Adobe Premiere Pro 2020 (version 14.9.0, Adobe Inc) was used for cutting and labeling.

### Immunohistochemistry

All the antibodies used in this study are listed in Extended Data Table 2-1.

#### Transgene expression pattern of the SS01671 driver strain

To validate specific expression in the larval APL neuron of the APL-GAL4 driver strain (Saumweber et al., 2018), it was crossed to UAS-ChR2XXL::tdtomato to express a tomato-tagged version of ChR2XXL (FlyBase ID: FBtp0131815; Saumweber et al., 2018). Double-heterozygous third-instar progeny (abbreviated as APL>ChR2XXL::tdtomato) were dissected in ice-cold Ringer’s solution, and the brains were fixed for 30 min in 10% formaldehyde dissolved in phosphate buffered saline (PBS, pH 7.2, P4417, Sigma Aldrich) at room temperature. After consecutive washing steps (3 x 10 min each) in PBT (0.3% Triton-X-100 [CAS: 9036-19-5, Roth] in PBS), the brains were blocked in 5% normal goat serum solution (NGS; 005-000-121, Jackson Immunoresearch Laboratories; in PBS) for 2 h at room temperature. To provide a reference staining of fiber tracts (including the mushroom bodies), tissues were incubated overnight at 4°C with a primary monoclonal mouse anti-FASII antibody (AB_528235, DSHB) diluted 1:50 in blocking solution containing 4% NGS in PBS. After six washes (10 min each) in PBS, the tissues were treated overnight at 4°C with a secondary polyclonal goat anti-mouse Alexa Fluor 488 antibody (A11001, Invitrogen) diluted 1:200 in PBS. The brains were then washed in PBS (6 x 10 min each) and mounted in Vectashield (Vector Laboratories Inc) on a cover slip. Signal detection from the tomato-tag of ChR2XXL (labeling the APL neuron) did not require antibodies; rather, the tomato fluorescence signal was detected directly under the microscope. Image z-stacks were acquired with a Leica TCS SP8 confocal microscope (Leica Mikrosysteme Vertrieb GmbH) at a format of 1024 × 1024. Image processing was performed using Imaris software (version 9.72, Bitplane).

To visualize the larval APL neuron together with the mushroom bodies, we crossed the APL-GAL4 driver with a recombined effector/enhancer>effector strain that includes a UAS-mIFP-T2A-HO1 effector (abbreviated as mIFP; Yu et al., 2015; Bloomington Stock Center no. 64181) recombined with the enhancer>effector construct MB247>mCherry-CAAX (Kobler et al., 2021). Third-instar larval progeny (abbreviated as APL>mIFP/MB247>mCherry-CAAX) were dissected in ice-cold Ca^2+^-free saline solution and fixed for 24 h in 4% paraformaldehyde (PFA; J19943, Alfa Aesar; in PBS) at 4°C. After six washes (3 x brief; 3 x 10 min) in 0.3% PBT, the brains were mounted in Vectashield (Vector Laboratories Inc) on a cover slip. Signal detection from the mCherry-CAAX reporter (labeling the mushroom bodies) and the mIFP reporter (labeling APL) did not require antibodies for signal amplification. The image z-stack was acquired with a Leica TCS SP8 confocal microscope (Leica Mikrosysteme Vertrieb GmbH) at a format of 1024 × 1024. The corresponding Movie 2 was produced in Imaris (version 9.72, Bitplane).

To examine the inter-hemispheric symmetry in the morphology of APL, the APL-GAL4 driver was crossed to UAS-mCD8::GFP (Lee and Luo, 1999; Bloomington Stock Center no. 5137) as the effector. Third-instar larvae were put on ice and dissected in PBS. The brains were fixed in 4% PFA for 20 min at room temperature. After a succession of washing steps (3 x brief; 1 x 5 min; 3 x 15 min; 1 x 90 min) in 3% PBT (3% Triton-X-100 [CAS: 9002-93-1, Sigma Aldrich] in PBS) on ice, the brains were blocked with 5% NGS (G9023, Sigma Aldrich) in PBT for 1 h at room temperature and incubated for 48 h with primary antibodies at 4°C. The brains were then washed (2 x brief; 3 x 15 min; 1 x 60 min; on ice; 1 x 30 min at room temperature) in 3% PBT before application of the secondary antibodies for at least 24 h at 4°C. After a final set of washing steps (3 x brief; 3 x 5 min; 2 x 15 min) in 3% PBT, the brains were mounted on poly-L-lysin-coated cover slips (following the Janelia FlyLight recipe), dehydrated by a series of increasing concentrations of ethanol (EtOH) (1x brief in distilled water; 1 x 10 min 30% EtOH; 1 x 10 min 50% EtOH; 1 x 10 min 75% EtOH; 1 x 10 min 95% EtOH; 3 x 10 min 100% EtOH) and cleared (3 x 5 min) in xylene (247642, CAS: 1330-20-7, Sigma Aldrich). Finally, the brains were mounted in DPX mounting medium (dibutyl phthalate in xylene; 06522, Sigma Aldrich) and left in darkness for at least 24 h before imaging.

The primary antibody mixture consisted of (i) 2% NGS diluted 1:25 in 3% PBT, (ii) a polyclonal rabbit anti-GFP antibody (A6455, Life Technologies) diluted 1:1000 in 3% PBT (for APL staining), (iii) a monoclonal mouse 4F3 anti-DLG antibody (AB_528203, Developmental Studies Hybridoma Bank) diluted 1:200 in 3% PBT (for mushroom body staining), and (iv) a monoclonal rat anti-N-Cadherin antibody (DN-Ex #8-s, Developmental Studies Hybridoma Bank) diluted 1:50 in 3% PBT (for neuropil staining).

The secondary antibody mixture consisted of (i) 2% NGS diluted 1:25 in 3% PBT, (ii) polyclonal goat anti-rabbit Alexa Fluor 488 (A11008, Life Technologies), (iii) polyclonal goat anti-mouse Alexa Fluor 568 (A10037, Life Technologies), and (iv) polyclonal goat anti-rat Alexa Fluor 647 (712-605-153, Jackson ImmunoResearch), all diluted 1:500 in 3% PBT. Confocal microscopy was conducted on a Zeiss LSM800 confocal laser scanning microscope with ZEN 2.3 software. Image z-stacks were acquired with a LSM800 confocal microscope (Zeiss) at a format of 1024 × 1024. Image processing was performed using Imaris software (version 9.72, Bitplane).

To compare the coverage of the mushroom body compartments between the APL neuron of each hemisphere, mean pixel intensities were measured using ImageJ (version 1.53c, Fiji ImageJ). Grayscale maximum intensity projections of the GFP-channel (labeling APL membranes) were created, whereas the DLG-channel (labeling the mushroom body) served as a template for orientation. The mushroom body compartments were selected using the ROI Manager function; mean gray values are documented in Extended Data Figure 3-1.

**Figure 3.**
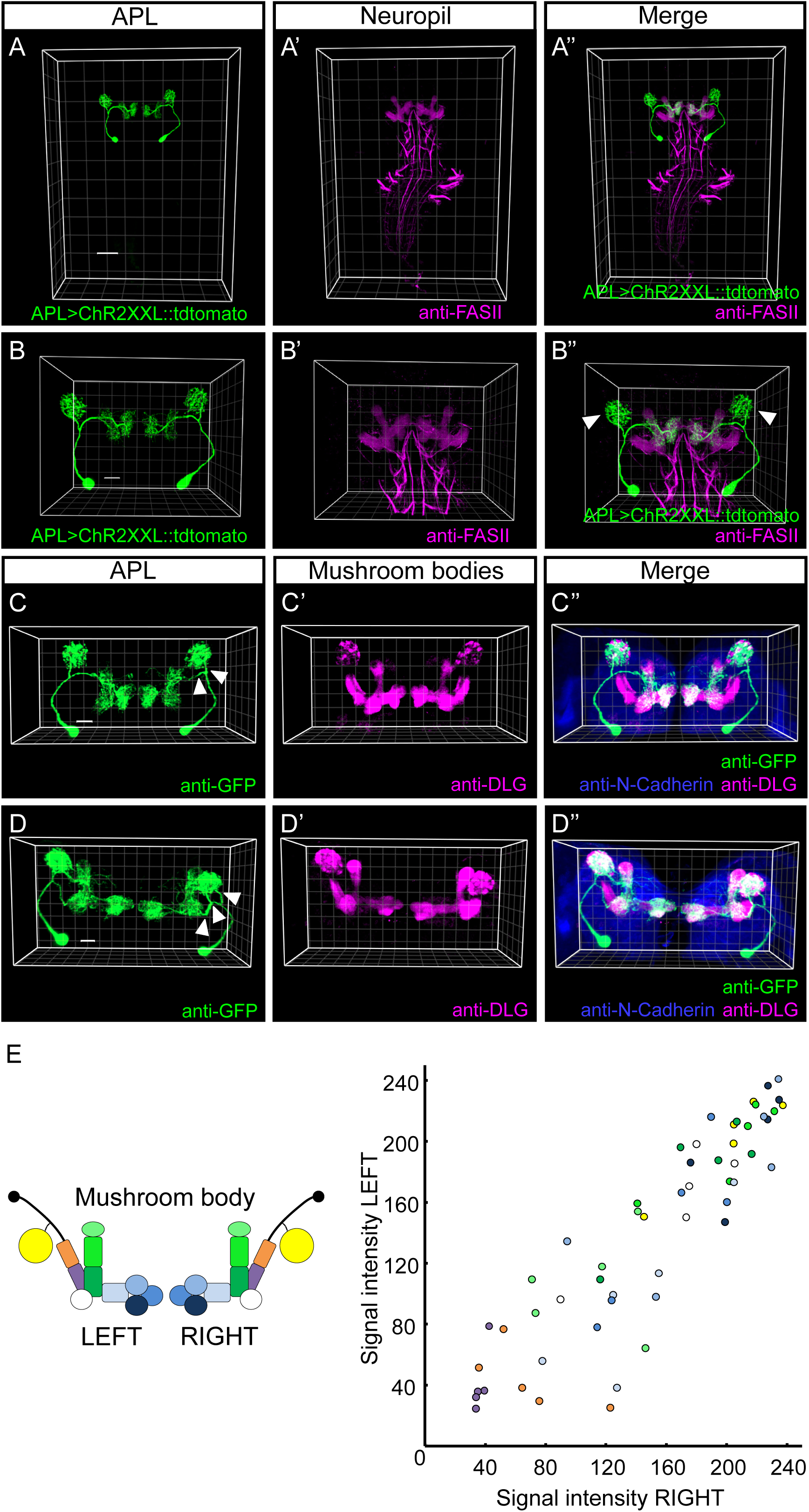
Brain expression of the APL-GAL4 driver is restricted to the APL neuron. **(A-A’’)** 3D view of the expression pattern from the APL-GAL4 driver in a third-instar larval brain visualized using the fluorescence signal from the UAS-ChR2XXL::tdtomato effector (APL>ChR2XXL::tdtomato; green). Axon-rich regions of the mushroom body peduncle and lobes can be discerned as references after labeling with a primary monoclonal mouse anti- FASII antibody and a secondary polyclonal goat anti-mouse Alexa Fluor 488 antibody (anti- FASII; magenta). Transgene expression is specific to the hemispherically unique APL neuron. The data were acquired with a 20x glycerol objective; scale bar and grid spacing: 50 µm. **(B- B’’)** As in (A-A’’), providing a close-up view of the mushroom bodies. APL sends projections into the calyx and a subset of the compartments of the medial and vertical lobes. White arrowheads in (B’’) point to the calyx, which is innervated by APL but is largely devoid of the axonal FASII marker. The data were acquired with a 63x glycerol objective; scale bar and grid spacing: 20 µm. **(C-D’’)** As in (B-B’’), except that the APL-GAL4 driver was crossed to UAS- mCD8::GFP as the effector. APL membranes can be visualized after labeling with a primary polyclonal rabbit anti-GFP antibody and a secondary polyclonal goat anti-rabbit Alexa Fluor 488 antibody (anti-GFP; green). The mushroom bodies are labeled by a primary monoclonal mouse anti-DLG antibody and a secondary polyclonal goat anti-mouse Alexa Fluor 568 antibody (anti-DLG; magenta); neuropils can be discerned as a reference by a primary monoclonal rat anti-N-Cadherin antibody and a secondary polyclonal goat anti-rat Alexa Fluor 647 antibody (anti-N-Cadherin; blue). Close-up analysis of the APL morphology revealed two, or in one case three, branches (white arrowheads in C-D, respectively) splitting from the primary neurite; notably, these numbers of branches do not differ between the two hemispheres (N = 11 brains). The data were acquired with a 16x glycerol objective; scale bar and grid spacing: 20 µm. **(E)** For each mushroom body compartment, the mean pixel intensities of APL labeling in the right hemisphere versus the left hemisphere are plotted (compartmental color code in accordance with the mushroom body schematic). The observed correlation indicates no inter-hemispheric difference in APL morphology (Pearson correlation (r)= 0.9747; p< 0.05). The sample size (number of brains) is given within the figure. The source data and results of all statistical tests are documented in Extended Data Figure 3-1.

To validate the expression of the ChR2XXL effector protein used for activating APL, the APL-GAL4 driver was crossed to the UAS-ChR2XXL effector (Dawydow et al., 2014; Bloomington Stock Center no. 58374). Following the procedure of Schleyer et al. (2020), brains of third-instar larval progeny (abbreviated as APL>ChR2XXL) were dissected in ice-cold Ca^2+^- free saline solution and fixed in Bouin’s solution (HT10132, Sigma-Aldrich) for 7 min at room temperature. After six successive washing steps (3 x brief; 3 x 15 min) in 0.2% PBT, the brains were incubated overnight at 4°C with a primary monoclonal mouse anti-ChR2 antibody (610180, ProGen Biotechnik) diluted 1:100 in 0.2% PBT. The brains were then washed (3 x 10 min each) in 0.2% PBT and incubated for 1 h at room temperature with a secondary polyclonal donkey anti-mouse Cy3 antibody (715-165-150, Jackson ImmunoResearch Laboratories) diluted 1:300 in 0.2% PBT. Finally, the samples were washed (3 x 10 min each) in 0.2% PBT and mounted in Vectashield (Vector Laboratories Inc) on a cover slip. Image z-stacks were acquired with a Leica TCS SP8 confocal microscope (Leica Mikrosysteme Vertrieb GmbH) at a format of 1024 × 1024. Image processing was performed using Imaris software (version 9.72, Bitplane).

#### GABA staining

To confirm the presence of GABA in APL, the APL-GAL4 driver was crossed to a UAS-CsChrimson::mVenus effector (Klapoetke et al., 2014; Bloomington Stock Center no. 55135), and third-instar progeny (abbreviated as APL>Chrimson) were dissected in PBS. Signal detection from the mVenus tag of the Chrimson transgene allows visualization of APL membranes without antibodies under the fluorescence microscope. The brains were fixed for 20 min with 4% PFA in 3% PBT on ice. After successive washing steps (2 x brief; 1 x 5 min; 3 x 15 min; 1 x 2 h) in 3% PBT, the brains were blocked for 1-2 h in 2% NGS solution (S-1000, Vector Laboratories Inc; in PBS) on ice. After two overnight incubations at 4°C with the primary antibodies, the brains were rinsed (2 x brief; 1 x 5 min; 3 x 15 min; 1 x 2 h) in 3% PBT and incubated overnight with the secondary antibodies at 4°C. The preparations were finally washed (2 x brief; 1 x 5 min; 5 x 15 min) in 3% PBT, mounted in Vectashield (Vector Laboratories Inc) on a cover slip, and scanned under a LSM510 confocal microscope (Zeiss) at a format of 1024 × 1024. Image processing was performed using Imaris software (version 9.72, Bitplane).

The primary antibody mixture consisted of (i) 2% NGS diluted 1:25 in 3% PBT, (ii) a monoclonal rat anti-N-Cadherin antibody (DN-Ex #8-s, Developmental Studies Hybridoma Bank) diluted 1:50 in 3% PBT (for neuropil staining), and (iii) a polyclonal rabbit anti-GABA antibody (A2052, Sigma Aldrich) diluted 1:500 in 3% PBT.

The secondary antibody mixture consisted of (i) 2% NGS diluted 1:25 in 3% PBT, (ii) a polyclonal Cy3-conjugated goat anti-rat antibody (A10522, Life Technologies) diluted 1:200 in 3% PBT, and (iii) a polyclonal Cy5-conjugated goat anti-rabbit antibody (A10523, Life Technologies) diluted 1:200 in 3% PBT.

#### APL regional synaptic polarity

To analyze the regional synaptic polarity of the larval APL neuron, the APL-GAL4 driver was crossed to a double effector with both UAS-Dsyd-1::GFP (Owald et al., 2015) and UAS-DenMark (Nicolai et al., 2010; Bloomington Stock Center no. 33062). Third-instar progeny (abbreviated as APL>Dsyd-1::GFP/DenMark) were dissected, fixed, dehydrated, and mounted as described in the preceding section. Image processing was performed and the corresponding Movie 3 was generated using Imaris software (version 9.72, Bitplane).

The primary antibody mixture consisted of (i) 2% NGS diluted 1:25 in 3% PBT, (ii) a monoclonal rat anti-N-Cadherin antibody (DN-Ex #8-s, Developmental Studies Hybridoma Bank) diluted 1:50 in 3% PBT (for neuropil staining), (iii) a polyclonal FITC-conjugated goat anti-GFP antibody (ab 6662, Abcam) diluted 1:1000 in 3% PBT (for visualization of the GFP-tag from Dsyd-1::GFP to label pre-synaptic regions), and iv) a polyclonal rabbit anti-DsRed antibody (632496, Clontech) diluted 1:200 in 3% PBT (for detecting the DenMark signal to label post-synaptic regions).

The secondary antibody mixture consisted of (i) 2% NGS diluted 1:25 in 3% PBT, (ii) a polyclonal Cy3-conjugated goat anti-rat antibody (A10522, Life Technologies) diluted 1:200 in 3% PBT, and (iii) a polyclonal Cy5-conjugated goat anti-rabbit antibody (A10523, Life Technologies) diluted 1:200 in 3% PBT.

### Chemical tagging for tracking APL development

Chemical tagging provides an alternative method to immunohistochemistry for labeling specific cells and structures in tissues. The tag-based approach uses genetically driven, enzyme-based protein “tags” that are expressed in specific cells and that covalently bind small fluorescent substrates, resulting in fast and specific tissue staining with low background signals (Kohl et al., 2014; Sutcliffe et al., 2017; Meissner et al., 2018).

We used such tagging to track the regional synaptic polarity of APL during development. Specifically, we used the synaptic reporters synaptotagmin fused to the chemical tag SNAPm (Syt1-SNAPm) to label pre-synaptic regions, and telencephalin fused to CLIPm (TLN-CLIPm) to label post-synaptic regions (Kohl et al., 2014). The effectors UAS-Syt1:SNAP (Kohl et al., 2014; Bloomington Stock Center no. 58379), UAS-TLN:CLIP (Kohl et al., 2014; Bloomington Stock Center no. 58382) and UAS-mCD8::GFP (for labeling APL; Lin et al., 2014) were used together with the intersectional driver APLi-GAL4 (NP2631-GAL4, GH146-FLP, tubP-FRT- GAL80-FRT) for specific expression in both larval and adult APL neurons (Lin et al., 2014; Mayseless et al., 2018) – since APL-GAL4 does not cover the adult APL neurons (not shown).

Our procedures followed Kohl et al. (2014). In brief, brains of third-instar larvae, pupae (6 h or 12 h after puparium formation), and adults of the genotype APLi/Syt1:SNAP> mCD8::GFP/TLN:CLIP were dissected in ice-cold phosphate buffer (PB; 0.1 M) and fixed in 4% PFA at room temperature for 20 min. The brains were permeabilized and washed (3 x 10 min) in PBT (0.3% Triton-X 100 in PBS). Chemical tag ligands were then applied in a 300 μL volume on a nutator for 15 min, at room temperature. The chemical substrates were SNAP-tag ligands (SNAP surface 549 - BG 549 [NEB, S9112S]) and CLIP-tag ligands (CLIP surface 647 - BC 647 [NEB, S9234S]) at final concentrations of 1 μM in 0.3% PBT. To minimize cross-reactivity, the SNAP-tag ligands were applied 10 min before the CLIP-tag ligands. To label APL, the brains were immunostained: three consecutive washing steps (10 min each) in 0.3% PBT were followed by 30 min incubation in blocking solution with 5% NGS (005-000-121, Jackson Immunoresearch Laboratories) in PBT. The brains were then incubated overnight with a primary antibody mixture consisting of (i) 5% NGS diluted in 0.3% PBT and (ii) a polyclonal chicken anti-GFP antibody (AB_10000240, Aves Labs) diluted 1:500. After five consecutive washing steps (3 x brief; 2 x 20 min) in 0.3% PBT, the brains were incubated for 2 h at room temperature with a polyclonal secondary FITC-conjugated goat anti-chicken antibody (A16055, Invitrogen) diluted 1:300 in 0.3% PBT. The preparations were mounted on slides in SlowFade (Invitrogen), and examined under a confocal microscope (Zeiss LSM 800). Image processing was performed using Imaris software (version 9.2 Bitplane).

### Volume reconstruction of APL from an EM dataset

We added radial volume annotations to an existing skeleton reconstruction of the APL neuron in both hemispheres (Eichler et al., 2017) from an electron microscopy dataset of a 6 h-old stage 1 larva (Ohyama et al., 2015). More details of the neuron reconstructions can be found in Eichler et al. (2017). Volume annotations were made manually using the web-based software CATMAID (Saalfeld et al., 2009; Schneider-Mizell et al., 2016), which was extended with a tool to allow for rapid graphical annotations of the radii of contiguous cable segments with similar radius. Radial annotations were used to create a volumetric representation of the cells’ morphology as conical frustum compartments. The radii were placed so as to preserve the approximate volume of the irregularly shaped processes while accounting for the anisotropic image resolution of 3.8 nm × 3.8 nm × 50 nm. We defined the axon and dendrite of both APL neurons as the two synapse-rich areas along the arbor separated from the neurite by the high Strahler branch point nearest to the cell body. Reconstructed neurons and their synapses were analyzed using the natverse package (http://natverse.org/) (Bates et al., 2020) in R (version 3.6.2) and plotted using Blender (version 2.79) with the CATMAID-to-Blender plugin (https://github.com/schlegelp/CATMAID-to-Blender; Schlegel et al., 2016).

### Dendrogram representations of APL synapses and branching

Neuron dendrograms are simplified, but topologically correct, two-dimensional representations of neurons with complex morphologies (Strauch et al., 2018). As relative branch lengths and synapse location are preserved, dendrograms can be employed to visualize the spatial distribution of synapses in an easily readable way. The APL dendrograms are derived from existing electron microscopy reconstructions (Eichler et al., 2017) and were created following established computational methods (Strauch et al., 2018). Additionally, mushroom body compartment boundaries were superimposed onto the dendrograms, based on the projection patterns of mushroom body extrinsic neurons (Saumweber et al., 2018). We used the natverse toolbox for R (Bates et al., 2020) and custom code (A. Bates, University of Cambridge) to extract the synaptic coordinates from the CATMAID L1 dataset and to generate an envelope that surrounds the synapses formed by the mushroom body extrinsic neurons belonging to each given compartment. Compartment boundaries were plotted onto the dendrograms as hulls around the synapses located within the respective compartment. Compartments that extend across several dendrogram branches are connected by dashed lines.

To analyze the relative distribution of APL-to-KC and KC-to-APL synapses in the calyx, we followed a procedure described in detail in Schleyer et al. (2020). In brief, we computed geodesic distances between synapses, i.e. "cable length" distances along the neuron’s branches. Based on the geodesic distances, a clustering algorithm served to partition all synapses, regardless of their type, into local synapse clusters, i.e. regions of high synapse density (domains). For each domain, the distances were then evaluated from the APL-to-KC (or KC-to-APL) synapses to the cluster’s centroid point, which served as a measure for the spatial distribution of APL-to-KC (or KC-to-APL) within the domain.

### Functional imaging

The functional imaging methods follow those described in greater detail previously (Selcho et al., 2017; Lyutova et al., 2019). In brief, to monitor intracellular Ca^2+^ levels of KCs in response to optogenetic activation of the APL neuron, the lexA-lexAop system was used to express the fluorescent Ca^2+^ reporter GCaMP6m in KCs (effector lexAOp-GCaMP6m (Chen et al., 2013; Bloomington Stock Center no. 44276); driver R14H06-lexA (Pfeiffer et al., 2013; Bloomington Stock Center no. 52482)), and the GAL4-UAS system was used to express ChR2-XXL in APL (effector UAS-ChR2XXL (Dawydow et al., 2014; Bloomington Stock Center no. 58374); driver R26G02-GAL4 (Saumweber et al., 2018; Bloomington Stock Center no. 48065)). Note that the R26G02-GAL4 driver covers APL plus additional cells outside the mushroom body; it was chosen because the more specific APL-GAL4 driver, as a split-GAL4 strain, could not be combined with the lexA-lexAop system. Using classical genetics, we generated strains that carried a combination of the drivers (R14H06-lexA/R26G02-GAL4) or the effectors (lexAop- GCaMP6m/UAS-ChR2XXL; Lyutova et al., 2019). These strains were crossed, and from the progeny (abbreviated as APL_26G02_>ChR2XXL; KC>GCaMP6m) larval brains were dissected and placed in a Petri dish containing 405 µl hemolymph-like HL3.1 Ringer solution. Images with regions of interest (ROI) around the calyx and KCs were recorded with an Axio Examiner D1 microscope (Zeiss) using a W Plan-Apochromat 20 x 1.0 DIC (UV) VIS-IR objective and a Axiocam 506 camera (Zeiss). We monitored fluorescence intensity upon pulsed 475-nm light (Colibri LED, Zeiss) at an intensity of 1.8 mW/cm^2^ for a first observation period, followed by pulses at an intensity of 4.1 mW/cm^2^ for a second observation period immediately thereafter. The light pulses were of 80 ms duration and were given at a 2-s onset-onset interval. We used larvae without the APL driver as the genetic control (abbreviated as +>ChR2XXL; KC>GCaMP6m).

To see whether APL activation would be potent enough to reduce the very high intracellular Ca^2+^ levels that result from cholinergic stimulation of the KCs, we used carbamylcholine (CAS: 51-83-2, Sigma Aldrich). Specifically, we monitored fluorescence intensity under pulsed 475- nm light as described above for a total of 8 min and after the first 2 min manually bath-applied 45 µl carbamylcholine, dissolved in HL3.1 to a final concentration of 10^-4^ M.

We present the fluorescence intensity (ΔF/F_0_) across the observation period normalized to the values at its beginning (Normalized ΔF/F_0_) as the baseline. To analyze the differences between the experimental conditions, we determined, for each calycal ROI/ KC, the maximum difference from the baseline.

### Behavioral assays

#### Experimental setup

Larvae were trained and tested on Petri dishes (9 cm inner diameter; Sarstedt) filled either with 1% agarose only (CAS: 9012-36-6, Roth) or with 1% agarose containing 2 mol/L D-fructose (99% purity; CAS: 57-48-7, Roth) as the taste reward (+). Once the contents had solidified, the dishes were covered with their lids and left at 4°C until the experiment started, and for a maximum of 2 weeks.

As the odors, *n*-amyl acetate (AM; CAS: 628-63-7, Merck) diluted 1:20 in paraffin oil (CAS: 042-47-5, AppliChem) and 1-octanol (OCT, undiluted; CAS: 111-87-5, Sigma Aldrich) were used. The paraffin oil is without behavioral effect as an odor (Saumweber et al., 2011). Before the experiments, 10 µL of the respective odor was added to odor containers (5 mm inner diameter) covered by perforated lids (5-10 holes of 0.5 mm diameter each). Larvae were collected from their food vial with a brush, briefly rinsed in tap water, and used immediately for behavioral experiments.

Behavioral assays were carried out in a light-shielded, custom-built box, as described in Schleyer et al. (2020). In brief, the box contained a 24 x 12 LED array light table (Solarox) with a 6-mm-thick Plexiglas diffusion panel placed above it, providing constant light conditions and intensity for the activation of light-gated ion channels expressed in neurons of interest (see section “*Genotypes and methods for optophysiology*”). Larvae in Petri dishes were placed onto the diffusion panel and were surrounded by a translucent polyethylene ring. The ring featured 30 infrared LEDs mounted behind to deliver light (invisible to the animals) allowing behavioral recording for offline tracking analysis (see section “*Video recording and tracking of locomotion*”).

#### Odor-fructose reward association

A two-group, reciprocal conditioning paradigm was used following standard procedures (Scherer et al., 2003; Neuser et al., 2005; Saumweber et al., 2011; for a detailed manual: Michels et al., 2017). In brief, one group of larvae received the odor presented together with the fructose reward (paired training), whereas a second group received separate presentations of the odor alone and the fructose reward alone (unpaired training).

For paired training, a cohort of ∼ 30 larvae was placed at the center of a Petri dish filled with agarose that was mixed with fructose as the reward (+). Two containers were filled with the odor (AM+) and placed on opposite sides of the Petri dish. The lid was then closed, and the larvae were allowed to move freely for 2.5 min. The larvae were then removed and placed on a fresh, pure-agarose Petri dish in the presence of two empty containers (EM), the lid was closed, and the larvae could again move freely for 2.5 min. This training cycle was performed once only, unless mentioned otherwise. The sequence of training was alternated across replications; i.e. for half of the cases we started with AM as described above (AM+/EM), and for the other half with EM (EM/AM+). After training, the larvae were tested for their odor preference. Specifically, the animals were transferred to the center of a fresh, pure-agarose Petri dish (i.e. without fructose reward, unless mentioned otherwise) featuring one AM container on one side, and one EM container on the other side. After 3 min, the number of larvae on the AM side (#AM), the EM side (#EM), as well as on the middle “neutral” stripe (10 mm), was counted, and the olfactory preference score (PREF) was calculated as:

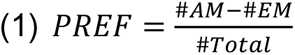

Thus, preference scores may range from 1 to -1, with positive values showing preference for AM, and negative values indicating avoidance of AM. Larvae crawling up onto the lid or onto the odorant containers during the test (< 5%) were discarded from the analysis.

For unpaired training, the procedure was the same except that the odor and the reward were presented separately to the animals. That is, after collection the larvae were placed on a fresh, pure-agarose Petri dish in the presence of two containers both filled with AM. Then, the larvae were transferred onto a fresh agarose Petri dish with fructose added, in the presence of two empty containers (EM+). Again, the training sequence started with AM (AM/EM+) in half of the cases, and in the other half of the cases with EM (EM+/AM). The larvae were then tested for their AM preference, and the olfactory preference score was calculated as for the paired group (equation 1).

Associative memory is indicated by a difference in preference for AM after paired training compared to the reciprocal, unpaired training. These differences in AM preference were quantified by the associative memory score:

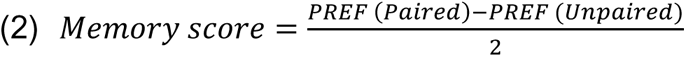

Thus, memory scores may range from 1 to -1, with positive values indicating appetitive associative memory, and negative values indicating aversive associative memory. These experiments were combined with optogenetic APL activation (see section “*Genotypes and methods for optophysiology*”), as mentioned in the Results section.

#### Odor-APL association

Based on early results in this study (see Results section), we suspected that optogenetic activation of the APL neuron might have a rewarding effect. Therefore, the associative learning paradigm described above was modified by using optogenetic APL activation (+) instead of a fructose reward (i.e. no real reward was presented). In the paired group, AM was presented together with continuous 2.5-min light stimulation to activate APL, whereas empty containers were subsequently presented in darkness, also for 2.5 min (AM+/EM). In the unpaired group, the larvae were exposed to odor and light separately (AM/EM+). This training cycle was performed once only, with the training sequence alternated across repetitions as described in the preceding section. After training, the larvae were tested on a fresh, pure-agarose Petri dish, and their odor preference as well as the memory score were calculated as detailed above (equation 1, equation 2).

In addition, a differential two-odor version of the paradigm using APL activation as the reinforcer was used. This was performed as described above except that instead of using empty containers (EM), the containers were filled with 1-octanol (OCT, undiluted). Thus, differential conditioning followed the logical structure of training being either AM+/OCT or in the reciprocal case AM/OCT+ (again, the training sequence was alternated across repetitions of the experiments). The larvae were then tested for their choice between AM and OCT on a fresh, pure-agarose Petri dish, and the data were analyzed, with due adjustment, as detailed above (equation 1, equation 2). In this case, positive memory score values thus indicate odor-specific appetitive associative memory, whereas negative memory score values indicate odor-specific aversive associative memory.

Whenever variations on the above paradigms were used, these are mentioned in the Results section.

#### Innate olfactory behavior

The odor preference of experimentally naïve larvae was assayed following standard procedures (Saumweber et al., 2011). Cohorts of ∼ 30 animals were transferred onto a pure- agarose plate in the presence of one odor-filled container and another empty container placed on opposite sides of the plate. Odor preference was calculated after 3 min following equation (1). To probe whether APL activation has an effect on innate olfactory behavior, the test was carried out either without light stimulation, or with light stimulation.

#### Genotypes and methods for optophysiology

For the experiments on APL activation, we used third-instar transgenic larvae expressing either ChR2XXL or Chrimson in APL. To this end, APL-GAL4 was crossed to UAS-ChR2XXL or to UAS-CsChrimson::mVenus as the effector. Double-heterozygous progeny (abbreviated as APL>ChR2XXL or APL>Chrimson) were used for activation of the APL neuron; larvae heterozygous for either the GAL4 element (APL>+) or the UAS element (+>ChR2XXL or +>Chrimson) were used as the driver and the effector genetic control, respectively. To obtain the driver controls, APL-GAL4 was crossed to w^1118^ (Bloomington Stock Center no. 3605, 5905, 6326). As regards the effector controls, a strain lacking the GAL4 domains but containing the two split-GAL4 landing sites (attP40/attP2) was crossed to UAS-ChR2XXL or UAS-CsChrimson::mVenus. For experiments using Chrimson, the flies were raised on food supplemented with all-trans retinal (100 mM final concentration; cat: R2500; CAS: 116-31-4, Sigma Aldrich), unless mentioned otherwise.

For the experiments on MBON activation, UAS-ChR2XXL was crossed to one of the three following drivers: (i) R36G04-GAL4, covering the two calyx MBONs in each hemisphere, plus additional cells in the ventral nerve cord (Saumweber et al., 2018; Bloomington Stock Center no. 49940; abbreviated as MBONa1,a2-GAL4); (ii) the split-GAL4 line SS02006, covering only one calyx MBON in each hemisphere (Eschbach et al., 2021; kindly provided by M. Zlatic, University of Cambridge; abbreviated as MBONa1-GAL4); (iii) SS01417, covering one, or in some cases both, of the calyx MBONs in each hemisphere (Extended Data Figure 16-1; Eschbach et al., 2021; kindly provided by M. Zlatic, University of Cambridge; abbreviated as MBONa2-GAL4). Again, double-heterozygous progeny (abbreviated as MBONa1,a2>ChR2XXL, MBONa1>ChR2XXL or MBONa2>ChR2XXL) were used for activating the calyx MBONs; driver and effector control larvae were obtained as detailed in the preceding paragraph.

For simultaneous activation of APL and ablation of the pPAM neurons, the lexA-lexAop system was used to express the pro-apoptotic *reaper* gene in the pPAM neurons (effector lexAop-reaper (Herranz et al., 2014); driver R58E02-lexA (Lyutova et al., 2019; Bloomington Stock Center no. 52740)), and the GAL4-UAS system was used to express ChR2-XXL in APL (effector UAS-ChR2XXL (Dawydow et al., 2014; Bloomington Stock Center no. 58374); driver R55D08-GAL4 (Saumweber et al., 2018; Bloomington Stock Center no. 39115)). Note that the R55D08-GAL4 driver covers APL plus additional cells outside the mushroom body; it was chosen because the more specific APL-GAL4 driver, as a split-GAL4 strain, could not be combined with the lexA-lexAop system. Using classical genetics, we generated strains that carried a combination of the drivers (R58E02-lexA/R55D08-GAL4) or the effectors (lexAop-reaper/UAS-ChR2XXL; Lyutova et al., 2019). These strains were crossed, and the progeny obtained (abbreviated as APL_55D08_>ChR2XXL; pPAM>reaper) were used for the experiment. As regards single effector controls, a strain lacking lexAop-reaper but containing UAS- ChR2XXL was crossed to R58E02-lexA/R55D08-GAL4 (abbreviated as APL_55D08_>ChR2XXL; pPAM>+). As regards single driver controls, a strain lacking R58E02-lexA but containing R55D08-GAL4 was crossed to lexAop-reaper/UAS-ChR2XXL (abbreviated as APL_55D08_>ChR2XXL; +>reaper).

The above-mentioned custom-built box (see section “Experimental setup”) was equipped for illumination from a blue LED light table when ChR2XXL was used (wavelength: 470 nm; intensity: 120 μW/cm^2^; Solarox), or from a red LED light table when Chrimson was used (wavelength: 630 nm; intensity: 350 µW/cm^2^; Solarox).

For silencing experiments, the light-gated chloride channel GtACR1 was used. Specifically, either APL-GAL4, R36G04-GAL4, SS02006-GAL4 or SS01417-GAL4 were crossed to UAS- GtACR1::YFP (König et al., 2019; Bloomington Stock Center no. 9736; kindly provided by R. Kittel, University of Leipzig). Double-heterozygous progeny (APL>GtACR1, MBONa1,a2>GtACR1, MBONa1>GtACR1 or MBONa2>GtACR1) were used for silencing the respective neurons; driver and effector control larvae were obtained as described in the preceding paragraphs. A green LED light table (wavelength: 520 nm; intensity: 2003 μW/cm^2^; Solarox) was used for illumination.

In all cases, the timing of illumination is mentioned for each experiment in the Results section. As all effectors are sensitive to daylight, the breeding of all transgenic animals was performed in darkness, effectuated by black covers wrapped around the food vials. All behavioral experiments were carried out in parallel for the respective experimental group and genetic controls; investigators were blind with respect to genotypes.

#### Video recording and tracking of locomotion

For a subset of experiments, larval behavior was videorecorded throughout the test and analyzed as described by Paisios et al. (2017). In brief, four behavioral features were analyzed in relation to odor:

> First, the olfactory preference (PREF time, in s) was calculated as:
>
>
> 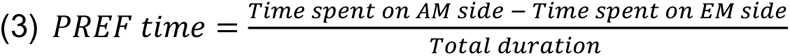

Thus, preference scores may range from 1 to -1, with positive scores indicating that larvae spent more time on the odor side, and negative values indicating more time spent on the non-odor side, representing approach and avoidance, respectively.

> Second, the head cast (HC) rate modulation was calculated as:
>
>
> 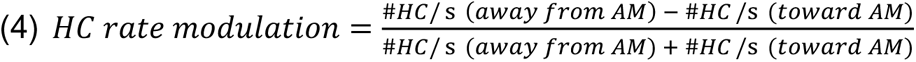

Thus, positive scores indicate odor approach; i.e. the larvae make more HCs when crawling away from the odor than when crawling towards it. Conversely, negative scores indicate odor avoidance.

> Third, the HC reorientation (°) was calculated as:
>
>
> 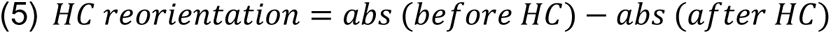

The absolute heading angle (abs) indicates how the larva’s head is oriented relative to the odor. For instance, at abs 180° or 0° the odor is located behind or in front of the animal, respectively. Thus, positive values indicate odor approach; i.e. the head cast directs the larva towards the odor instead of away from it. Conversely, negative values indicate odor avoidance.

> Fourth, the run speed modulation was calculated as:
>
>
> 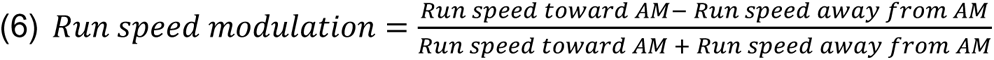

Thus, positive values for run speed modulation indicate that animals slow down whenever they head away from the odor, and speed up whenever they move towards it, indicating approach. Conversely, negative values indicate avoidance.

### Pharmacological manipulation of dopamine synthesis

To test whether the dopaminergic system is implicated in odor-APL associative learning, a systemic pharmacological approach was used to disrupt dopamine synthesis (Neckameyer, 1996; Kaun et al., 2011; Thoener et al., 2020). This approach was combined with behavioral experiments using optogenetic APL activation as the reinforcer (see section “*Odor-APL association”*) and followed the procedures described in Thoener et al. (2020). In brief, a 0.5 mg/ml yeast solution was produced and kept for up to one week at 4 °C. The dopamine-synthesis inhibitor 3-Iodo-L-tyrosine (3IY; CAS: 70-78-0, Sigma Aldrich; concentration: 5 mg/ml) was added to samples of 2 ml yeast solution. In the instances mentioned in the Results section, the dopamine precursor 3,4-dihydroxyphenylalanine (L-DOPA; CAS: 59-92-7, Sigma Aldrich; concentration: 10 mg/ml) was added to a yeast solution with or without 3IY. After mixing on a shaker for 1 h, the solutions were transferred into vials containing two pieces of PET mesh. Third-instar progeny of the APL-GAL4 driver crossed to UAS-ChR2XXL (APL>ChR2XXL) were transferred from their food vials to the respective yeast solutions. After a feeding period of 4 h at 25°C and 60-70% relative humidity, the larvae were briefly washed in water and immediately used in behavioral experiments.

### Experimental design and statistical analyses

The source data and the results of all statistical tests, performed in Statistica 13 unless mentioned otherwise (SCR_014213, StatSoft Inc, Tulsa), are included in Extended Data Figure 3-1. Graphs, figures, and sketches were generated with Statistica 13, Corel Draw 2019 (SCR_013674, Corel Corporation), and GraphPad Prism 6 (SCR_002798, GraphPad Software Inc); references are documented in Extended Data Table 2-1.

To compare the compartmental coverage between the APL neuron of each hemisphere (see Figure 3E), a Pearson correlation was performed; the data are displayed as a scatter plot.

To compare the radii of the neurite, dendrite, and axon of APL in both hemispheres (see Figure 6D), Kruskal-Wallis tests (KW) and Mann-Whitney-Wilcoxon (MWW) tests (total of three MWW tests per APL) were used for multiple and two-group comparisons, respectively (R Core Team, 2016). The Bonferroni-Holm correction was applied to maintain an error rate below 5% (Holm, 1979). The data are displayed as violin plots, the bars showing the mean.

The experiments in Figure 5 followed a two-group design with two genotypes (APL_26G02_>ChR2XXL; KC>GCaMP6m as the experimental genotype and +>ChR2XXL; KC>GCaMP6m as the genetic control). In Figure 5A, C and E, the data for each calycal ROI/ KC are plotted over time (showing mean +/- SEM). In Figure 5B, D and F, the data are plotted as the maximum difference from the baseline for each calycal ROI/ KC (showing mean and data-point scatter); MWW tests were performed between genotypes.

For the experiments in Figures 7, 9-18 and Extended Data Figure 7-4, the data are displayed as box plots, the middle line showing the median, the box boundaries the 25 and 75% quantiles, and the whiskers the 10 and 90% quantiles; outliers are not displayed.

**Figure 4.**
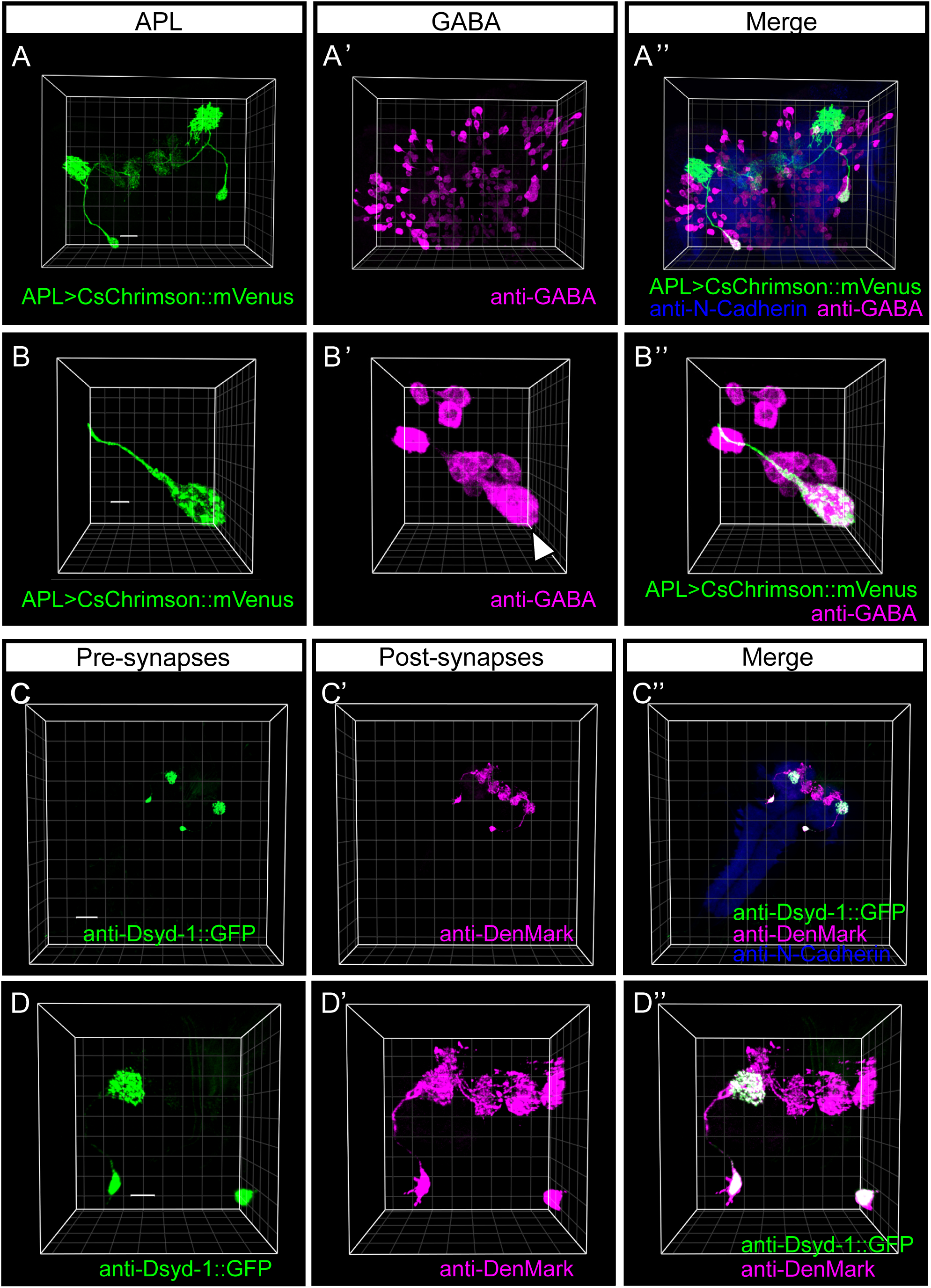
The larval APL neuron is GABAergic and is pre-synaptic in the calyx and post- synaptic in both the calyx and the lobes. (**A-A’’**) 3D view of the expression pattern from the APL-GAL4 driver in the third-instar larval brain visualized using the fluorescence signal from the Chrimson effector (APL>CsChrimson::mVenus; green). GABAergic signals can be visualized after labeling with a polyclonal rabbit anti-GABA antibody and a polyclonal Cy5-conjugated goat anti-rabbit antibody (anti-GABA; magenta). The white arrowheads in (A’’) point to an overlap of the GABA signal and the fluorescence signal in the APL soma. Neuropil regions are visualized as a reference by using a primary monoclonal rat anti-N-Cadherin antibody and a secondary polyclonal goat anti-rat Cy3 antibody (anti-N-Cadherin; blue). The data were acquired with a 63x glycerol objective; scale bar and grid spacing: 20 µm. (**B-B**’’) As in (A-A’’), providing a close-up view of the APL soma. The white arrowhead in (B’) points to the APL soma surrounded by additional GABAergic cells. The data were acquired with a 63x glycerol objective; scale bar and grid spacing: 5 µm. (**C-C**’’) The APL-GAL4 driver was crossed to a double effector with both UAS-Dsyd-1::GFP and UAS-DenMark to label the pre- and post- synaptic sites of the APL neuron in third-instar larvae. Pre-synaptic regions of APL can be visualized after labeling with a polyclonal FITC-conjugated goat anti-GFP antibody (anti-Dsyd-1::GFP; green). Post-synaptic regions are revealed after labeling with a primary polyclonal rabbit anti-DsRed antibody and a secondary polyclonal goat anti-rabbit Cy5 antibody (anti- DenMark; magenta). Neuropil regions are visualized as a reference by using a primary monoclonal rat anti-N-Cadherin antibody and a secondary polyclonal goat anti-rat Cy3 antibody (anti-N-Cadherin; blue). The pre-synaptic marker Dsyd-1 is mainly restricted to the calyx, whereas the post-synaptic marker DenMark localizes to both the calyx and a subset of the compartments in the lobes, confirming the regional synaptic polarities of the larval APL neuron (Masuda-Nakagawa et al., 2014; Eichler et al., 2017). The data were acquired with a 16x glycerol objective; scale bar and grid spacing: 50 μm. (**D-D**’’) As in (C-C’’), providing a close-up view of the pre- and post-synaptic regions of APL; scale bar and grid spacing: 25 μm. For a corresponding movie see Movie 3.

To compare the geodesic distances of synapses to the center of their respective center-surround structure on the left-hemisphere APL neuron (see Figure 7D), we performed a MWW test between KC-to-APL and APL-to-KC synapses. Corresponding analyses for the right-hemisphere APL neuron can be found in Extended Data Figure 7-4.

For the behavioral results displayed in Figures 9-18, Kruskal-Wallis tests (KW) and MWW tests were used for multiple and two-group comparisons, respectively. For comparisons to chance levels (i.e. to zero), one-sample sign tests (OSS; corresponding to binom.test in R version 3.3.2, R Core Team, 2016) were used. The Bonferroni-Holm correction was applied to maintain an error rate below 5%. Sample sizes (biological replications) were chosen based on previous studies that had revealed moderate to mild effect sizes (Paisios et al., 2017; Saumweber et al., 2018) and are indicated in the figure legends. A sample size of N = 1 included ∼ 30 animals of both sexes for each reciprocally trained group, and ∼ 30 animals of both sexes for all innate preference experiments. All behavioral experiments were performed in parallel for the respective experimental group and genetic controls; experimenters were blind to genotypes.

The experiments in Figure 9A-B, Figure 12A-B and E, Figure 15A, Figure 16A-B, and Figure 18 followed a three-group design with three genotypes (experimental genotype expressing ChR2XXL or GtACR1 or ChR2XXL/reaper, plus effector and driver controls). In case of significance, a KW test across all groups was followed by pairwise MWW tests between genotypes (three MWW tests in total). Comparisons to chance levels (i.e. to zero) were tested for each group by one-sample sign tests; in Figure 16A-B no one-sample sign tests were performed.

The experiments in Figures 9D, F-I and Figure 15D followed a two-group design with two test conditions (the presence or absence of light) for the experimental genotype expressing ChR2XXL (Figure 9D, F-I) or Chrimson (Figure 15D). MWW tests were performed between the two test conditions; no one-sample sign tests were performed.

The experiment in Figure 10 followed a six-group design with two test conditions (the presence or absence of the fructose reward) for the experimental genotype expressing ChR2XXL. The animals received paired or unpaired odor-fructose training, either not activating APL during training at all, or activating APL during odor presentation, or in the absence of odor. A KW test across all groups was followed by pairwise MWW tests between groups within the same test condition, as well as between test conditions for a given kind of training regimen (nine MWW tests in total). Differences from chance levels (i.e. from zero) were tested for in each group by one-sample sign tests.

The preference scores shown in Figure 11A-C underlie the associative memory scores in Figure 10. The experiment followed a four-group design with two test conditions (the presence or absence of the fructose reward) for the experimental genotype expressing ChR2XXL. A KW test across all groups was followed by pairwise MWW tests between groups that had received paired or unpaired odor-fructose training within the same test condition; in addition, MWW tests were performed between the respectively trained groups that were tested in the absence of fructose and the baseline odor preference scores (four MWW tests in total). Differences from chance levels (i.e. from zero) were tested for in each group by one-sample sign tests.

Figure 11D shows pooled preferences from Figure 11A-C and follows a three-group design for the experimental genotype expressing ChR2XXL. A KW test across all groups was followed by pairwise MWW tests between groups (three MWW tests in total); no one-sample sign tests were performed.

The experiment in Figure 12C followed a four-group design for the experimental genotype expressing ChR2XXL. After a KW test across all groups, MWW tests were performed between the group tested immediately after training (retention interval 0 min) and the groups tested 5, 10 or 20 min after training (three MWW tests in total). Differences from chance levels (i.e. from zero) were tested for in each group by one-sample sign tests.

The experiment in Figure 12D followed a nine-group design for the experimental genotype expressing ChR2XXL. A KW test was performed across all groups; no one-sample sign tests were performed.

Figure 13B describes the odor preferences of larvae of the experimental genotype expressing ChR2XXL tested after paired or unpaired training, over time. No statistical analyses were performed; rather, the data were collated over time and were statistically compared between the two training groups in Figure 13C (see next paragraph).

The experiments in Figure 13C-F, Figure 15B-C and Figure 17C followed a two-group design for the experimental genotype expressing Chrimson (Figure 15B-C) or ChR2XXL (Figures 13C-F, Figure 17C). MWW tests were performed between the two groups. Differences from chance levels (i.e. from zero) were tested for in each group by one-sample sign tests (in Figure 13C-F no one-sample sign tests were performed).

The experiments in Figure 14 followed a six-group design with three genotypes (experimental genotype expressing ChR2XXL, effector and driver controls) and two test conditions (the presence or absence of light). A KW test across all the groups was performed first. In case of significance, pairwise MWW tests were performed between genotypes within the same test condition, as well as between test conditions for a given genotype (nine MWW tests in total). In Figure 14A-B differences from chance levels (i.e. from zero) were tested for in each group by one-sample sign tests; in Figure 14C-E no one-sample sign tests were performed.

The experiments in Figure 16C-D followed a four-group design with two test conditions (the presence or absence of light) for the experimental genotype expressing GtACR1 (Figure 16C) or ChR2XXL (Figure 16D). A KW test was performed across all groups; no one-sample sign tests were performed.

The experiment in Figure 17D followed a four-group design for the experimental genotype expressing ChR2XXL. After a KW test across all groups, MWW tests were performed between the untreated group and (i) the group fed 3IY, (ii) the group fed 3IY plus L-DOPA, and (iii) the group fed L-DOPA; as well as between the group fed 3IY and the group fed 3IY plus L-DOPA (four MWW tests in total). Differences from chance levels (i.e. from zero) were tested for in each group by one-sample sign tests.

## Results

### Organization of the APL neuron

We first investigated the expression pattern of the APL-GAL4 driver in third-instar larvae in the full-body context. This took advantage of a combination of autofluorescence signals with fluorescence from UAS-mCherry-CAAX as the effector. These signals can be detected conveniently under a light-sheet microscope upon clearing the sample (Kobler et al., 2021). Fluorescence signals that can be observed across wavelengths and in both the experimental genotype (APL>mCherry-CAAX) and the genetic controls (+>mCherry-CAAX and APL>+ as the effector and the driver control, respectively) constitute autofluorescence and allow rich bodily detail to be discerned (Figure 2). We note that signals from fluorescent food particles contribute to individually-variable signals along the alimentary canal (Kobler et al., 2021). Fluorescence reflecting the expression of mCherry-CAAX, however, was reproducibly seen specifically in a giant pair of cells in the mushroom body region (Figure 2B**’’’**, **D’’’**, **F’’’**) that can be identified as APL (**Movie 1**). Thus, to the extent tested here and in the absence of a standard body against which full-body preparations can be systematically registered at high resolution, the effects observed using the APL-GAL4 driver can be interpreted without reference to transgene expression elsewhere in the body.

Combining the APL-GAL4 driver with the UAS-ChR2XXL::tdtomato effector and using the resulting fluorescence signal confirms that within the central nervous system and the mushroom body APL-GAL4 specifically covers APL (Figure 3A-B**’’**) (Saumweber et al., 2018). In addition, APL-GAL4 was crossed to UAS-mIFP/MB247>mCherry-CAAX (Kobler et al., 2021) to label APL together with the mushroom body (**Movie 2**). In both cases our results confirm that the primary neurite of APL splits to send projections separately into the calyx and the lobes of the mushroom bodies (Figure 3B**’’**, **Movie 2**) (Masuda-Nakagawa et al., 2014; Mayseless et al., 2018; Saumweber et al., 2018). This morphology was seen in 10 out of 11 preparations of third-instar larval brains with the APL-driver and UAS-mCD8::GFP as the effector (Figure 3C-C**’’**). Only in one preparation did the primary neurite split into three branches in both hemispheres (Figure 3D-D**’’**). This made us wonder whether there is any variability in the compartmental coverage of APL in the lobes, in particular in the third-instar larvae that we intended to use later in our behavioral analyses. Across five specimens of third-instar larval brains with the APL-driver and UAS-mCD8::GFP as the effector, coverage of the calyx and of the compartments was similar between the APL neuron of each hemisphere (Figure 3E). GFP signals were consistently strong in the calyx, close to absent in the two peduncle compartments, weak in the upper vertical lobe and the shaft of the medial lobe, and moderate to strong in the other compartments (Figure 3E). Taking the present data together with previously published data, we conclude that the larval APL innervates the calyx and six of the 10 compartments, namely the lateral appendix, the upper, intermediate, and lower vertical lobe, as well as the upper and lower toe (Figure 1C, Figure 3; Masuda-Nakagawa et al., 2014; Eichler et al., 2017; Saumweber et al., 2018).

We next confirmed that APL is GABAergic (Figure 4A-B**’’**; Masuda-Nakagawa et al., 2014) and studied the regional organization of pre- and post-synaptic sites of APL using the APL- GAL4 driver together with the double effector UAS-Dsyd-1::GFP/UAS-DenMark (Owald et al., 2015). Our results show that APL is pre-synaptic in the calyx, whereas it is post-synaptic in both the calyx and the lobes (Figure 4C-D**’’**, **Movie 3**), confirming earlier reports (Masuda-Nakagawa et al., 2014; Eichler et al., 2017). Indeed, we also confirm a functionally inhibitory connection from APL to the KCs in the calyx. Combining transgene expression by the GAL4- UAS and the lexA-lexAop systems, we monitored changes in intracellular Ca^2+^ through changes in fluorescence intensity in the KCs of isolated brain preparations of controls (+>ChR2XXL; KC>GCaMP6m) and upon optogenetic activation of APL (APL_26G02_>ChR2XXL; KC>GCaMP6m). At the relatively lower blue-light intensity (80-ms pulses every 2 s, at 1.8 mW/cm^2^), the control brains showed no specific response to the light; rather, fluorescence intensity decreased slightly across the recording period (Figure 5A, **gray**). Activation of APL resulted in a trend for yet further decreased levels of fluorescence (Figure 5A-B, **blue**). We then exploited the observation that for increased light intensity (4.1 mW/cm^2^) fluorescence intensity increased under control conditions (Figure 5C, **gray**), reflecting the responsiveness of the KCs themselves or of neurons upstream of the KCs. Relative to these fluorescence signals observed in the controls, signals were reduced by APL activation (Figure 5D). Remarkably, even high levels of fluorescence induced by pharmacologically stimulating KCs via the acetylcholine receptor agonist carbamylcholine in control brains (Figure 5E, **gray**) were reduced by simultaneous APL activation (Figure 5E-F). Together, these data confirm a functionally inhibitory effect of APL activation on the KCs in the calyx.

**Figure 5.**
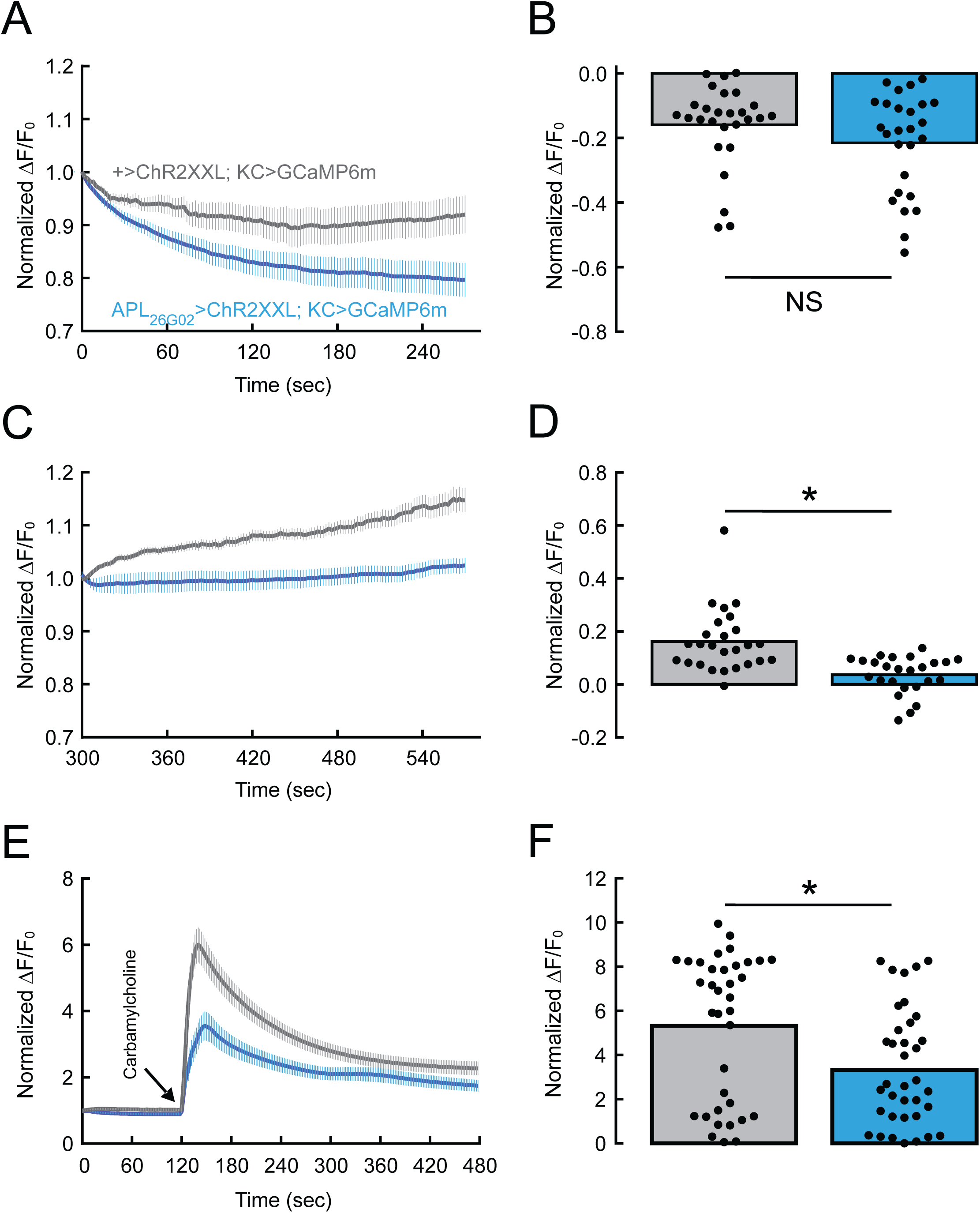
Optogenetic activation of APL can reduce levels of activity in mushroom body Kenyon cells. **(A, B)** At relatively low light intensity (from 0 s to 270 s), activation of APL in isolated brain preparations of APL_26G02_>ChR2XXL; KC>GCaMP6m larvae (blue) had no significant effect on intracellular Ca^2+^ signals in the calycal ROIs/ KCs (Normalized ΔF/F_0_) relative to genetic controls (+>ChR2XXL; KC>GCaMP6m) (gray). For each calycal ROI/ KC, the data are normalized to the beginning of the indicated observation period as a baseline and are plotted over time in (A) (showing mean +/- SEM). The maximum difference from the baseline for each calycal ROI/ KC is plotted in (B) (showing mean and data-point scatter). **(C, D)** In contrast, compared to genetic controls, activation of APL by a more intense light in the same specimen (from 300 s to 570 s) reduced Ca^2+^ signals in the KCs. **(E, F)** Using the same light intensity as in (C-D), bath-application of the acetylcholine receptor agonist carbamylcholine (at 120 s) massively increased Ca^2+^ signals in genetic controls (gray), an effect that was reduced to about half under conditions of APL activation (blue). Preparation and imaging according to Selcho et al. (2017) and Lyutova et al. (2019). The number of brains for genetic controls and APL activation, respectively, was 7 and 9 in (A-D) and 10 and 8 in (E-F); the number of calycal ROIs/ KCs was 27 and 26 in (A-B), 26 and 25 in (C-D), and 36 and 37 in (E-F). NS and * refer to MWW comparisons with p> 0.05 and p< 0.05, respectively. The source data and results of all statistical tests are documented in Extended Data Figure 3-1.

In terms of the organization of APL, the above results also match the situation in first-instar larvae, as shown here for a volume reconstruction of APL generated from the electron microscopy reconstruction of the mushroom body in Eichler et al. (2017) (Figure 6A-B, **Movie 4**). Specifically, this volume reconstruction shows that the relatively slender axonal and dendritic branches of APL arise separately from a thicker neurite (Figure 6C-D), similar to the locust homologue of APL, called GGN (for <u>g</u>iant <u>G</u>ABAergic <u>n</u>euron; Papadopoulou et al., 2011; Ray et al., 2020). The electron microscope dataset of Eichler et al. (2017) further allowed the site of the synapses for the different classes of synaptic partners of APL to be mapped onto its volume reconstruction (Figure 6E-F, **Movies 5-6**). Furthermore, the connectomic data allowed dendrograms of APL to be derived, that is, two-dimensional representations of APL preserving branch lengths and synaptic locations in a topologically correct manner (Figure 7). It can be discerned within such a topology that wherever they coexist, the synapses that APL hosts with mushroom body extrinsic neurons are not segregated from, but are intermingled with, the connections to the mushroom body intrinsic neurons, the KCs (Figure 7B-C). In the lobes, the almost exclusively post-synaptic sites of APL are relatively sparse (Figure 7B-C) and with some variation in topology between the APL neuron of the left and the right brain hemisphere (for the right hemisphere APL neuron, see **Extended Data** Figure 7-1). In the calyx, an analysis of cable length reveals that reciprocal synapses between APL and the KCs are arranged in four synapse-rich center-surround structures such that APL-to-KC synapses are found towards their center, whereas KC-to-APL synapses are located mainly in the surround (Figure 7D).

**Figure 6.**
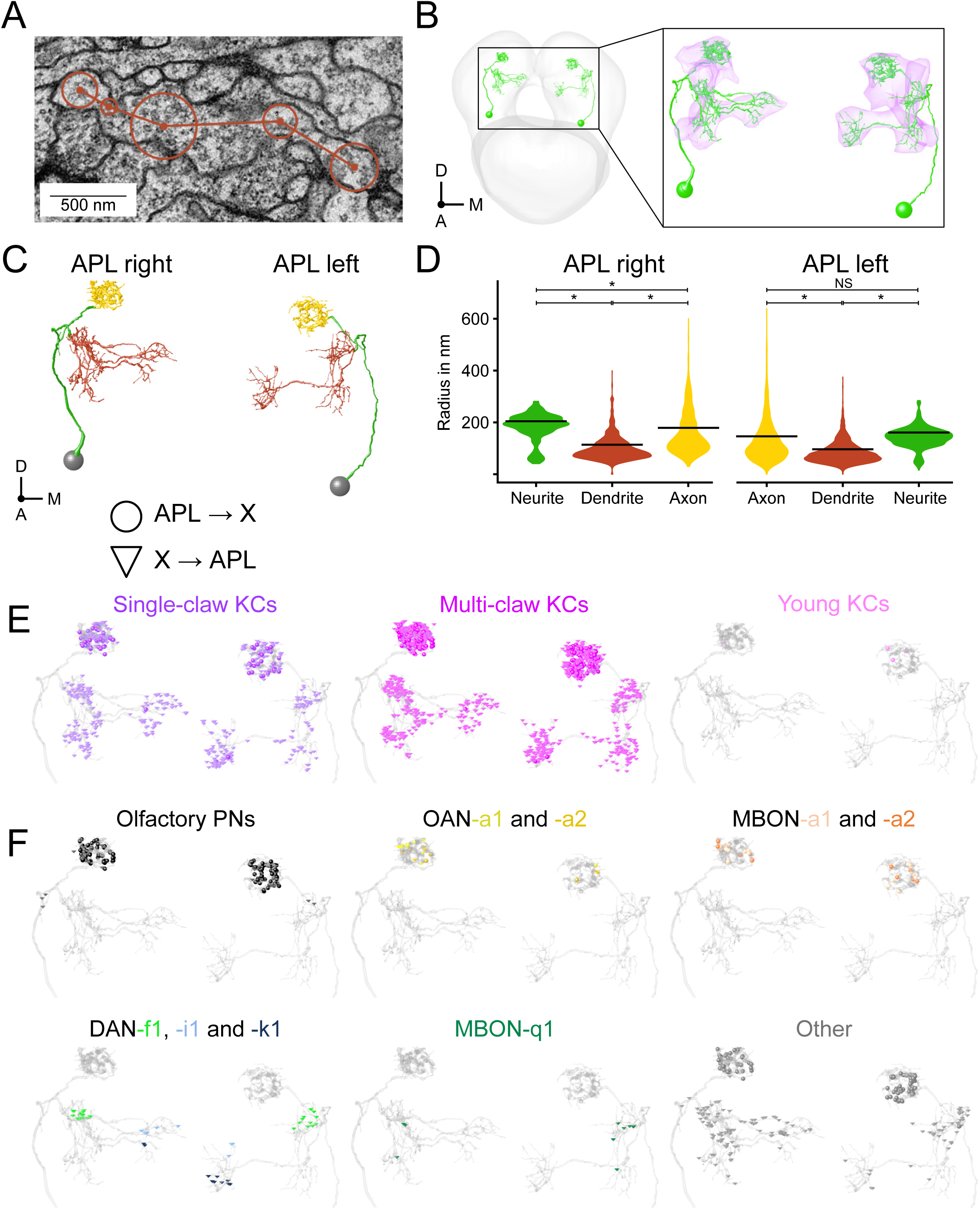
Volume reconstruction of the larval APL neuron. **(A)** Electron microscopy cross-section of the APL neuron in a first-instar larva. Points connected by lines represent the skeletonized reconstruction of the neuron (for details see Eichler et al., 2017). Circles represent radius annotations for volume reconstruction; scale bar: 500 nm**. (B)** Reconstructed volume of the left- and the right-hemisphere APL neuron (green) in the context of the complete central nervous system (left; gray mesh), and in a close-up of the mushroom body region (right; magenta). For a corresponding movie see Movie 4. **(C)** Reconstructed volume of both APL neurons separated into axonal (yellow) and dendritic (red) regions, and the neurite and its branches (green). **(D)** Quantification of the radii of the APL neurons, showing that the neurite is thicker than the axonal regions (significant only for the right hemisphere APL neuron), which in turn are thicker than the dendritic regions. The data are displayed as violin plots; bars represent mean; NS and * refer to MWW comparisons between the APL regions with a Bonferroni-Holm correction (p> 0.05 and p< 0.05, respectively). The source data and results of all statistical tests are documented in Extended Data Figure 3-1. **(E-F)** Pre- and post-synaptic sites annotated by dots and triangles, respectively, selectively for different types of connected neuron, namely: (E) single-claw, multi- claw, and young KCs; (F) neurons with connections in the calyx (top row: olfactory PNs; OAN- a1/a2; MBON-a1/a2), as well as neurons that have been studied elsewhere in functional experiments, such as DAN-i1 (Saumweber et al., 2018; Schleyer et al., 2020), DAN-f1 (Eschbach et al., 2020; Weiglein et al., 2021), DAN-k1 (Saumweber et al., 2018). Neurons with less than two synapses with APL in both hemispheres are shown as “Other”. In (B-C), A: anterior; D: dorsal; M: medial. Corresponding three-dimensional visualizations can be found in Movies 4-6.

**Figure 7.**
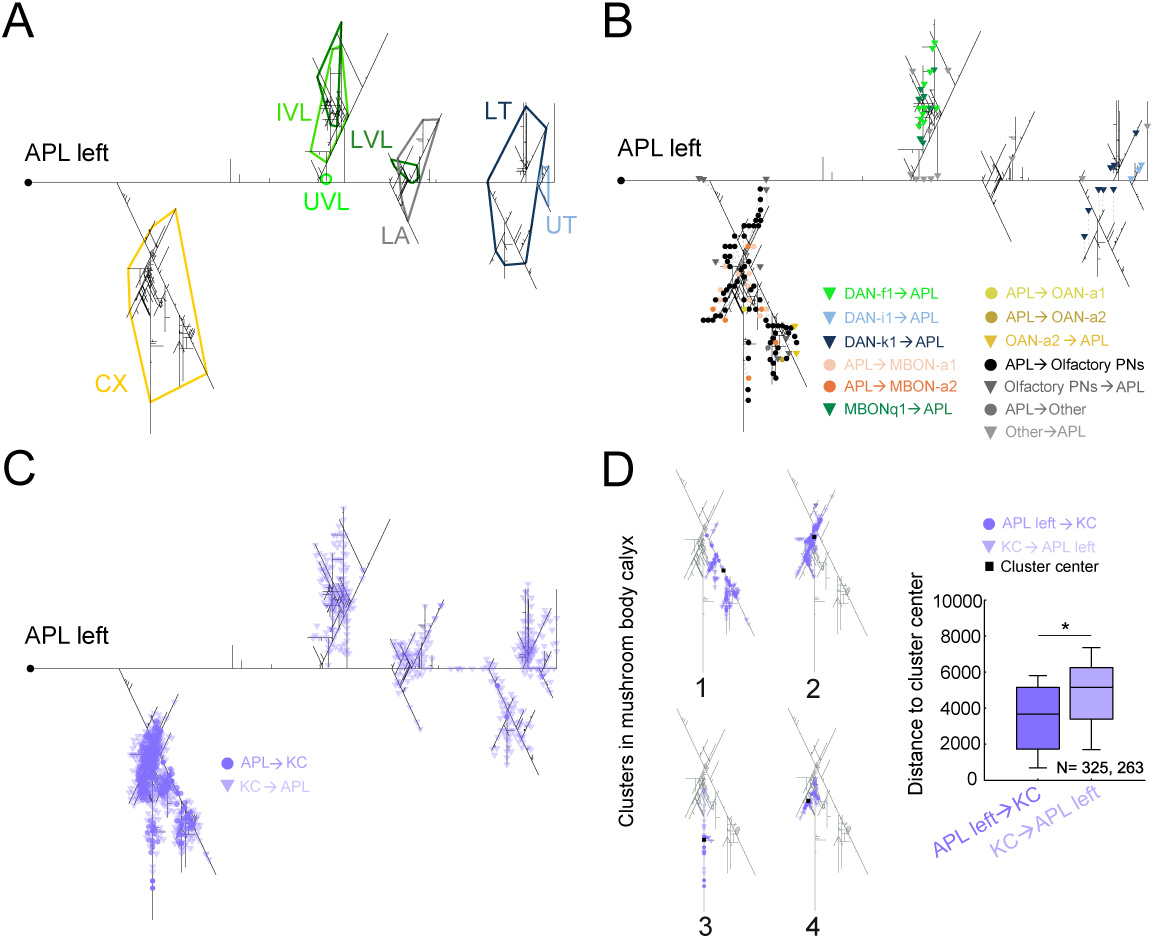
Dendrogram analysis of the larval APL neuron. **(A)** Two-dimensional dendrogram of the APL neuron from the left hemisphere, based on an electron microscope reconstruction in a first-instar larva (data from Eichler et al., 2017). Branch lengths are preserved in a topologically correct manner. The colored envelopes indicate the mushroom body calyx and compartments innervated by APL (see also Figure 1A-B). High- resolution versions of this figure, for the APL neurons of both hemispheres, can be found in Extended Data Figure 7-1. (B) Synapses at their topologically correct site on the left- hemisphere APL neuron with the mushroom body extrinsic neurons indicated. Pre- and post- synaptic sites of APL are annotated with dots and triangles, respectively. For better readability some symbols were displaced and their true locations indicated by a dashed line. High- resolution versions of this figure, for the APL neurons of both hemispheres, can be found in Extended Data Figure 7-2. (C) As in (B), but showing synaptic sites with the mushroom body intrinsic neurons, the Kenyon cells (KCs); dark purple dots and bright purple triangles show APL-to-KC and KC-to-APL synapses, respectively. High-resolution versions of this figure, for the APL neurons of both hemispheres, can be found in Extended Data Figure 7-3. (D) Cluster analysis revealed that calycal synaptic sites of the left-hemisphere APL with the KCs are organized in four clusters (1-4). The accompanying quantification shows geodesic distances of synapses to the center of their respective center-surround structure on the left-hemisphere APL neuron. Most of the APL-to-KC synapses (dark purple dots) are observed towards the center of these clusters (dark square), whereas KC-to-APL synapses (bright purple triangles) are observed mainly in the surround. The data are displayed as box plots, the middle line showing the median, the box boundaries the 25 and 75% quantiles, and the whiskers the 10 and 90% quantiles. The sample sizes (number of synapses) are given within the figure. * refers to MWW comparisons between APL-to-KC and KC-to-APL synapses (* p< 0.05). The source data and results of all statistical tests are documented in Extended Data Figure 3-1. Corresponding analyses, in the context of the full dendrograms and for the APL neurons of both hemispheres, can be found in Extended Data Figure 7-4.

### Regional synaptic polarity of APL across metamorphosis

Given the conserved regional synaptic polarity of APL across larval stages (see preceding section) and given that APL persists into adulthood yet in adults is not regionally polarized (Wu et al., 2013; Lin et al., 2014; Mayseless et al., 2018; Saumweber et al., 2018), we examined how APL develops across metamorphosis. To this end, we used genetically encoded protein ‘tags’ coupled with chemical fluorophore ligands (Kohl et al., 2014; Sutcliffe et al., 2017; Meissner et al., 2018). The synaptic reporter synaptotagmin fused to the tag SNAPm (Syt1- SNAPm) allowed us to label pre-synapses, and the reporter telencephalin fused to the tag CLIPm (TLN-CLIPm) allowed us to label post-synapses (Kohl et al., 2014). These constructs were expressed in APL throughout development using the intersectional driver APLi-GAL4, which specifically expresses in APL of both larvae and adults (Lin et al., 2014; Mayseless et al., 2018) (as stated earlier, APL-GAL4 does not express in adult APLs). In addition, UAS-mCD8::GFP was expressed to visualize APL membranes. In third-instar larvae of the genotype APLi/Syt1:SNAP>mCD8::GFP/TLN:CLIP, pre-synaptic staining was mostly found in the calyx (Figure 8A**’**), whereas post-synaptic staining was distributed in the calyx and the lobes (Figure 8A**’’**), consistent with previous observations (Figure 4; Masuda-Nakagawa et al., 2014). As early as 6 h after puparium formation, we detected pre-synaptic structures that were more punctate (Figure 8B**’**, C’) and observed fewer post-synaptic structures overall (Figure 8B**’’**, C’’), consistent with the previously reported pruning of APL secondary neurites during pupal stages (Mayseless et al., 2018). Interestingly, at 12 h after puparium formation — a stage where APL pruning is almost at its peak (Mayseless et al., 2018) — we could still detect both pre- and post-synaptic structures (Figure 8D-D’’’), although some post-synaptic structures were observed detached from the neurite (Figure 8D’’, D’’’; yellow arrowhead). Nonetheless, the polarized organization of APL in third-instar larvae was no longer observed in the adult stage, as we detected both pre- and post-synaptic markers across both the calyx and the lobes of the adult mushroom bodies (Figure 8E-F’’’; Wu et al., 2013; Lin et al., 2014).

**Figure 8.**
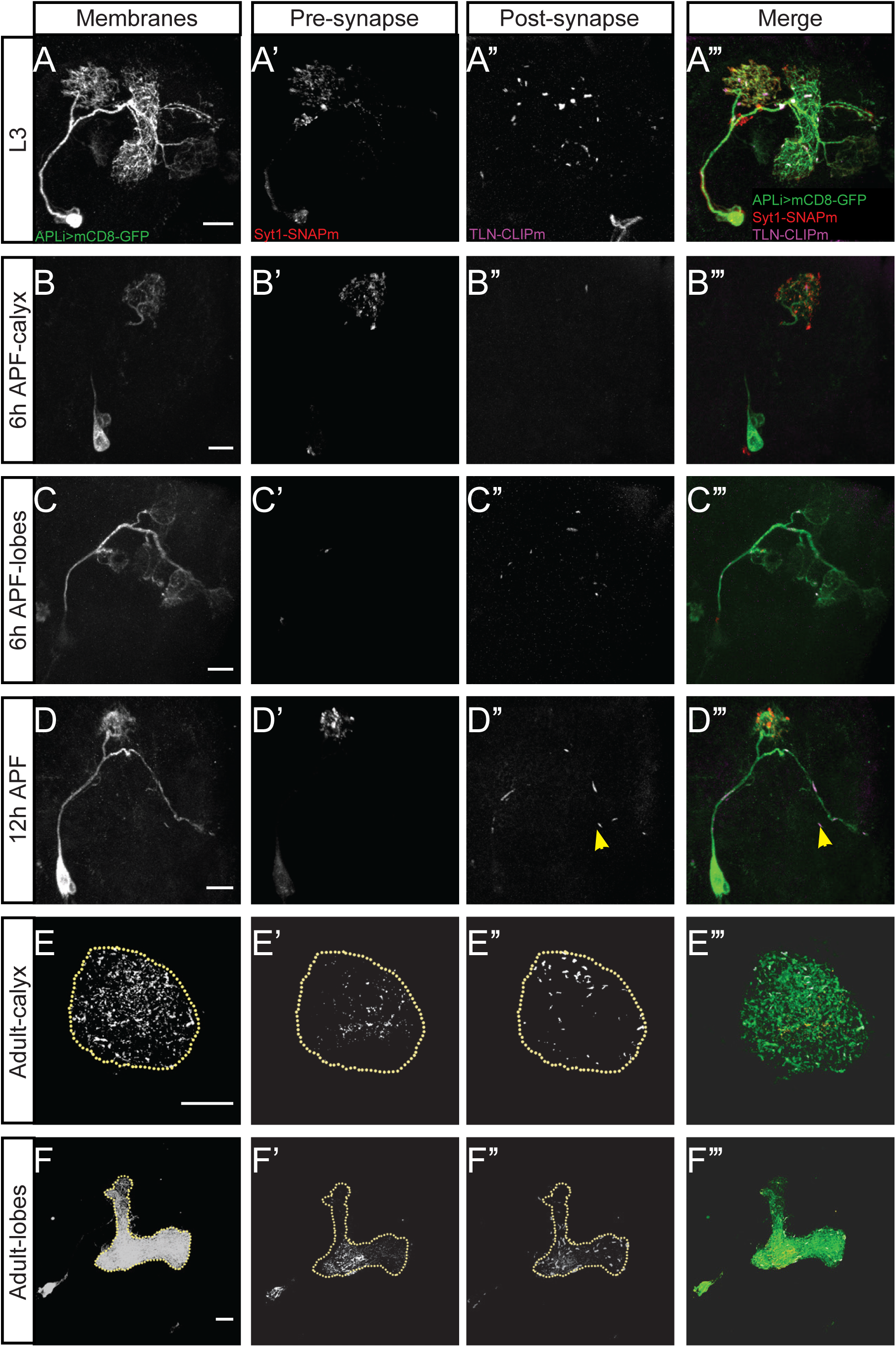
Regional synaptic polarity of APL across metamorphosis. Confocal maximum projection images of stainings for mCD8::GFP, Syt1::SNAP, and TLN::CLIP (Kohl et al., 2014), driven by the APL-specific intersectional driver APLi (Lin et al., 2014; Mayseless et al., 2018) at the following developmental times: **(A-A’’’)** third-instar larva (L3); **(B-C’’’)** 6h after puparium formation (6h APF: calyx: B-B’’’; lobes: C-C’’’); **(D-D’’’)** 12h APF; **(E-F’’’)** adult (calyx: E-E’’’; lobes: F-F’’’). Brains were stained with a polyclonal chicken anti-GFP antibody to label the APL neuron (A-F). To label pre-synapses (A’-F’) and post- synapses (A’’-F’’), the pre-synaptic reporter synaptotagmin was fused to the chemical tag SNAPm (Syt1-SNAPm), and the post-synaptic reporter telencephalin was fused to CLIPm (TLN-CLIPm), respectively (Kohl et al., 2014). Merged images are shown in (A’’’-F’’’). In the third-instar larva, pre-synaptic staining was largely restricted to the calyx (A’), whereas post- synaptic staining was distributed in both the calyx and the lobes (A’’). At 6h APF, both pre- and post-synaptic staining are similarly distributed to that in the larvae (B’-C’’’); notably, pre- synaptic structures seem to be more punctate (B’, C’), and fewer post-synaptic structures are detectable (B’’, C’’). As late as 12h APF, both pre- and post-synaptic structures are still detectable (D’, D’’); post-synaptic structures appear to be detached from the neurite (D’’, D’’’; yellow arrowhead). In adults, both pre- and post-synaptic markers are detectable in both the calyx and the lobes (E-F’’’). The data were acquired with a 40x oil objective; scale bars: 20 μm.

Taken together, our results indicate that whereas APL is regionally polarized throughout larval stages, it undergoes rearrangement during metamorphosis to give rise to a regionally more diffuse organization. In addition, the DPM neuron, one of the main synaptic partners of APL involved in memory consolidation in adults, does not exist in larvae (Pitman et al., 2011; Wu et al., 2011; Eichler et al., 2017; Saumweber et al., 2018). In light of these differences, and despite the rich insights recently gained into the function of APL in adults (Inada et al., 2017; Zhou et al., 2019; Amin et al., 2020; Apostolopoulou and Lin, 2020; Kanellopoulos et al., 2020; Yamagata et al., 2021), a detailed look into the function of the larval APL neuron seemed warranted.

### Memory scores are abolished upon activating APL throughout odor-fructose training

We first asked whether the optogenetic activation of APL affects associative memory formation. Third-instar larvae were trained in a standard Pavlovian conditioning paradigm, using an odor (*n*-amyl acetate) as the conditioned stimulus, and a fructose reward as the unconditioned stimulus (Scherer et al., 2003; Neuser et al., 2005; Saumweber et al., 2011; Michels et al., 2017). One group of larvae received the odor presented together with the fructose reward (paired training), whereas a second group received separate presentations of the odor and the fructose reward (unpaired training). After training, both groups were tested for their odor preference. A difference in odor preference between paired and unpaired training thus reflects associative memory, and is quantified by the memory score. According to convention, positive memory scores reflect appetitive associative memory, whereas negative scores reveal aversive memory (equation 2; Materials and Methods section). Notably, paired and unpaired training both establish associative memory, yet of opposite “sign”. After paired training the odor predicts the occurrence of the reward, leading to an associative increase in odor preference. In contrast, unpaired training establishes the odor as a predictor of the non- occurrence of the reward and supports an associative decrease in odor preference (for a detailed discussion, see Schleyer et al., 2018).

Repeating an experiment from Saumweber et al. (2018), APL was optogenetically activated throughout odor-fructose training (Figure 9). Confirming that report, odor-fructose memory scores in the experimental genotype (APL>ChR2XXL) were reduced to chance levels and were reduced relative to both genetic controls, heterozygous for either only the effector (+>ChR2XXL) or only the driver (APL>+) construct (Figure 9A). The same abolishment of memory scores was observed in a shortened, one-trial version of this experiment (Figure 9B). For practical reasons, this shortened experimental design was used throughout the rest of the study. In addition, the expression of ChR2XXL in APL was directly confirmed by immunohistochemistry (Figure 9C, **Movie 7**). Critically, the behavior of experimentally naïve larvae toward the odor (i.e. innate odor preference) was unaffected by APL activation (Figure 9D; Saumweber et al., 2018). Further, as shown here from offline analyses of video tracking data, APL activation did not affect the modulation of the patterns of locomotion by which these odor preferences came about (i.e. modulations of head cast rate and direction, but not of run speed: Figure 9E-I).

**Figure 9.**
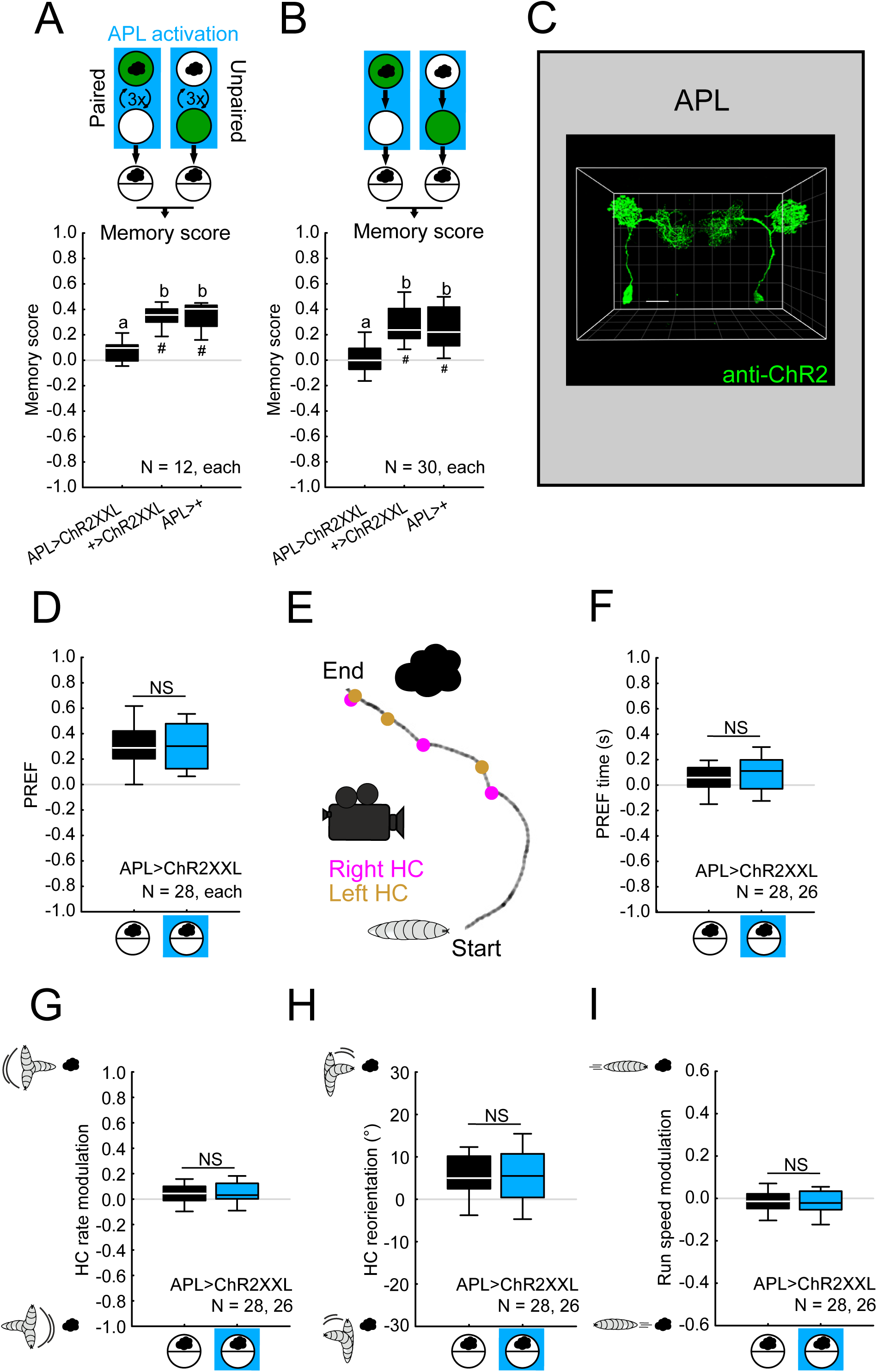
Memory scores are abolished upon activation of APL throughout training. **(A)** Larvae were trained such that in one group of animals, the odor *n*-amyl acetate (black cloud) was paired with the fructose reward (green fill of circle indicating a Petri dish), alternating with blank trials (open circle), whereas in a reciprocal group, the odor was presented unpaired from the fructose reward; please note that here and throughout this study, the sequence of training events was as depicted in half of the cases, and in the reverse order in the other half of the cases. The APL neuron was optogenetically activated with blue light illumination (blue rectangle) during the complete training phase. The larvae from both groups were then tested for their odor preference, and associative memory was quantified by the memory score as the difference in preference between these reciprocally trained groups of animals. Double- heterozygous animals of the genotype APL>ChR2XXL were used for APL activation; larvae heterozygous for either the GAL4 (APL>+) or the effector construct (+>ChR2XXL) were used as the genetic controls. Optogenetic activation of the APL neuron during the complete training phase abolished associative memory scores. **(B)** The same effects were observed in a shortened, one-training-cycle version of this experiment. **(C)** Full projection of the expression pattern from the APL-GAL4 driver crossed to UAS-ChR2XXL in the third-instar larval brain. ChR2XXL is visualized by a primary monoclonal mouse anti-ChR2 antibody and a secondary polyclonal donkey anti-mouse Cy3 antibody. Confirming our results from Figures 2-4, this reveals strong and reliable transgene expression in the APL neuron of both hemispheres (anti- ChR2XXL; green). The data were acquired with a 63x glycerol objective; scale bar and grid spacing: 20 µm. For a corresponding movie see Movie 7. **(D)** The behavior of experimentally naïve larvae of the experimental genotype (APL>ChR2XXL) toward *n*-amyl acetate (black cloud) was tested, without APL being activated during testing or with APL activated (blue square). Naïve odor preference was unaffected by APL activation. **(E-I)** The behavior of larvae in (D) was videorecorded and analyzed offline as described by Paisios et al. (2017). (E) shows a short sample from a video recording of a larva with successive runs and head casts (HCs). Displayed is the track of the midpoint. Magenta and orange dots indicate right and left HCs, respectively. Specifically, three features of locomotion were analyzed in addition to the olfactory preference (i.e. the time spent by the larvae on the odor and the non-odor side, F): the HC rate modulation (G), the HC reorientation (H), and the run speed modulation (I). In all cases APL activation had no effect. The data are displayed as box plots, the middle line showing the median, the box boundaries the 25 and 75% quantiles, and the whiskers the 10 and 90% quantiles. The sample sizes (number of biological replications) and the genotypes are given within the figure. In (A-B), different letters refer to significant differences between groups in MWW comparisons with a Bonferroni-Holm correction (p< 0.05), as specified in the section “Experimental design and statistical analyses”. In (D, F-I), NS refers to the absence of significance between groups in MWW comparisons (NS p> 0.05). In (A-B), ^#^ refers to OSS comparisons to chance levels (i.e. to zero), also with a Bonferroni-Holm correction (^#^ p< 0.05). The source data and results of all statistical tests are documented in Extended Data Figure 3-1.

### Activating APL either in the presence or in the absence of the odor reduces memory scores

As argued in Saumweber et al. (2018), the abolishment of memory scores upon activation of APL during the complete training phase (Figure 9) may arise because APL provides an inhibitory GABAergic signal onto the KCs (Figure 5; Masuda-Nakagawa et al., 2014). Taking the argument to the extreme, the activation of APL would silence the KCs, preventing a proper odor representation in the mushroom body and thereby also preventing odor-fructose memory formation. If so, memory formation should be disrupted when APL is activated while the odor is presented, but should not be disrupted when APL is activated while the odor is not presented. To our surprise, however, *in both cases* odor-fructose memory scores were partially reduced compared to a control condition in which APL was not activated at all (Figure 10, **left**). As regards these residual memory scores, we considered the interpretation of odor-fructose memory as a learned search for the fructose reward (Saumweber et al., 2011; Schleyer et al., 2011). This interpretation implies that memory is behaviorally expressed if the sought-for fructose reward is indeed absent during the test, but that memory is not expressed if the testing is carried out in the presence of the sought-for fructose reward. This was indeed the case in all three conditions, namely (i) when APL was not activated during training at all, and (ii) when APL was activated during odor presentation or (iii) in the absence of odor (Figure 10, **right**); please note that innate olfactory behavior is not changed in the presence of fructose or other tastants: Schleyer et al., 2011). In other words, the residual memory scores after APL activation during either period of the training also reflect a search for the fructose reward.

**Figure 10.**
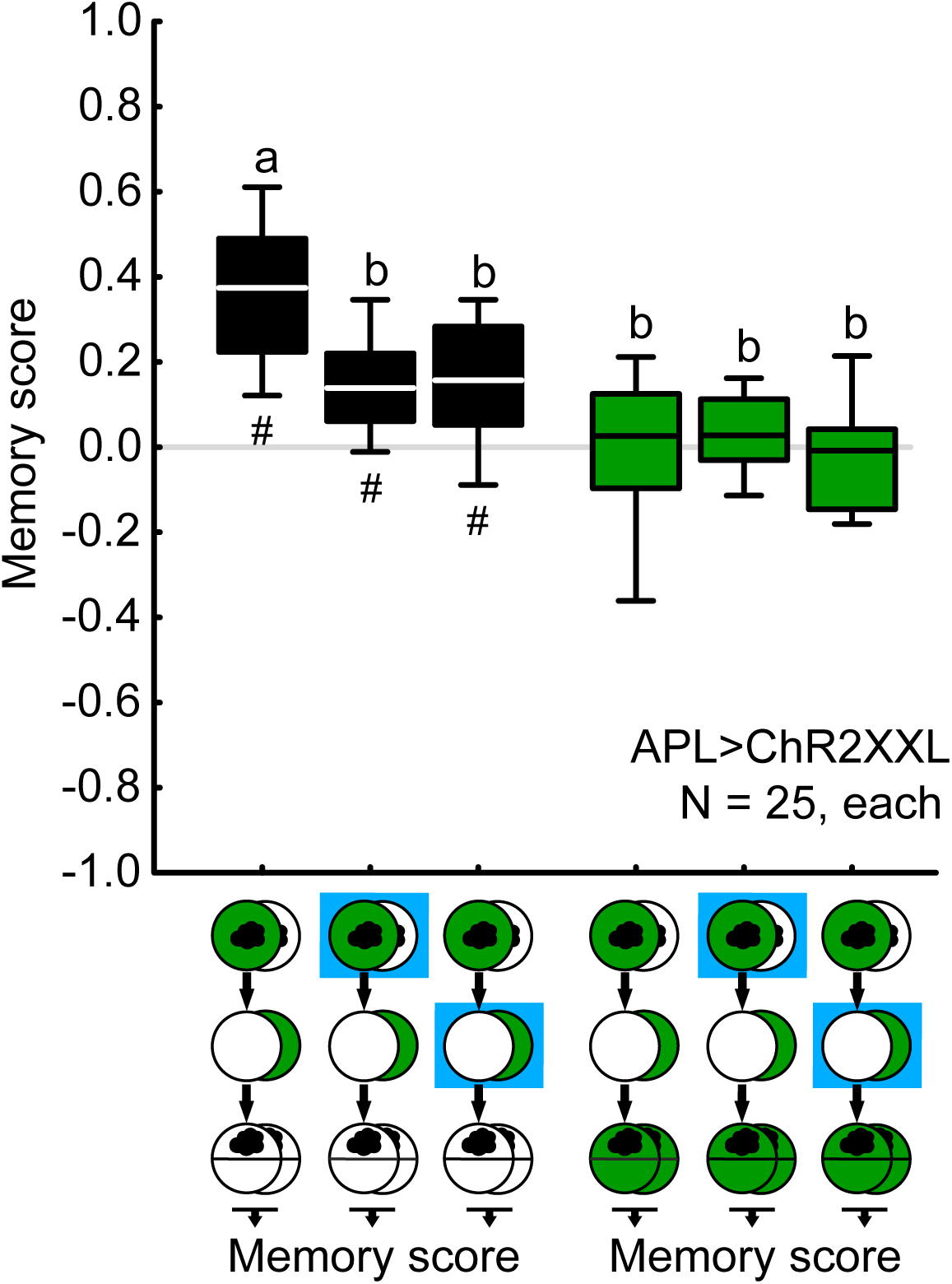
Activation of APL only in the presence or only in the absence of the odor reduces memory scores. Optogenetic activation of APL (blue square) either only when the odor was presented during training, or only when the odor was not presented during training, reduced memory scores to about half the level of control animals that did not receive any APL activation (black-filled box plots). Testing the animals in the presence of the training reward (i.e. fructose) abolished the behavioral expression of memory in all cases (green-filled box plots). The sample sizes and the genotype are given within the figure. ^#^ refers to OSS comparisons to chance levels (i.e. to zero) with a Bonferroni-Holm correction (^#^ p< 0.05); different letters refer to significant differences between groups in MWW comparisons also with a Bonferroni-Holm correction (p< 0.05). The source data and results of all statistical tests are documented in Extended Data Figure 3-1. Other details as in Figure 9.

### Differential effects of activating APL only in the presence or only in the absence of the odor

As mentioned, it was unexpected that odor-fructose memory was impaired by activation of APL during the odor-absent periods of training. In order to understand this result, we separately analyzed the odor preference scores underlying the memory scores from Figure 10. In all three cases — (i) when APL was not activated at all during training, (ii) when it was activated while the odor was presented, and (iii) when it was activated while the odor was not presented — odor preference scores after paired *vs*. unpaired training were indistinguishable from each other when the fructose reward was present during testing (open boxes in Figure 11A-C). In other words, in all cases learned search ceased once the sought-for reward was found. As discussed in detail in Schleyer et al. (2018), this allows the odor preferences after paired and unpaired training to be pooled in order to determine baseline levels of odor preference, cleared of associative memory (stippled lines in Figure 11A-C). In all three cases, these baseline preference scores were intermediate between the paired-trained and the unpaired-trained animals that were tested in the absence of fructose, consistent with earlier reports (Schleyer et al., 2018). This is adaptive because after paired training the larvae search for fructose where the odor is, whereas after unpaired training they search for fructose where the odor is not, and accordingly in either case their search is suppressed in the presence of the sought-for fructose. Important for the current context, however, is that these baseline levels varied strikingly with the contingency between APL activation and odor presentation (Figure 11D): as compared to the control baseline scores when APL was not activated at all (stippled line in Figure 11A and plotted in Figure 11D, **left**), the baseline scores were increased when APL was activated in the presence of the odor (stippled line in Figure 11B; plotted in Figure 11D, **middle**), and were decreased when APL was activated in the absence of the odor (stippled line in Figure 11C; plotted in Figure 11D, **right**). In other words, activation of APL paired with odor increased odor preferences, whereas activation of APL unpaired from odor presentation decreased odor preferences (Figure 11D) – as if, above and beyond the fructose that we intended to be the only reward in these experiments, activation of APL also had a rewarding effect! The next experiment tested this hypothesis.

**Figure 11.**
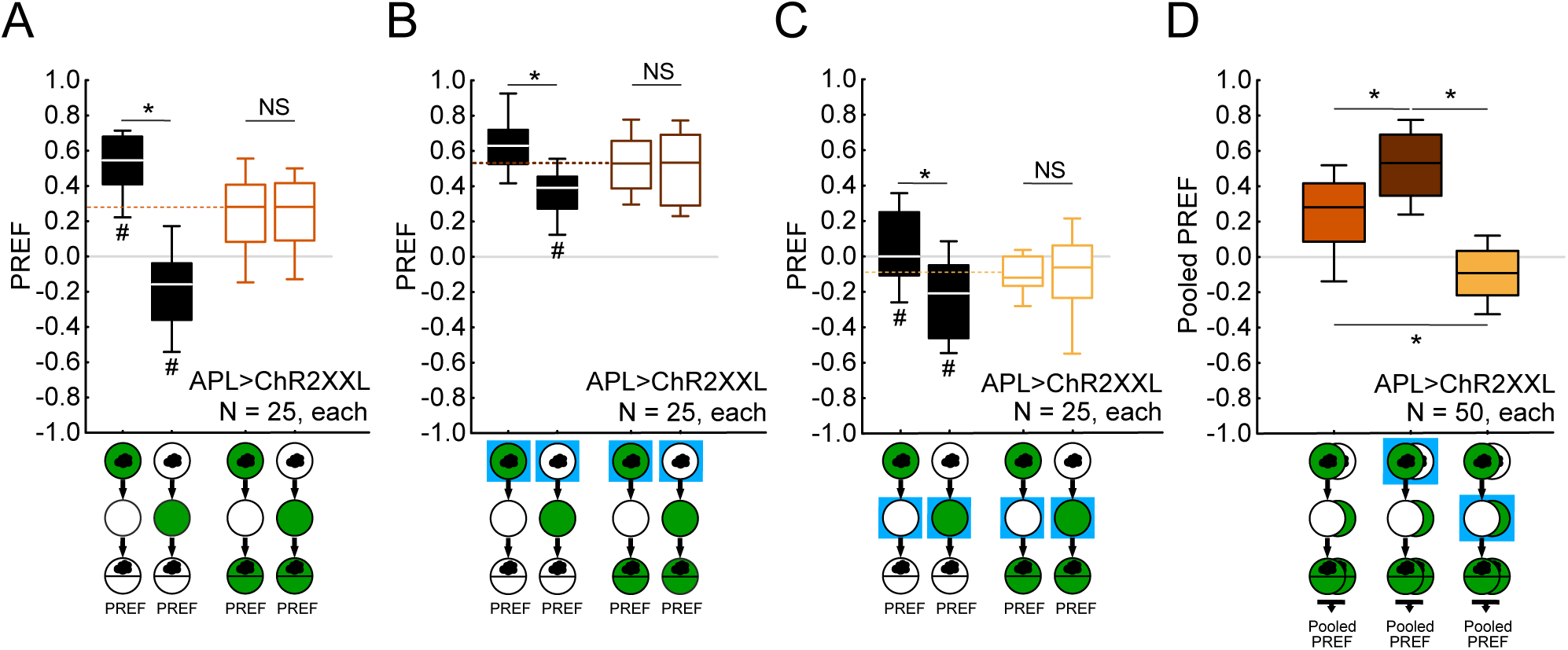
Activation of APL only in the presence or only in the absence of the odor has differential effects on odor preference. **(A)** Examination of the preference scores (PREF) underlying the associative memory scores from Figure 10 reveals that odor preference scores are higher after paired than after unpaired training with odor and fructose reward (black-line plots to the left), a difference that is abolished when testing is carried out in the presence of the training reward (right-most colored plots). This is adaptive because learned search for the reward is obsolete in its presence. These preference scores can thus be pooled to serve as the baseline odor preference cleared of associative memory (stippled line). This reveals that odor preference scores are higher than the baseline after paired training and lower than the baseline after unpaired training. (B-C) shows the same upon activation of APL during training (blue square), whether only during odor presentation (B), or unpaired from odor presentation (C). Remarkably, baseline levels of odor preference differ between these three training conditions (D): As compared to the control condition without APL activation, baseline odor preference scores (Pooled PREF) are increased when APL is activated together with odor presentation, and decreased when APL is activated unpaired from odor presentation. The sample sizes and the genotype are given within the figure. In (A-D), * and NS refer to MWW comparisons between groups with a Bonferroni- Holm correction (* p< 0.05; NS p> 0.05). In (A-C), ^#^ refers to MWW comparisons to baseline levels of odor preference, also with a Bonferroni-Holm correction (^#^ p< 0.05). The source data and results of all statistical tests are documented in Extended Data Figure 3-1. Other details as in Figures 9-10.

### Activating APL has a rewarding effect

To test whether optogenetically activating APL has a rewarding effect, animals were trained such that the odor was presented either paired or unpaired with APL activation instead of with the fructose reward. This established positive memory scores in the experimental genotype, differing from the genetic controls (Figure 12A). Thus, APL activation during training has a rewarding effect and can establish appetitive, associative memory. In turn, optogenetically silencing APL leads to an aversive memory (Figure 12B). Throughout the rest of the study, we decided to focus on the rewarding effect of APL upon its activation. We found that the resulting appetitive “odor-APL memory” was transient and lasted for less than 10 min (Figure 12C), as is the case for odor-fructose memory after one training trial (Weiglein et al., 2019) and for appetitive olfactory memories formed by optogenetic activation of large sets of KCs (Lyutova et al., 2019). A rewarding effect of APL activation was likewise observed when a brief- stimulation protocol was used (Figure 12D-E). In addition, offline analysis of video-recorded larval locomotion revealed that the same aspects of larval locomotion were modulated by odor- APL memory (Figure 13) as previously shown for similarly strong odor-taste reward associative memories, namely the rate of head casts and their orientation but not run speed (Schleyer et al., 2015b; Paisios et al., 2017; Saumweber et al., 2018; Thane et al., 2019; Schleyer et al., 2020). Inspired by what has been reported on fructose as a taste reward (Schleyer et al., 2015a; see also Figure 10, Figure 11), we reasoned that if odor-APL memory scores reflect a learned search for the training reward (which is APL activation in the present case), these memory scores should be abolished if the sought-for reward is present during the test. We therefore repeated the experiment from Figure 12A and added an experimental condition whereby APL was also activated during testing. This prevented the behavioral expression of appetitive odor-APL memory (Figure 14A; for a similar effect of DAN activation see Schleyer et al., 2020). The same was observed for a two-odor, differential conditioning version of the paradigm, using 1-octanol as the second odor (Figure 14B). We indeed find it striking that APL activation is effective as a reward even in differential conditioning, because it implies that an associable, odor-specific representation can be established in the mushroom body under the condition of an optogenetically activated APL neuron. In line with our earlier results from Figure 9D, naïve odor preferences were unaffected by APL activation (Figure 14C-E). Thus, APL activation has two kinds of effect previously reported for taste rewards: it both induces associative memory when paired with odor during training (Figure 12, Figure 14; with the same locomotor ‘footprint’ as for taste rewards: Figure 13), and it terminates the search behavior that is based on this memory during the test (Figure 14). These two effects of reward are adaptive because they help the animals to search for the reward, and prevent them from drifting away from a reward once it is found, respectively. Both of these reward-like effects of APL activation, plus the lack of any effect of APL activation on naïve odor preference, were confirmed using Chrimson as the effector (Figure 15A-D) – although the effect of APL activation on search behavior during testing was only partial for Chrimson (Figure 15C).

**Figure 12.**
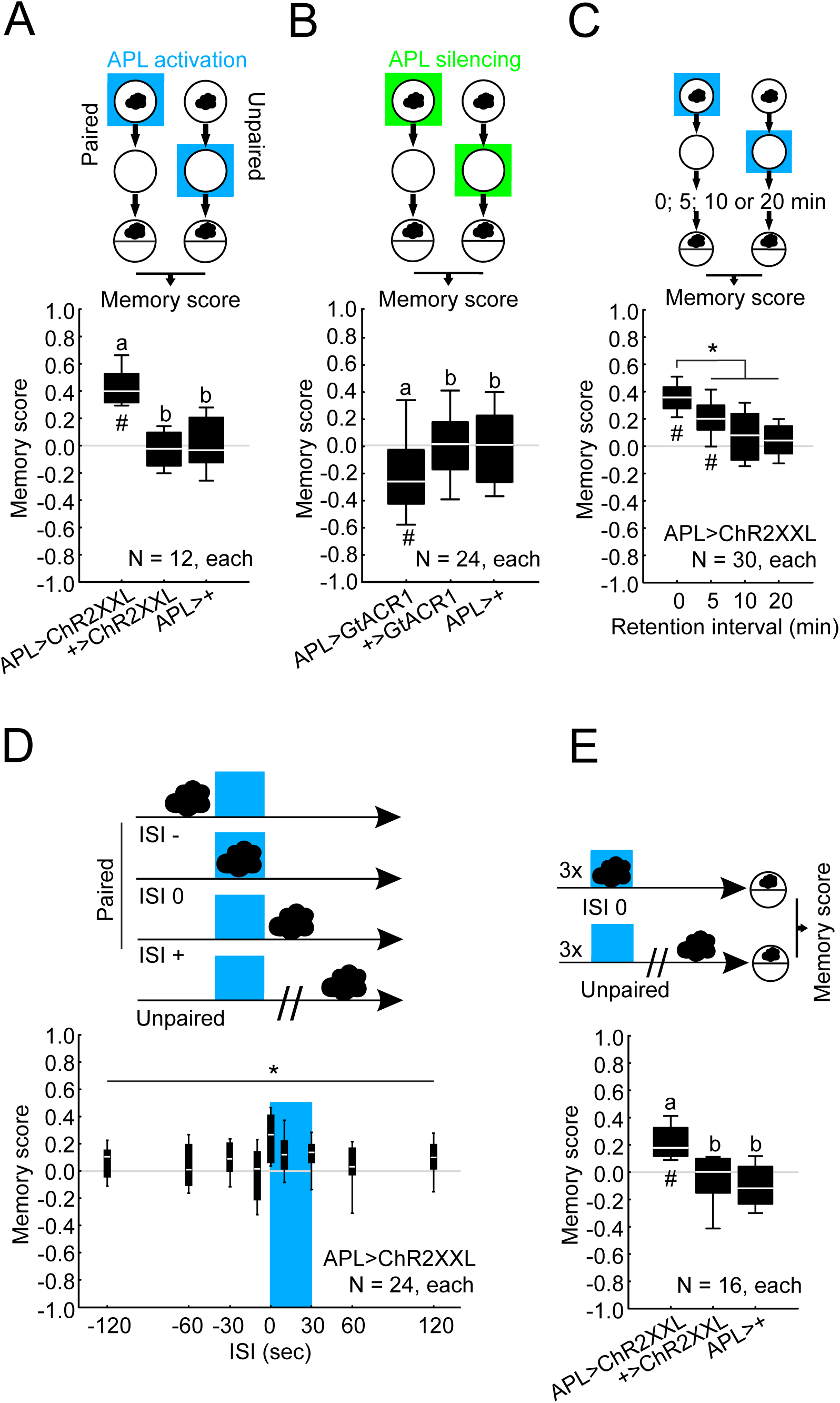
Activating APL has a rewarding effect. **(A)** Animals were trained by presenting odor either paired with, or unpaired from, activation of APL using ChR2XXL as the effector through blue light illumination (blue square). The effect of APL activation as a reward is quantified by positive memory scores, differing significantly from the genetic controls. **(B)** Procedure as in (A) except that APL was optogenetically silenced using GtACR1 as the effector through green light illumination (green square). The effect of APL silencing as a punishment is quantified by negative memory scores, differing significantly from the genetic controls. **(C)** Larvae of the experimental genotype (APL>ChR2XXL) were trained as described in (A) and tested either immediately after training (retention interval 0 min) or 5, 10 or 20 min after training. Expression of odor-APL memory was observed immediately after training and was still detectable at a 5 min retention interval; it was significantly reduced compared to immediate testing when assessed at 5, 10 or 20 min retention intervals. **(D)** Larvae of the experimental genotype (APL>ChR2XXL) were trained as in (A) (i.e. paired or unpaired) but with modifications of the paradigm in accordance with Weiglein et al. (2020). Specifically, odor presentation and APL activation lasted for 30 s each with different timings relative to their onset (inter-stimulus interval, ISI): either the odor was presented *before* the APL activation (negative ISI values), *during* the APL activation (ISI 0), or *after* the APL activation (positive ISI values); in all cases reciprocal training involved odor presentation unpaired from APL activation. Three training trials were performed, followed by the test of odor preference. Memory scores differed according to the ISI. **(E)** Repetition of the experiment from (D) for simultaneous presentation of odor and APL activation (ISI 0), including genetic controls. Positive memory scores for the experimental genotype (APL>ChR2XXL) indicate that a brief stimulation of APL is sufficient to be rewarding, an effect that was not observed in the genetic controls. The sample sizes and the genotypes are indicated within the figure. In (A-B, E), different letters refer to significant differences between groups in MWW comparisons with a Bonferroni-Holm correction (p< 0.05); in (C), **\*** refers to significant differences between groups in MWW comparisons with a Bonferroni-Holm correction (**\*** p< 0.05); in (D), * refers to a KW multiple-group comparison (**\*** p< 0.05); in (A-C, E), ^#^ refers to OSS comparisons to chance levels (i.e. to zero), also with a Bonferroni-Holm correction (^#^ p< 0.05). The source data and results of all statistical tests are documented in Extended Data Figure 3-1. Other details as in Figures 9-11.

**Figure 13.**
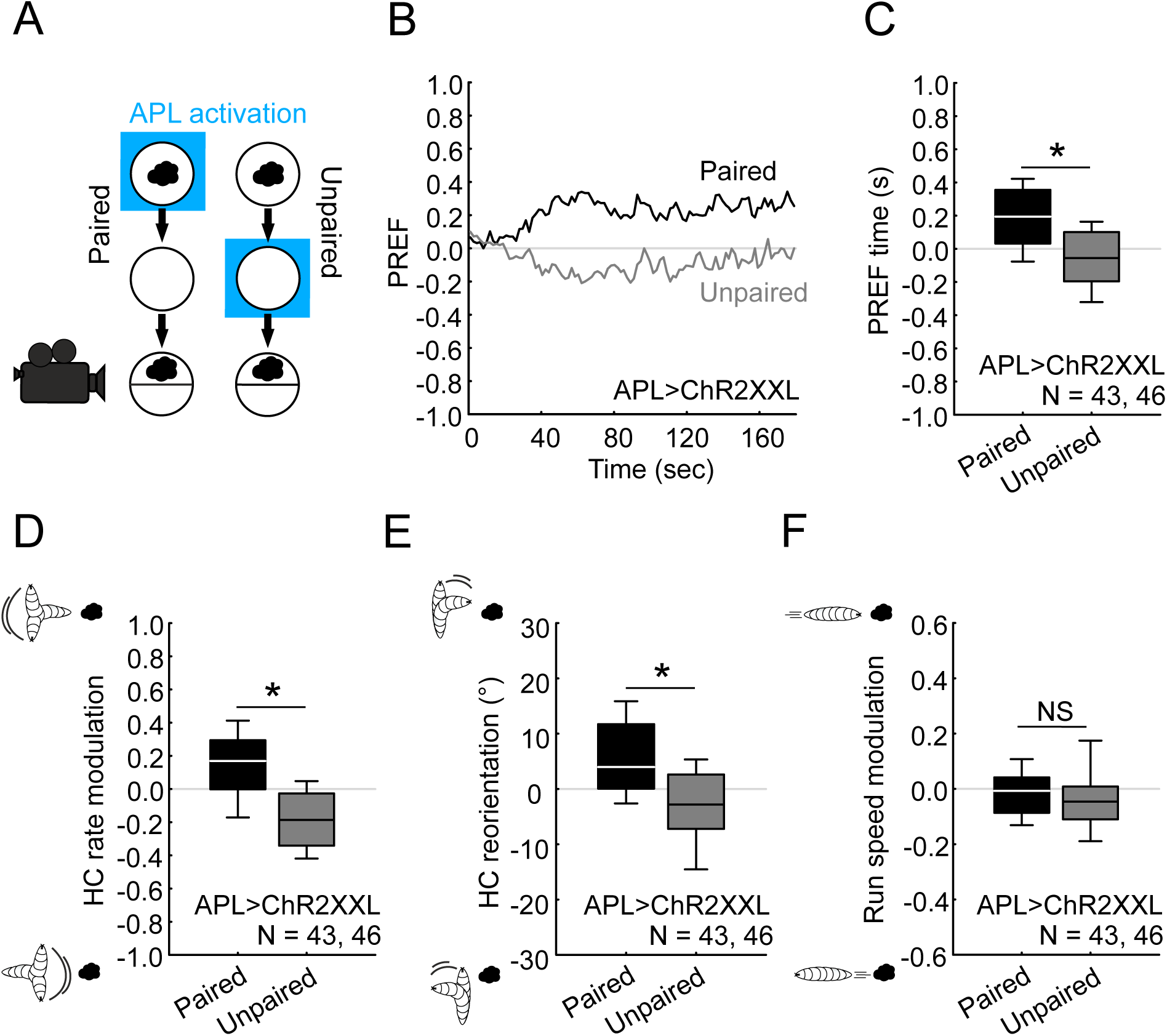
Locomotor ‘footprint’ of odor-APL memory. **(A)** The behavior of larvae of the genotype APL>ChR2XXL was videorecorded after paired or unpaired training with odor and APL activation. **(B)** Larvae showed a higher preference for the odor after paired training than after unpaired training; dataset split into 100 bins (1.8 s, each), showing the median of odor preferences across Petri dishes over time. **(C)** Larvae from the paired group spent more time on the odor side than on the non-odor side during testing, whereas the contrary was observed for the unpaired group. **(D)** Paired-trained larvae exhibited more HCs when crawling away from the odor than when moving towards it; the opposite was observed for the unpaired group. **(E)** Larvae from the paired group oriented their HCs more in the direction of the odor compared to larvae from the unpaired group. **(F)** The run speed when heading towards versus when heading away from the odor did not differ between paired- and unpaired-trained animals. Analyses in (B-F) are based on data available from the experiments shown in Figure 12A, C and Figure 14A. Similar results were observed using Chrimson as the optogenetic effector (not shown). The sample sizes (number of biological replications) and the genotype are given within the figure. In (C-E), * refers to significant differences between groups in MWW comparisons (* p< 0.05). In (F), NS refers to the absence of significance between groups in MWW comparisons (NS p> 0.05). The source data and results of all statistical tests are documented in Extended Data Figure 3-1. Other details as in Figures 9-12.

**Figure 14.**
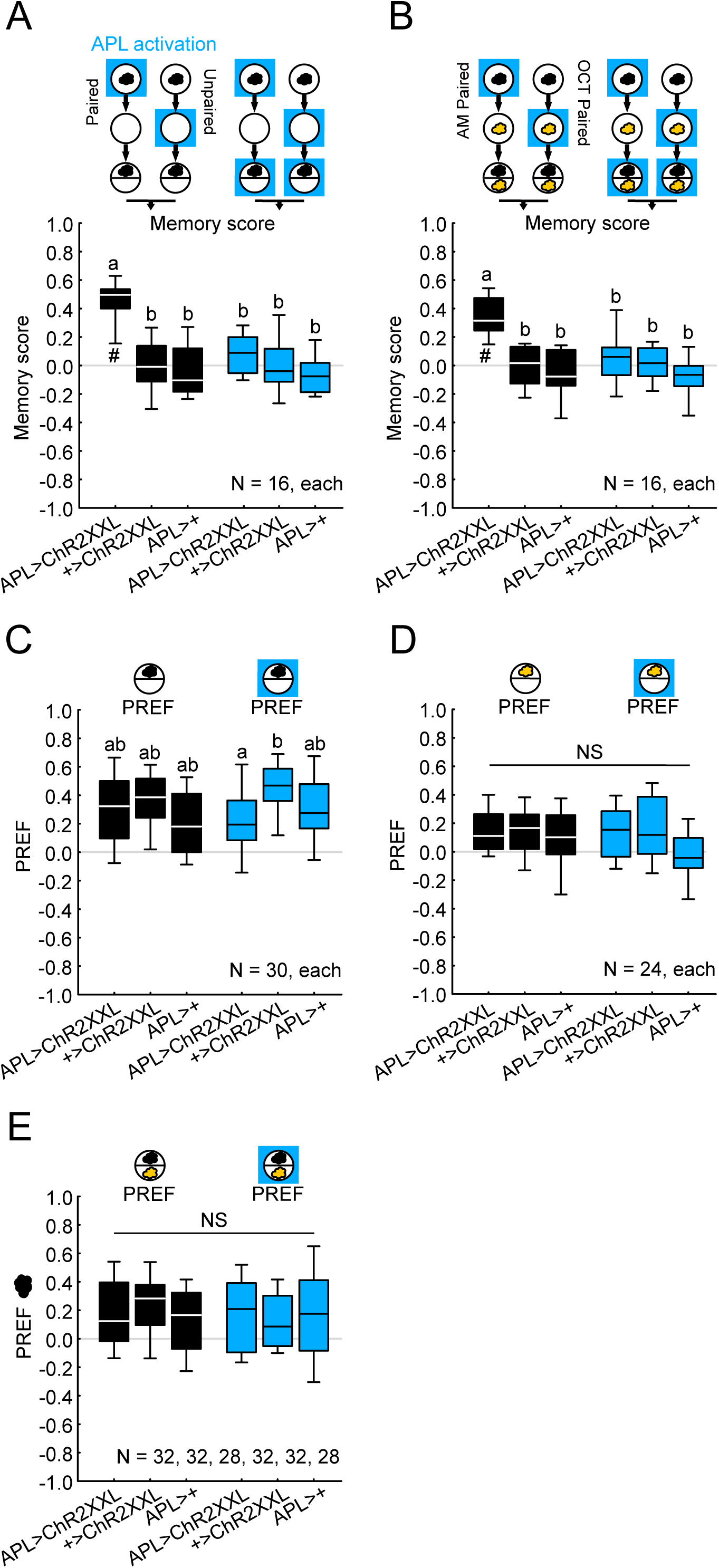
Activation of APL during testing prevents the behavioral expression of appetitive odor-APL memory. **(A)** Repetition of the experiment from Figure 12A, confirming that APL activation has a rewarding effect (black-filled box plots). Activating APL during testing as well prevented the behavioral expression of appetitive odor-APL memory (blue-filled box plots). **(B)** Larvae were trained and tested as in (A), except that in a differential conditioning protocol, 1-octanol was used as a second odor (OCT; yellow cloud) in all training trials in which *n*-amyl acetate (AM; black cloud) was not presented. Presenting one of the two odors paired with APL activation induced odor-specific appetitive memory; as in (A), testing the animals while activating APL prevented the behavioral expression of memory. **(C)** The behavior of experimentally naïve larvae toward *n*-amyl acetate (black cloud) was tested either without APL activation or with APL activation during the test. Naïve odor preference in the experimental group was unaffected by APL activation (APL>ChR2XXL) (see also Figure 9D), with the caveat that it did differ from the effector (+>ChR2XXL), but not from the driver control (APL>+). **(D)** As in (C), except that OCT was used as a single odor (yellow cloud). Naïve odor preference in the experimental group was unaffected by APL activation (APL>ChR2XXL), and did not differ from the genetic controls. **(E)** As in (C-D), except that OCT was used as a second odor (yellow cloud). Again, naïve odor preference in the experimental group was unaffected by APL activation (APL>ChR2XXL) and did not differ from the genetic controls. The sample sizes and the genotypes are given within the figure. In (A-C), # refers to OSS comparisons to chance levels (i.e. to zero) with a Bonferroni-Holm correction (# p< 0.05); different letters refer to significant differences between groups in MWW comparisons, also with a Bonferroni-Holm correction (p< 0.05). In (D-E), NS refers to the absence of significance between groups in KW comparisons (NS p> 0.05). The source data and results of all statistical tests are documented in Extended Data Figure 3-1. Other details as in Figures 9-13.

**Figure 15.**
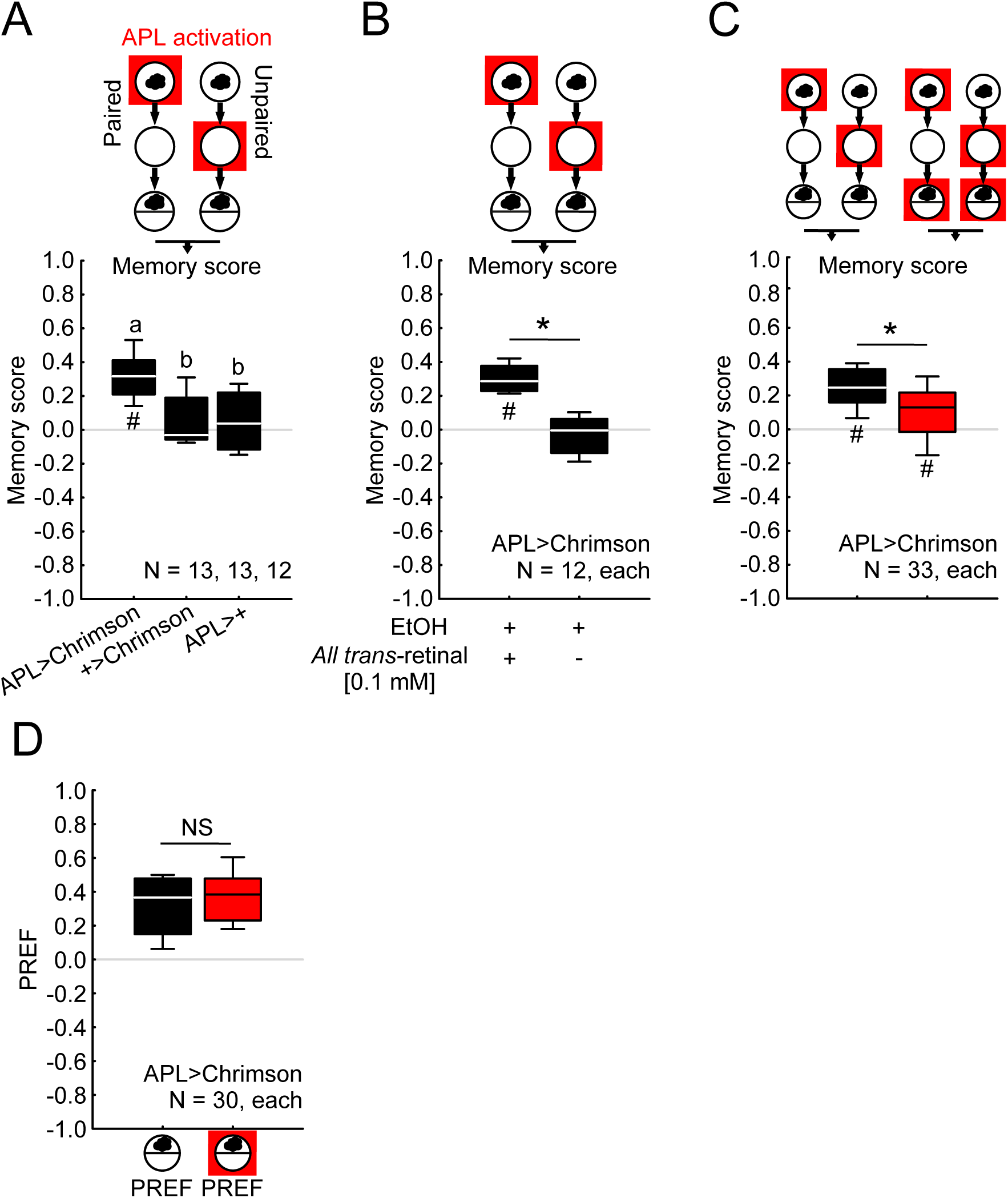
Activating APL with Chrimson has a rewarding effect. **(A)** The rewarding effect of APL activation was confirmed using Chrimson as the effector and red light illumination (red square), and quantified through positive memory scores in the experimental group (APL>Chrimson), differing significantly from the genetic controls. Transgenic flies were raised on standard food supplemented with retinal (100 mM final concentration). **(B)** Larvae of the experimental genotype (APL>Chrimson) were trained and tested after being raised on food either supplemented with retinal (final concentration in ethanol [EtOH 99.9%] 100 mM), or without retinal (food medium supplemented with EtOH only). The rewarding effect of APL activation was observed in retinal-fed animals, but was not observed without retinal feeding. **(C)** The behavioral expression of odor-APL memory was reduced but not abolished by testing the animals while APL was activated (red-filled box plot). **(D)** The behavior of experimentally naïve larvae of the genotype APL>Chrimson toward *n*-amyl acetate (black cloud) was tested, either without APL activation or with APL activated during testing (red square). Naïve odor preference remained unaffected by APL activation. The sample sizes and the genotypes are given within the figure. In (A), different letters refer to significant differences between groups in MWW comparisons with a Bonferroni-Holm correction (p< 0.05). In (B-C), * refers to significant differences between groups in MWW comparisons, also with a Bonferroni-Holm correction (* p< 0.05). In (D), NS refers to the absence of significance between groups in MWW comparisons (NS p> 0.05). In (A-C), ^#^ refers to OSS comparisons to chance levels (i.e. to zero) with a Bonferroni-Holm correction (^#^ p< 0.05). The source data and results of all statistical tests are documented in Extended Data Figure 3-1. Other details as in Figures 9-14.

### Manipulating activity in the calyx MBONs has no reinforcing effect

Considering the circuit mechanisms by which APL activation exerts a rewarding effect, we first focused on the calyx MBONs to which APL is pre-synaptic (MBONa1 and MBONa2; also known as “Odd” neurons: Figure 6F, Figure 7B, **Movie 6**; Slater et al., 2015; Eichler et al., 2017; Saumweber et al., 2018). We reasoned that if activation of the GABAergic APL neuron exerts its rewarding effect by inhibiting the calyx MBONs, then optogenetically silencing these MBONs should also have a rewarding effect. Using the chloride channel GtACR1 as the effector, however, this was found not to be the case (Figure 16A). We then considered the possibility that, unlike the KCs (Figure 5), the calyx MBONs might actually be activated rather than inhibited by activation of APL, e.g. through GABA-induced chloride spikes as reported for insect neurons (Ryglewski et al., 2017) or by yet-to-be-identified excitatory transmitters co- released by APL. We thus repeated the same experiment but this time activated the MBONs: again, no rewarding effect was observed upon such manipulation (Figure 16B). Before ruling out the involvement of the two calyx MBONs in the rewarding effect of APL, however, it seemed important to test for the effects of manipulating each of them separately. This is because activation of MBONa1 and MBONa2 induces approach and avoidance, respectively (Eschbach et al., 2021). We therefore reasoned that they might exert a rewarding and punishing effect, respectively, which would sum to zero when both these MBONs were manipulated together. However, neither silencing nor activating either one of the calyx MBONs yielded evidence of such oppositely-reinforcing effects (Figure 16C-D, **Extended Data** Figure 16-1; we included groups tested in the presence of light because we reasoned that, similar to what has been observed for tastant punishment (Gerber and Hendel, 2006; Selcho et al., 2009; Schleyer et al., 2011; Widmann et al., 2016; Weber et al., 2022), this might promote aversive memory expression). These results suggest that the rewarding effect of APL activation does not involve APL-to-MBONa1/a2 connections.

**Figure 16.**
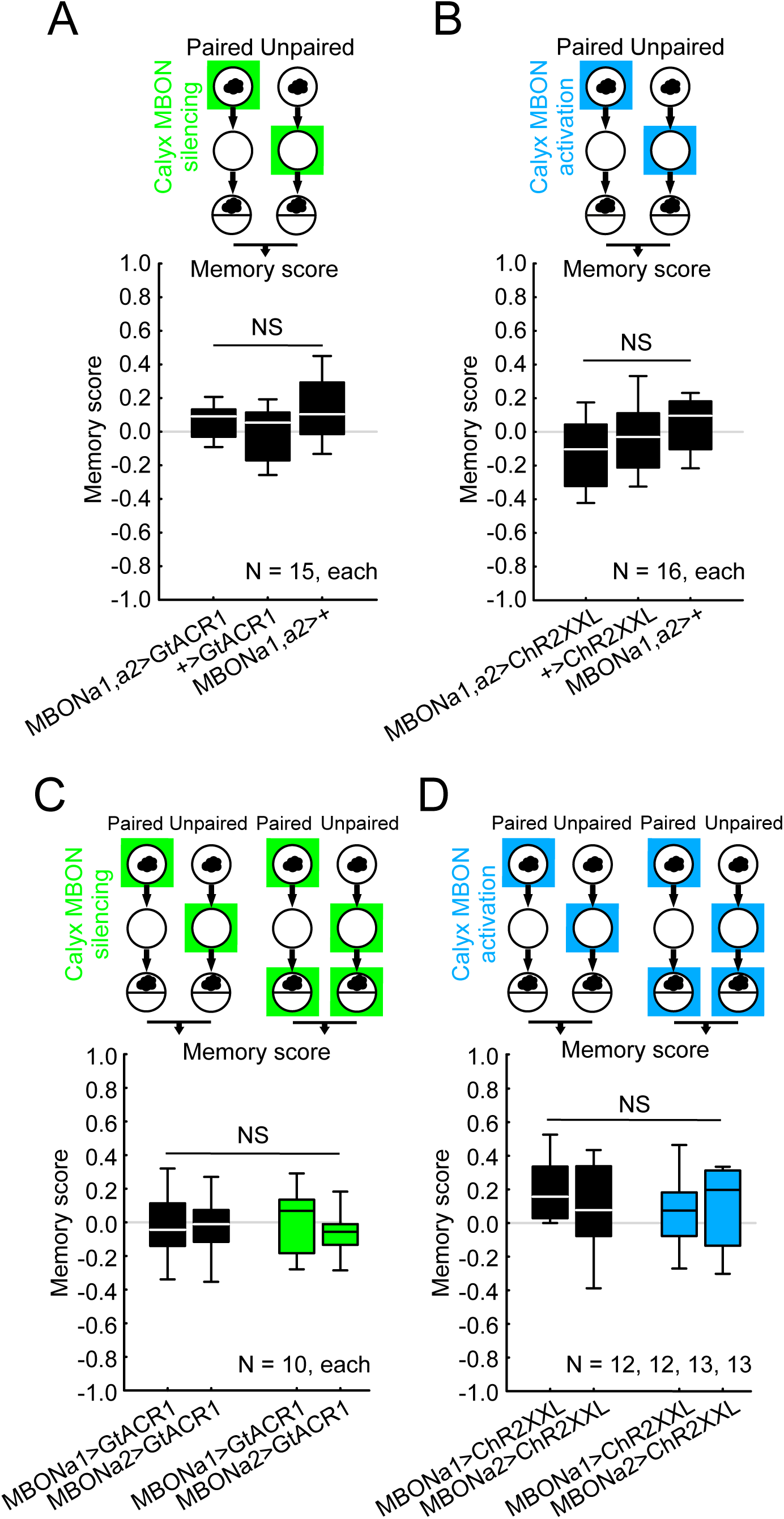
Manipulating activity in the calyx MBONs has no reinforcing effect. **(A)** Larvae were trained such that odor was presented either paired or unpaired with the silencing of the two calyx MBONs, using GtACR1 as the effector and green light illumination (green square). Silencing the two calyx MBONs together was seen to have no rewarding effect, as larvae of the experimental genotype (MBONa1,a2>GtACR1) did not behave differently from the genetic controls. **(B)** Activating the two calyx MBONs together likewise had no rewarding effect. **(C-D)** Silencing (C) or activating (D) the two calyx MBONs separately had no reinforcing effect, either. The sample sizes and the genotypes are given within the figure. NS refers to the absence of significance between groups in MWW comparisons (NS p> 0.05). The source data and results of all statistical tests are documented in Extended Data Figure 3-1. Other details as in Figures 9-15. Expression patterns of the calyx MBON drivers used in (C-D) are shown in Extended Data Figure 16-1.

Given the role of dopamine in conveying reward signals in larval *Drosophila* (Selcho et al., 2009; Rohwedder et al., 2016; Thoener et al., 2020), we next inquired into the dopamine- dependency of APL’s rewarding effect.

### Inhibition of dopamine synthesis impairs odor-APL memory

We used a systemic pharmacological approach to acutely disrupt dopamine-synthesis. This was done by inhibiting the enzyme tyrosine hydroxylase (TH), which is rate-limiting for dopamine synthesis (Neckameyer, 1996; Bainton et al., 2000; Fernandez et al., 2017; Thoener et al., 2020). The TH-inhibitor 3IY was added to the larval food at a dose which leaves intact task-relevant behavioral faculties (i.e. innate odor preference and locomotion) (Thoener et al., 2020). When 4 h later the larvae were trained and tested for odor-APL memory, they exhibited reduced memory scores (Figure 17A-C). In a repetition of this experiment, we showed that this reduction in memory was rescued in larvae that were additionally fed with the dopamine precursor L-DOPA (Figure 17D; notably, L-DOPA alone did not increase memory scores: Figure 17D). These results suggest that the rewarding effect of APL activation involves a dopaminergic process.

**Figure 17.**
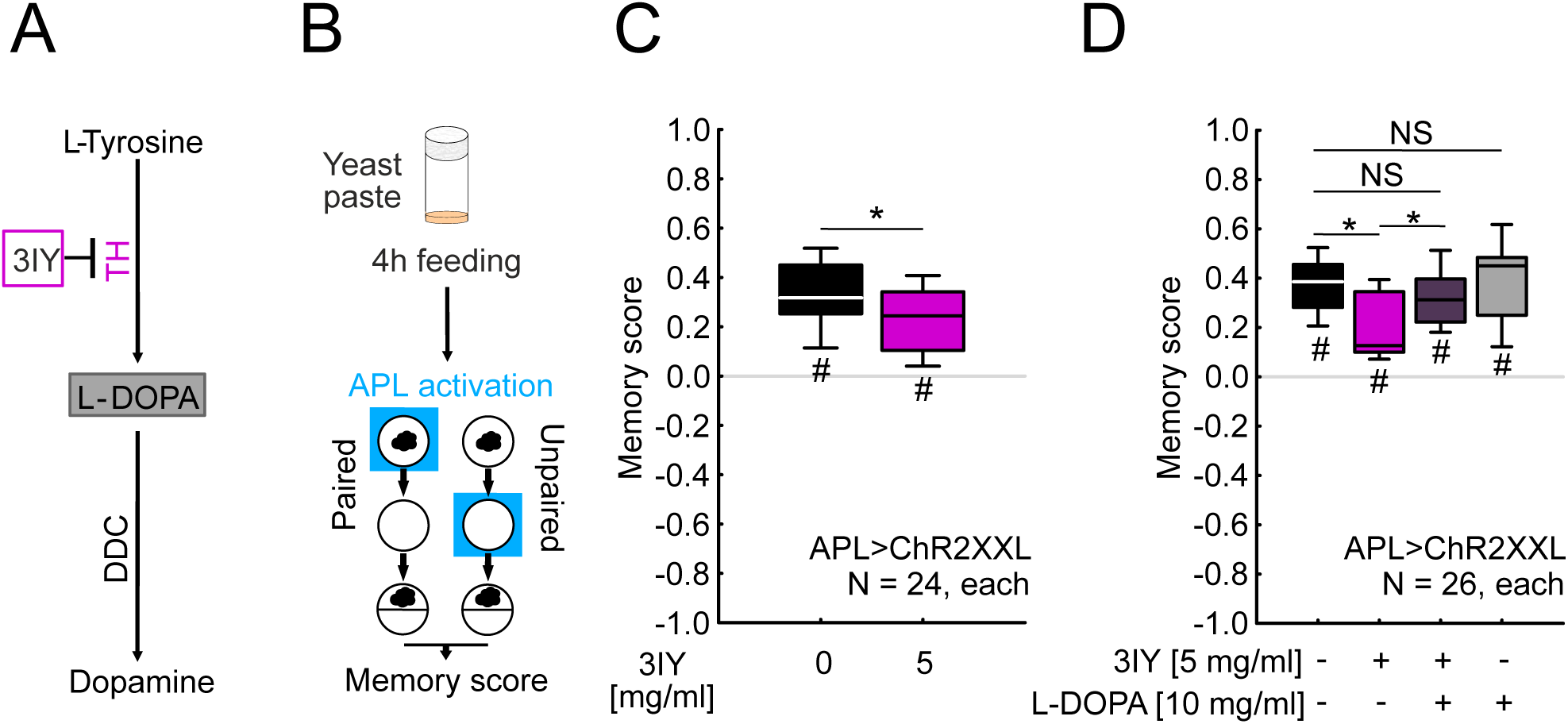
Inhibition of dopamine synthesis impairs odor-APL memory. A systemic pharmacological approach was used to disrupt dopamine synthesis (Thoener et al., 2020). **(A)** Sketch of dopamine biosynthesis. The enzyme tyrosine hydroxylase (TH) converts the amino acid L-tyrosine to L-3,4- dihydroxyphenylalanine (L-DOPA). In the next step the enzyme dopa-decarboxylase (DDC) converts L-DOPA to dopamine. Application of 3- Iodo-L-tyrosine (3IY) inhibits the TH enzyme. **(B)** Third-instar APL>ChR2XXL larvae were transferred from their food vials to a yeast solution either without 3IY or supplemented with 3IY. After 4 h of such feeding, the animals were trained and tested as in Figure 12A. **(C)** Relative to control larvae the 3IY-fed larvae exhibited impaired odor-APL memory scores. **(D)** As in (C) except that the yeast solution was prepared either (i) without additional substances, (ii) with 3IY added, (iii) with 3IY plus L-DOPA (a dopamine precursor), or (iv) with L-DOPA only, at the concentrations indicated. Again, relative to control larvae, reduced memory scores were observed in 3IY-fed larvae; feeding them additionally with L-DOPA rescued that memory impairment, leading to scores similar to those of the control animals. L-DOPA alone had no impact on odor-APL memory. The sample sizes and the genotypes are given within the figure. In (C-D), # refers to OSS comparisons to chance levels (i.e. to zero) with a Bonferroni-Holm correction (# p< 0.05); * refers to significant differences between groups in MWW comparisons with a Bonferroni-Holm correction (* p< 0.05). The source data and results of all statistical tests are documented in Extended Data Figure 3-1. Other details as in Figures 9-16.

### Ablation of the dopaminergic pPAM neurons impairs odor-APL memory

We next sought to identify the dopaminergic neurons that mediate the rewarding effect of APL activation. Given the role of the dopaminergic pPAM neurons in larval reward learning (Rohwedder et al., 2016; Saumweber et al., 2018; Schleyer et al., 2020; Thoener et al., 2022), we combined the GAL4/UAS with the lexA/lexAop systems to optogenetically activate APL (APL-GAL4>UAS-ChR2XXL) in animals expressing the pro-apoptotic *reaper* gene in the pPAM neurons, leading to their ablation (pPAM-lexA>lexAop-reaper). Larvae of the experimental group (APL_55D08_>ChR2XXL; pPAM>reaper) showed reduced odor-APL memory scores relative to genetic controls that lacked either the reaper effector or the pPAM driver for ablation (APL_55D08_>ChR2XXL; pPAM>+ and APL_55D08_>ChR2XXL; +>reaper, respectively) (Figure 18). Taken together, these results suggest that the rewarding effect of the activation of APL comes about, in part, by engaging a dopaminergic and pPAM-dependent process.

**Figure 18.**
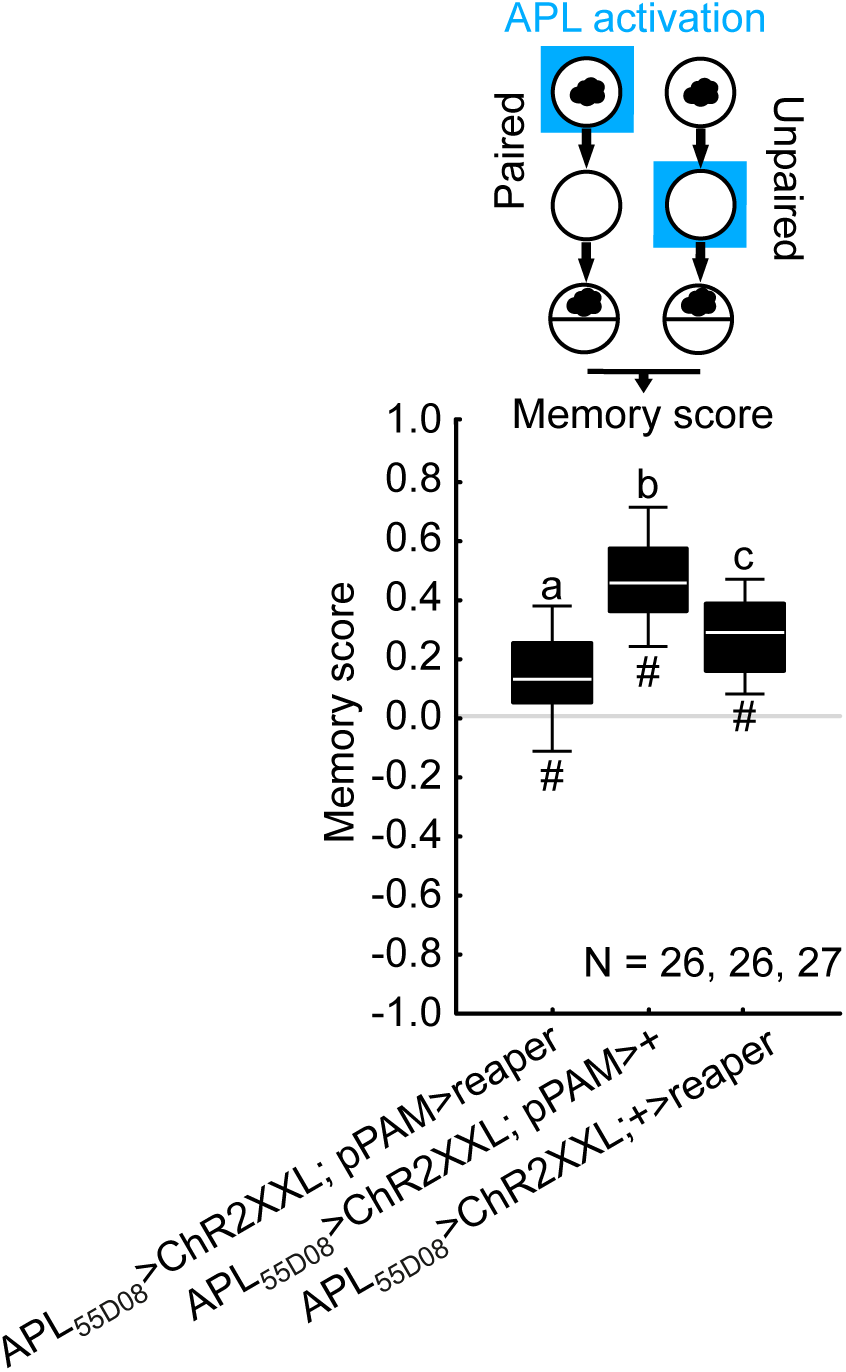
Ablation of dopaminergic pPAM neurons impairs odor-APL memory. Larvae were trained and tested as in Figure 12A. Optogenetic activation of APL and simultaneous ablation of the pPAM neurons (APL_55D08_>ChR2XXL; pPAM>reaper) reduced odor-APL memory scores relative to genetic controls (APL_55D08_>ChR2XXL; pPAM>+ and APL_55D08_>ChR2XXL; +>reaper). The sample sizes and the genotypes are given within the figure. Different letters refer to significant differences between groups in MWW comparisons with a Bonferroni-Holm correction (p< 0.05). # refers to OSS comparisons to chance levels (i.e. to zero), also with a Bonferroni-Holm correction (# p< 0.05). The source data and results of all statistical tests are documented in Extended Data Figure 3-1. Other details as in Figures 9-17.

## Discussion

The APL neuron is among the most complex neurons in the brain of both larvae and adults. The present study consolidates and broadens our knowledge of the morphology of the larval APL, of its GABAergic nature and its capacity to inhibit mushroom body KCs, of the polarity and topology of its chemical synapses, its development through metamorphosis, and of the exquisite specificity of a transgenic driver strain for studying it (Figures 2-8; Masuda- Nakagawa et al., 2014; Eichler et al., 2017; Saumweber et al., 2018). All these findings are consistent with APL playing a role in the sparsening of neuronal activity across the mushroom body (Masuda-Nakagawa et al., 2014; adults: Lin et al., 2014; Amin et al., 2020) and establish APL as the most comprehensively described neuron in the larval mushroom body. The present study further uncovers unexpected functional complexity by revealing a rewarding effect of optogenetically activating APL (Figure 12A). Our experiments were then designed to ascertain key features of this effect and investigate how it comes about (Figures 12C-18).

### Features of the rewarding effect of APL activation

Optogenetic activation of APL, using either ChR2XXL or Chrimson as the effector, induces appetitive memory for an associated odor after only one training trial (Figures 12-15). This is similar to odor-sugar associative learning and to the rewarding effect of activating DAN-i1 (Weiglein et al., 2019; Thoener et al., 2022). Associative learning by APL activation has a symmetrical ‘temporal fingerprint’, with a fairly narrow temporal window for presenting the odor (Figure 12D-E). This mirrors what was found for DAN-d1 in the aversive domain (Weiglein et al., 2021), adding to the heterogeneity of internal reinforcement signals (Saumweber et al., 2018; Weiglein et al., 2021; Thoener et al., 2022; adults: Aso and Rubin, 2019; König et al., 2018; Handler et al., 2019).

The appetitive memories established by activating APL decay within a few minutes (Figure 12C), as do memories for odor-fructose, odor-DAN-i1, and odor-KC association (Neuser et al., 2005; Kleber et al., 2016; Lyutova et al., 2019; Weiglein et al., 2019; Thoener et al., 2022). Similar to odor-fructose memories and odor-DAN-i1 memories (Paisios et al., 2017; Schleyer et al., 2020), APL-induced memories are expressed as modulations of turning, but not of run speed (Figure 13).

In addition to establishing appetitive associative memory during training, the activation of APL can also terminate learned search behavior during the test (Figure 14A-B, Figure 15C). This is in line with reports on taste rewards (Schleyer et al., 2015a) and on activation of DAN- i1 as a reward (Schleyer et al., 2020). Likewise similar to taste rewards and DAN-i1 activation, activating APL does not affect task-relevant innate olfactory behavior (Figure 14C-E, Figure 15D).

The rewarding effect of the activation of APL offers a new perspective on the result from Saumweber et al. (2018) showing that activating APL *throughout the complete training phase* abolishes odor-fructose memory scores, an effect that we replicated (Figure 9A-B). In these experiments, a fructose reward is presented paired with or unpaired from the odor, and the protocol of APL activation engages an additional, indeed much stronger reward signal *throughout training* and thus *regardless of the presence or absence of the odor*, thus overriding the learning of the relationship between odor and fructose.

Thus, the rewarding effect of activating APL is compatible with the previous results from Saumweber et al. (2018), and the resulting appetitive, associative memories do not seem at odds with those established by DANs or taste reinforcement.

### Possible mechanisms of odor-APL learning: notes upfront

Our results confirm that the larval APL neuron is GABAergic (Figure 4A-B’’; Masuda- Nakagawa et al., 2014). Regarding octopamine, acetylcholine and glutamate, negative results have been reported previously (Masuda-Nakagawa et al., 2014; Eichler et al., 2017). This matches a recent transcriptome analysis in adults (Aso et al., 2019), whereas two earlier reports had suggested the presence of GABA, octopamine and glutamate in the adult APL (Wu et al., 2013; Li et al., 2017). In the absence of evidence suggesting otherwise for the larval APL, however, the following discussion considers only GABAergic signaling.

GABA binding to ionotropic GABA-A receptors and the ensuing chloride influx confer the typical inhibitory effect of GABA on post-synaptic neurons. Accordingly, APL activation reduces activity in KCs (Figure 5E-F). However, additional effects of GABA via metabotropic receptors may be reckoned with. In motoneurons of pupal *Drosophila*, moreover, GABA- induced spikelets have been observed, arguably because of a relatively positive reversal potential for chloride during this life stage (Ryglewski et al., 2017; mammals: Ben-Ari, 2002). While it thus cannot be ruled out that GABA release from APL leads to excitatory effects in a minority of KCs or in non-KC targets of APL, the discussion below maintains the conventional notion of GABA as a transmitter with inhibitory effect. However, it seems plausible that upon the offset of inhibition there may be post-inhibitory rebound activation in the target neurons, a widely observed physiological phenomenon (Huguenard and McCormick, 2007; for evidence in adult *Drosophila* after prolonged APL activation: Apostolopoulou and Lin, 2020).

The fact that activation of APL can establish appetitive memory for an associated odor means that even under conditions of GABAergic, inhibitory input an associable odor representation can be established across the KCs. Indeed, these odor representations can be specific enough to allow for differential conditioning (Figure 14B). The scenario could be that under baseline conditions odors activate ‘their’ subset of KCs relatively strongly while the other KCs are inactive or mildly inhibited; upon optogenetic activation of APL, however, odors only mildly activate an even sparser set of KCs while most other KCs would be relatively strongly inhibited. Interestingly, associable odor representations can be established also under conditions of optogenetically *in-*creased levels of activity across the KCs (Lyutova et al., 2019).

We are not aware of any data suggesting that GABA can have a direct associative memory- trace-inducing effect. Rather, our results suggest that the rewarding effect of APL activation comes about, at least in part, by engaging a dopaminergic reward signal from the pPAM neurons (Figure 17, Figure 18). We will therefore focus on plausible pathways from APL towards the dopaminergic pPAM neurons.

### From APL to dopaminergic pPAM neurons?

The larval APL hosts pre-synapses only in the calyx (Figure 4, Figures 6-8, **Movie 3**). The post-synaptic partners of APL include the KCs (see next paragraph) and the two calyx MBONs (Eichler et al., 2017; Saumweber et al., 2018). One of these calyx MBONs promotes approach when optogenetically activated (MBON-a1), whereas the other promotes avoidance (MBON- a2) (Eschbach et al., 2021). Both calyx MBONs give rise to indirect feedback to DANs, including to the punishing DAN-d1 and the rewarding DAN-i1 neuron of the pPAM cluster (Eschbach et al., 2020). However, presenting an odor together with activating or silencing the two calyx MBONs, alone or in combination, did not establish either appetitive or aversive associative odor memory (Figure 16). Thus, although it is a connectomic possibility, there is no evidence for a reinforcing APL-MBONa1/a2-pPAM loop. What about a loop from APL via the KCs to the pPAM neurons?

As argued above, odors presented under conditions of APL activation will only mildly activate a rather sparse, odor-specific set of KCs, while the great majority of the KCs will be strongly inhibited, directly by APL and possibly by lateral inhibition from the activated KCs in addition (Manoim et al., 2022). As with the excitatory synapses from KCs to DANs (Lyutova et al., 2019; adults: Cervantes-Sandoval et al., 2017), this would provide little, if any, drive for the dopaminergic pPAM neurons. Once the inhibition is lifted, however, post-inhibitory rebound activation from a large number of previously inhibited KCs might provide a volley of activity sufficient to activate DANs (Apostolopoulou and Lin, 2020). Indeed, the optogenetic activation of broad sets of KCs can activate dopaminergic pPAM neurons and exert a rewarding effect (Lyutova et al., 2019). Thus, such an APL-KC-pPAM loop could account for the rewarding effects of APL activation. Notably, punishing DANs also receive input from KCs (Eichler et al., 2017; Eschbach et al., 2020). In addition to the appetitive memories established by activation of KCs (Lyutova et al., 2019) or APL, therefore, it seems possible that an aversive memory is established, too, in the compartments innervated by these punishing DANs. Under the test conditions used by us and by Lyutova et al. (2019), however, the observed result of these manipulations is appetitive memory. This would suggest either that the appetitive memory is stronger than the aversive, or that under these test conditions the aversive memory remains behaviorally silent (Gerber and Hendel, 2006; Schleyer et al., 2011, Schleyer et al., 2015a).

An alternative could be that, similar to the situation in adults for APL and the DPM neuron (Wu et al., 2011), there is direct signaling from APL to rewarding pPAMs rather than to punishing DANs via electrical synapses in the lobes (Figures 3-4, Figures 6-7).

In summary, we report a case of complex circuit function in a numerically simple brain, and demonstrate the capacity of a central-brain GABAergic neuron to engage dopaminergic reward signaling when optogenetically activated.

## Acknowledgements

This study received institutional support from the Otto von Guericke University Magdeburg (OvGU), the Wissenschaftsgemeinschaft Gottfried Wilhelm Leibniz (WGL), the Leibniz Institute for Neurobiology (LIN), as well as grant support from the Deutsche Forschungsgemeinschaft (DFG-GE 1091/4-1 and FOR 2705 Mushroom body to B.G.; PA1979/3-1 to D.P.; DFG-TH 1584/ 8-1 and DFG-TH 1584/ 7-1 to A.S.T.; NIH Grant R01 NS054814 to V.H.). We thank A. Bates (Cambridge) for providing the script allowing compartment boundaries to be generated on the dendrograms; H. Tanimoto (U Tohoku), M. Stopfer (NIH), Y. Aso (HHMI Janelia), A. Lin (U Sheffield), O. Schuldiner (Weizmann), C. Duch (U Mainz), G. Tavosanis (U Bonn), A. Khalili (U Göttingen), C. König (LIN), B. Michels (LIN) and N. Toshima (LIN) for discussions; A. Ciuraszkiewicz (LIN), K. Hartung (LIN), M.L. Rodriguez Lopez (LIN), B. Kracht (LIN), T. Niewalda (LIN), M. Paisios (LIN), H. Reim (LIN) and H. Wickborn (LIN) and the workshop team at the LIN for their technical assistance. Current affiliations: N.M.: Max Planck Florida Institute for Neuroscience, 33458 Jupiter, Florida, USA; A.W.: Institute of Anatomy, Otto von Guericke University, 39120 Magdeburg, Germany; O.M.: Biozentrum, University of Basel, 4056 Basel, Switzerland; M. Schleyer: Hokkaido University, Institute for the Advancement of Higher Education, Hokkaido 060-0808, Japan; A.C.: Department of Zoology, University of Cambridge, Cambridge, CB2 3EJ, United Kingdom and MRC Laboratory of Molecular Biology, Cambridge, CB2 0QH, United Kingdom; K.E.: Department of Zoology, University of Cambridge, Cambridge, CB2 3EJ, United Kingdom.

## Extended Data

**Extended Data Table 2-1.** Word file table containing the key reagents (fly strains, antibodies, software) used in this study.

**Extended Data Figure 3-1**. Excel file containing the source data presented in Figures 3, 5, 6, 7, 9-18 along with all statistical results. The data for each figure are presented in a separate sheet, and the statistical results are grouped by figure number, figure panel, and statistical tests. As regards the behavioral results displayed in Figures 9-18, data are grouped by figure panel, genotype, food supplementation, and test condition; each sheet includes the odor preference values underlying the memory scores shown in the main figures.

**Extended Data** Figure 7-1. **High-resolution dendrograms of APL.** The colored envelopes indicate the mushroom body compartments innervated by the left- and right-hemisphere APL. Other details as in Figure 7A.

**Extended Data** Figure 7-2. **High-resolution dendrograms of APL showing sites of synapses with mushroom body extrinsic neurons.** Synapses are shown at their topologically correct site on APL from both hemispheres with the mushroom body extrinsic neurons indicated. Pre- and post-synaptic sites of APL are annotated with dots and triangles, respectively. Other details as in Figure 7B.

Extended Data Figure 7-3. High-resolution dendrograms of APL showing sites of synapses with mushroom body intrinsic Kenyon cells. Synapses with the Kenyon cells (KCs) are shown at their topologically correct site on APL from both hemispheres. Dark purple dots and bright purple triangles show APL-to-KC and KC-to-APL synapses, respectively. Other details as in Figure 7C.

**Extended Data** Figure 7-4. **Cluster analysis of APL calycal synaptic sites with the KCs.** Cluster analysis showing that calycal synaptic sites of the left- and right-hemisphere APL with the KCs are organized in four clusters (1-4). Most of the APL-to-KC synapses (dark purple dots) are observed towards the center of these clusters (dark square), whereas KC-to-APL synapses (bright purple triangles) are observed mainly in the surround. Other details as in Figure 7D.

**Extended Data** Figure 16-1. **Expression patterns of the calyx MBON drivers used in** Figure 16C-D. **(A)** Full projection of the expression pattern from the SS02006-GAL4 driver (MBON-a1) covering only one calyx MBON in each hemisphere (N = 7 brains tested). **(B)** As in (A), but for SS01417-GAL4 (MBON-a2); in two out of N= 5 brains tested this driver covered both calyx MBONs (white arrowheads; left: cell bodies overlap one another). The data were acquired with a 63x glycerol objective; scale bar and grid spacing: 20 µm. Other details as in Figure 9C.

## Movie legends

**Mancini et al., Movie 1**. Volume rendering of fluorescence signals in a third-instar larva of the genotype APL>mCherry-CAAX at the indicated combinations of excitation and emission wavelengths (same specimen as in Figure 2A-B’’’). The movie starts in the brain region from a dorsal view (rostral to the left), and after rotation zooms out to show the full body. The data were acquired with a 12x objective; grid spacing: 200 µm.

**Mancini et al., Movie 2**. 3D rendering of the APL neuron (green) and the mushroom bodies (magenta) in a third-instar larval brain. Genotype: APL>mIFP/MB247>mCherry-CAAX. The data were acquired with a 40x oil objective; grid spacing: 20 μm.

**Mancini et al., Movie 3**. 3D rendering and segmentation of pre- (green) and post-synaptic (magenta) regions of the APL neuron in a third-instar larval brain, based on data shown in Figure 4C-C’’. Genotype: APL>Dsyd-1::GFP/DenMark. The data were acquired with a 63x glycerol objective; grid spacing: 20 μm.

**Mancini et al., Movie 4**. Volume reconstruction of the left- and the right-hemisphere APL neuron (green) in a first-instar larval brain (CNS; gray), with the focus on APL’s connectivity within the mushroom bodies (MB; magenta). “X” refers to synapses with any type of partner. The location of pre- and post-synaptic sites of both APLs is annotated with yellow spheres and red pyramids, respectively. A: anterior; D: dorsal; M: medial. Based on the dataset from Eichler et al. (2017). See also Figure 6B.

**Mancini et al., Movie 5**. Volume reconstruction of the left- and the right-hemisphere APL neuron in a first-instar larva and sites of pre- and post-synapses with different subclasses of mushroom body intrinsic neurons (Kenyon cells, KCs), namely single-claw, multi-claw, and young KCs. Pre- and post-synaptic sites of APL are annotated with spheres and pyramids, respectively. A: anterior; D: dorsal; M: medial. Based on the dataset from Eichler et al. (2017). See also Figure 6E.

**Mancini et al., Movie 6**. Volume reconstruction of the left- and the right-hemisphere APL neuron in a first-instar larva and sites of pre- and post-synapses with different types of partner, separated into all subclasses of mushroom body intrinsic neurons (Kenyon cells, KCs) and the mushroom body extrinsic neurons indicated. Remaining neurons bearing less than two synapses with APL in both hemispheres are shown as “Other”; “X” refers to synapses with non-KCs. Pre- and post-synaptic sites are annotated with spheres and pyramids, respectively. A: anterior; D: dorsal; M: medial. Based on the dataset from Eichler et al. (2017). See also Figure 6F.

**Mancini et al., Movie 7**. 3D rendering of the expression pattern of ChR2XXL in APL in a third- instar larva, based on data shown in Figure 9C. Genotype: APL>ChR2XXL. The data were acquired with a 63x glycerol objective; grid spacing: 20 μm.

