## Extended Data Figure 7-1 for "Rewarding capacity of optogenetically activating a giant GABAergic central-brain interneuron in larval *Drosophila*"

Mancini et al.

### Extended Data Figure 7-1

**High-resolution dendrograms of APL.** The colored envelopes indicate the mushroom body compartments innervated by the left- and right-hemisphere APL. Other details as in Figure 7A.

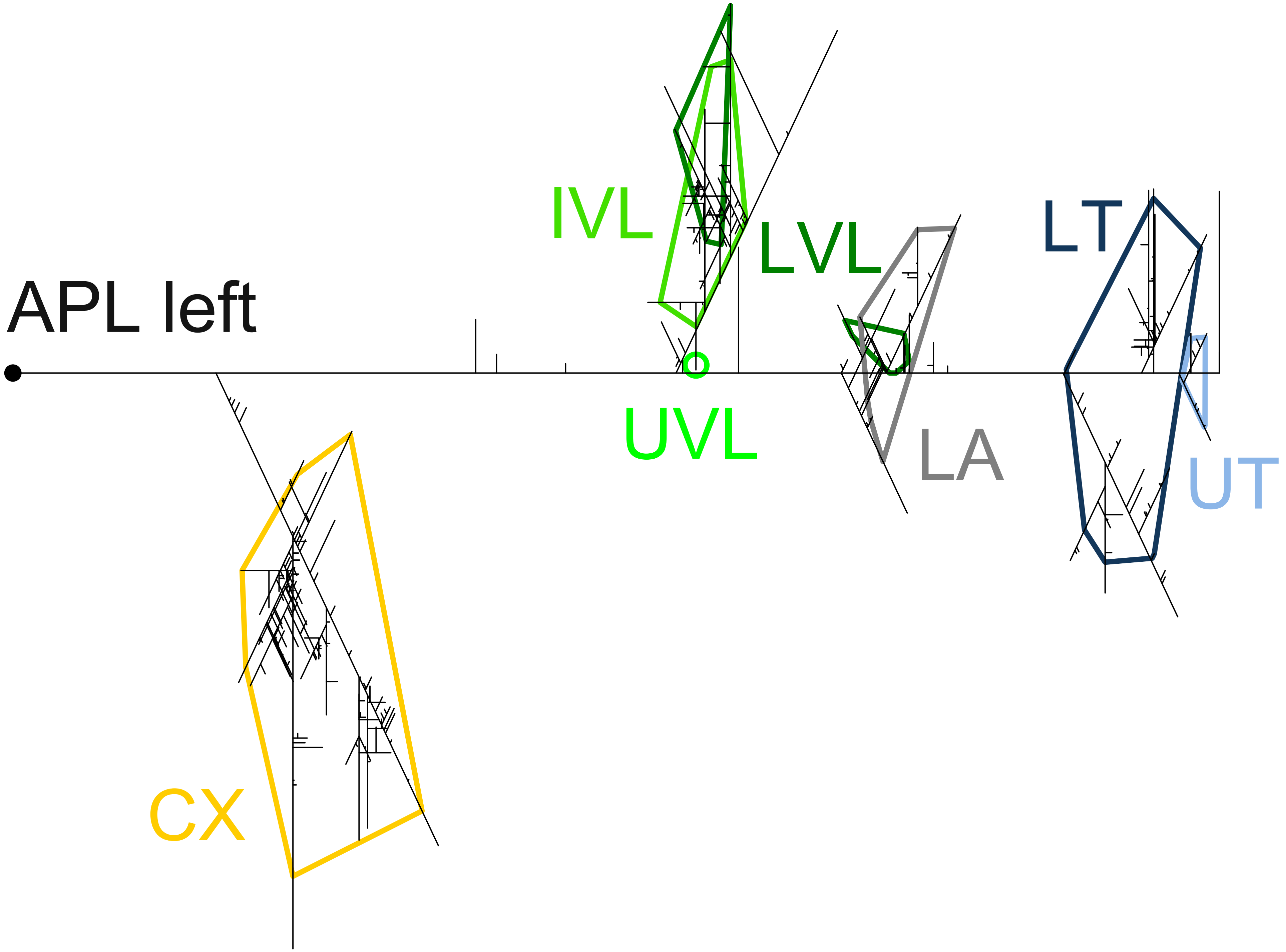
APL left
IVL
LVL
LT
UVL
LA
UT
CX

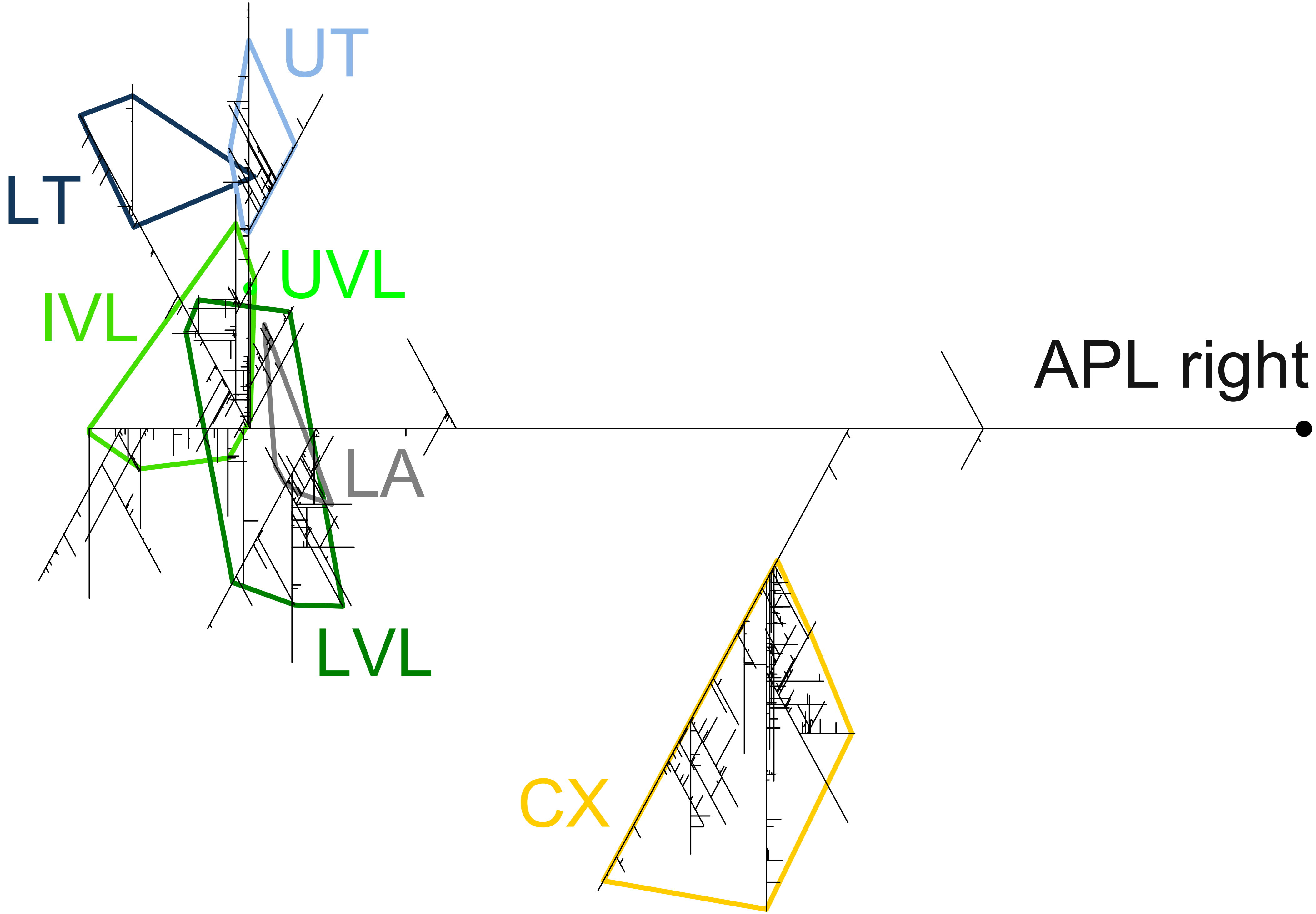
UT
LT
UVL
IVL
APL right
LA
LVL
CX
