## Extended Data Figure 7-2 for "Rewarding capacity of optogenetically activating a giant GABAergic central-brain interneuron in larval *Drosophila*"

Mancini et al.

### Extended Data Figure 7-2

**High-resolution dendrograms of APL showing sites of synapses with mushroom body extrinsic neurons.** Synapses are shown at their topologically correct site on APL from both hemispheres with the mushroom body extrinsic neurons indicated. Pre- and post-synaptic sites of APL are annotated with dots and triangles, respectively. Other details as in Figure 7B.

Color legend throughout:

- ▼ DAN-f1→APL
- ▼ DAN-i1→APL
- ▼ DAN-k1→APL
- ● APL→MBON-a1
- ● APL→MBON-a2
- ▼ MBONq1→APL
- ● APL→OAN-a1
- ● APL→OAN-a2
- ▼ OAN-a2→APL
- ● APL→Olfactory PNs
- ▼ Olfactory PNs→APL
- ● APL→Other
- ▼ Other→APL

#### APL left

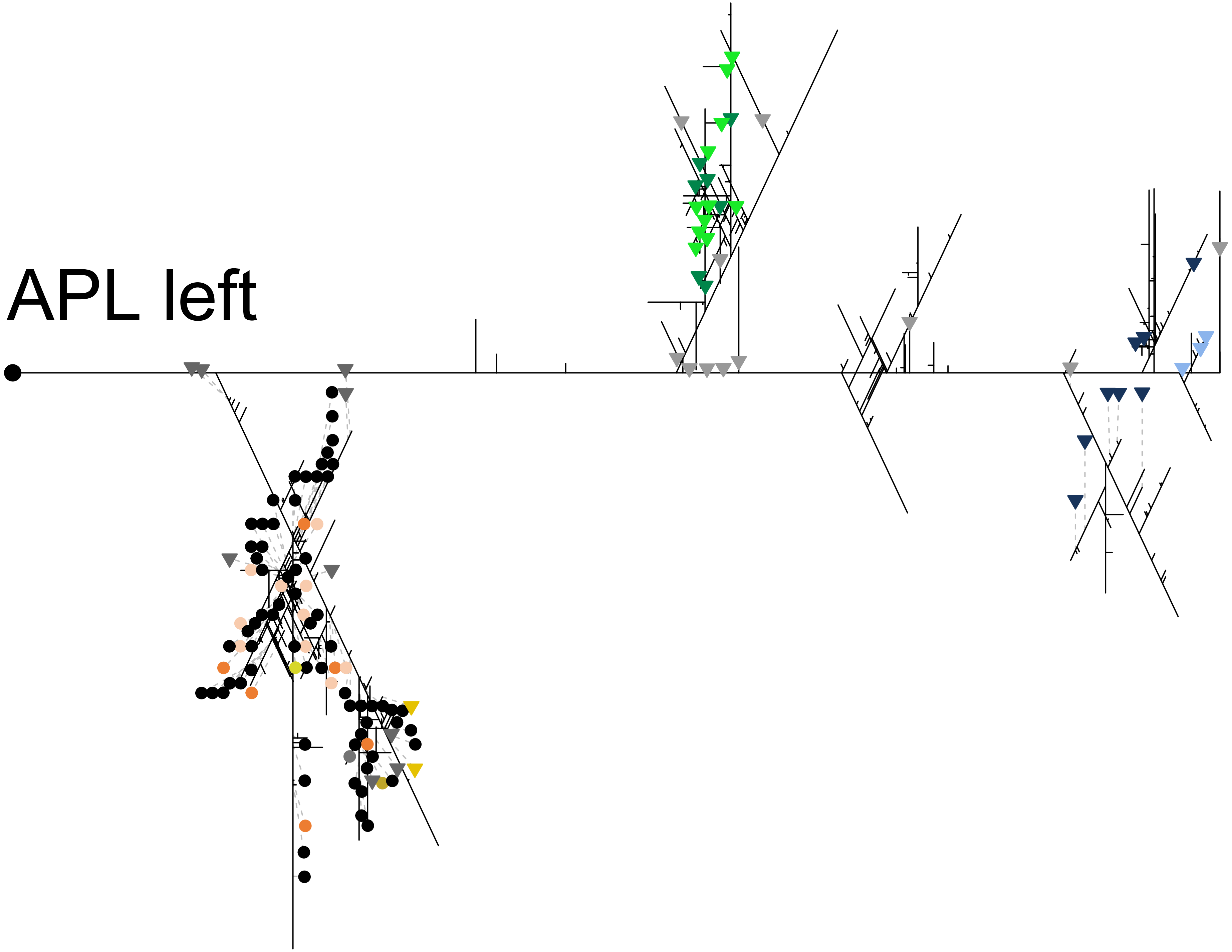

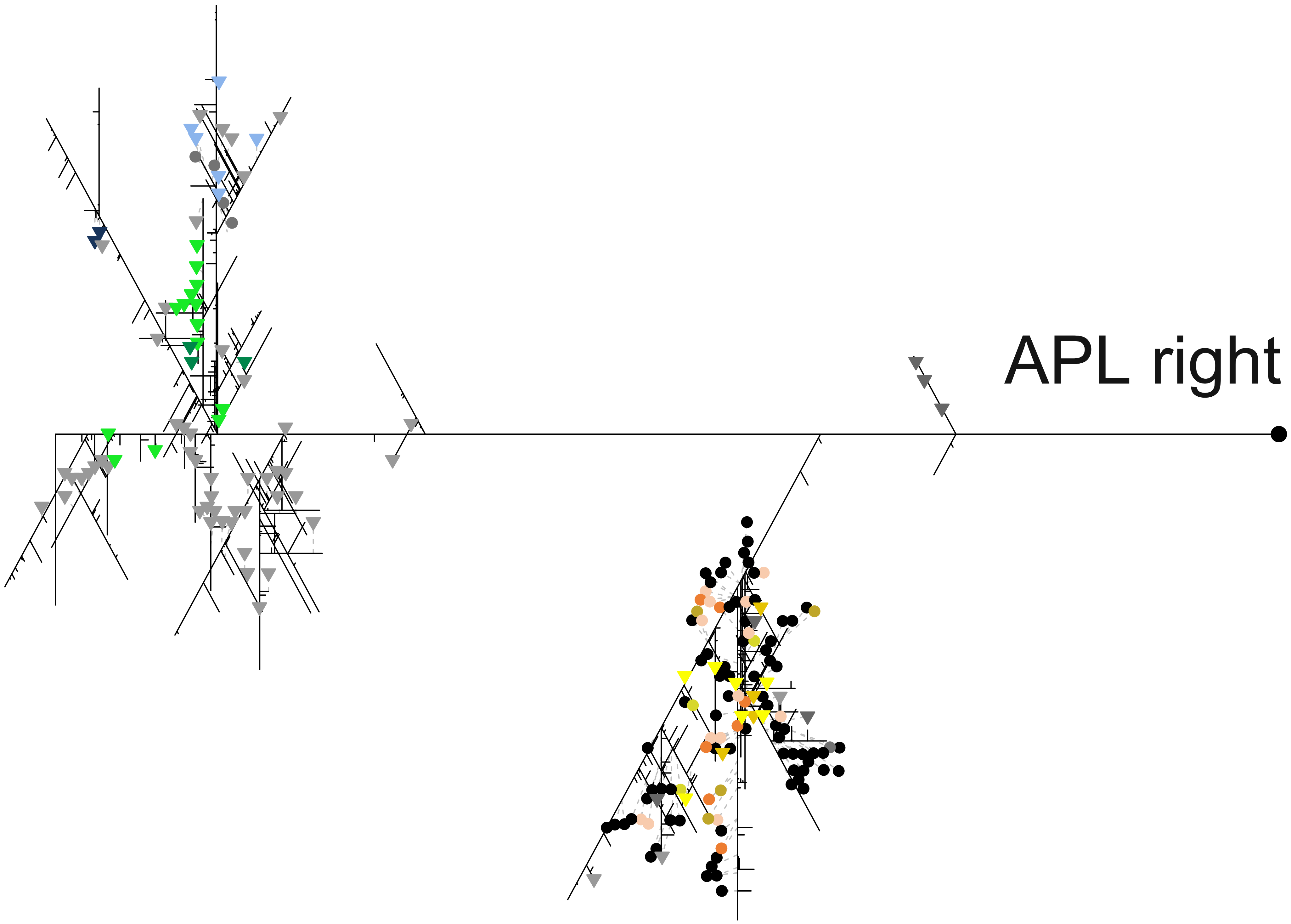
APL right
