## Extended Data Figure 7-3 for "Rewarding capacity of optogenetically activating a giant GABAergic central-brain interneuron in larval *Drosophila*"

Mancini et al.

### Extended Data Figure 7-3

**High-resolution dendrograms of APL showing sites of synapses with mushroom body intrinsic Kenyon cells.** Synapses with the Kenyon cells (KCs) are shown at their topologically correct site on APL from both hemispheres. Dark purple dots and bright purple triangles show APL-to-KC and KC-to-APL synapses, respectively. Other details as in Figure 7C.

Color legend throughout: ● APL→KC
▼ KC→APL
■ Cluster center

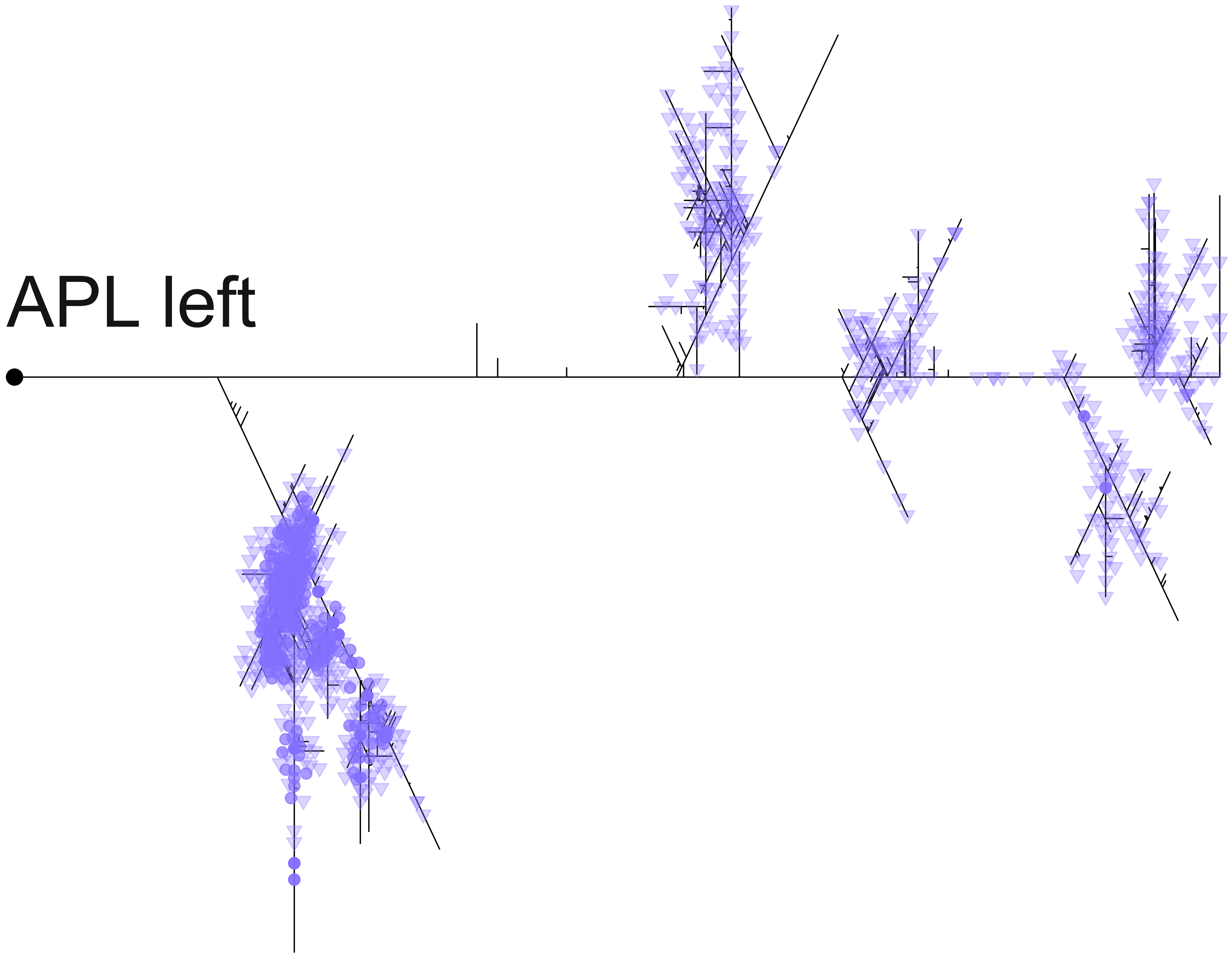
APL left

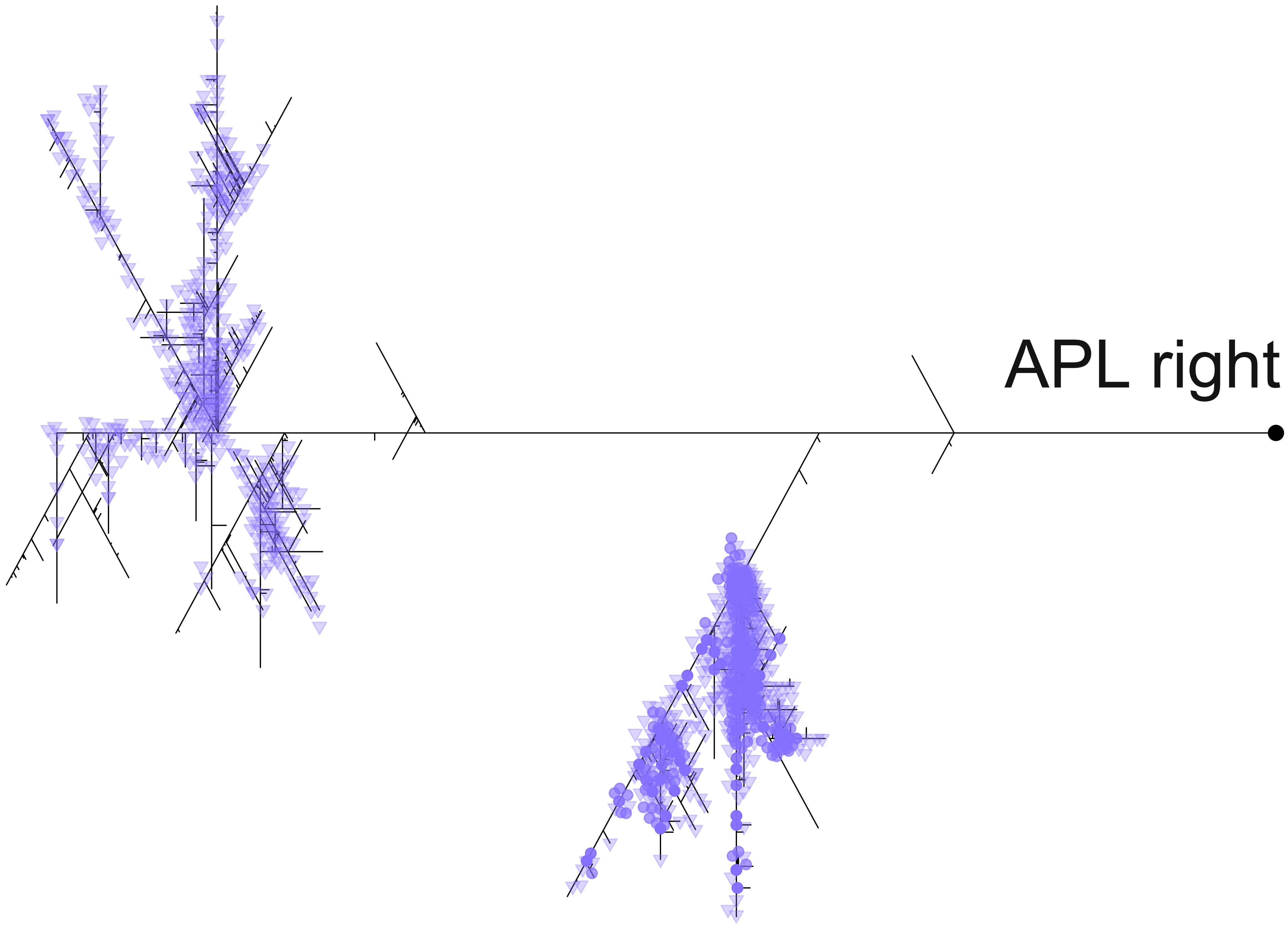
