## Extended Data Figure 7-4 for "Rewarding capacity of optogenetically activating a giant GABAergic central-brain interneuron in larval *Drosophila*"

Mancini et al.

### Extended Data Figure 7-4

**Cluster analysis of APL calycal synaptic sites with the KCs.** Cluster analysis showing that calycal synaptic sites of the left- and right-hemisphere APL with the KCs are organized in four clusters (1-4). Most of the APL-to-KC synapses (dark purple dots) are observed towards the center of these clusters (dark square), whereas KC-to-APL synapses (bright purple triangles) are observed mainly in the surround. Other details as in Figure 7D.

Color legend throughout: ● APL→KC

▼ KC→APL

■ Cluster center

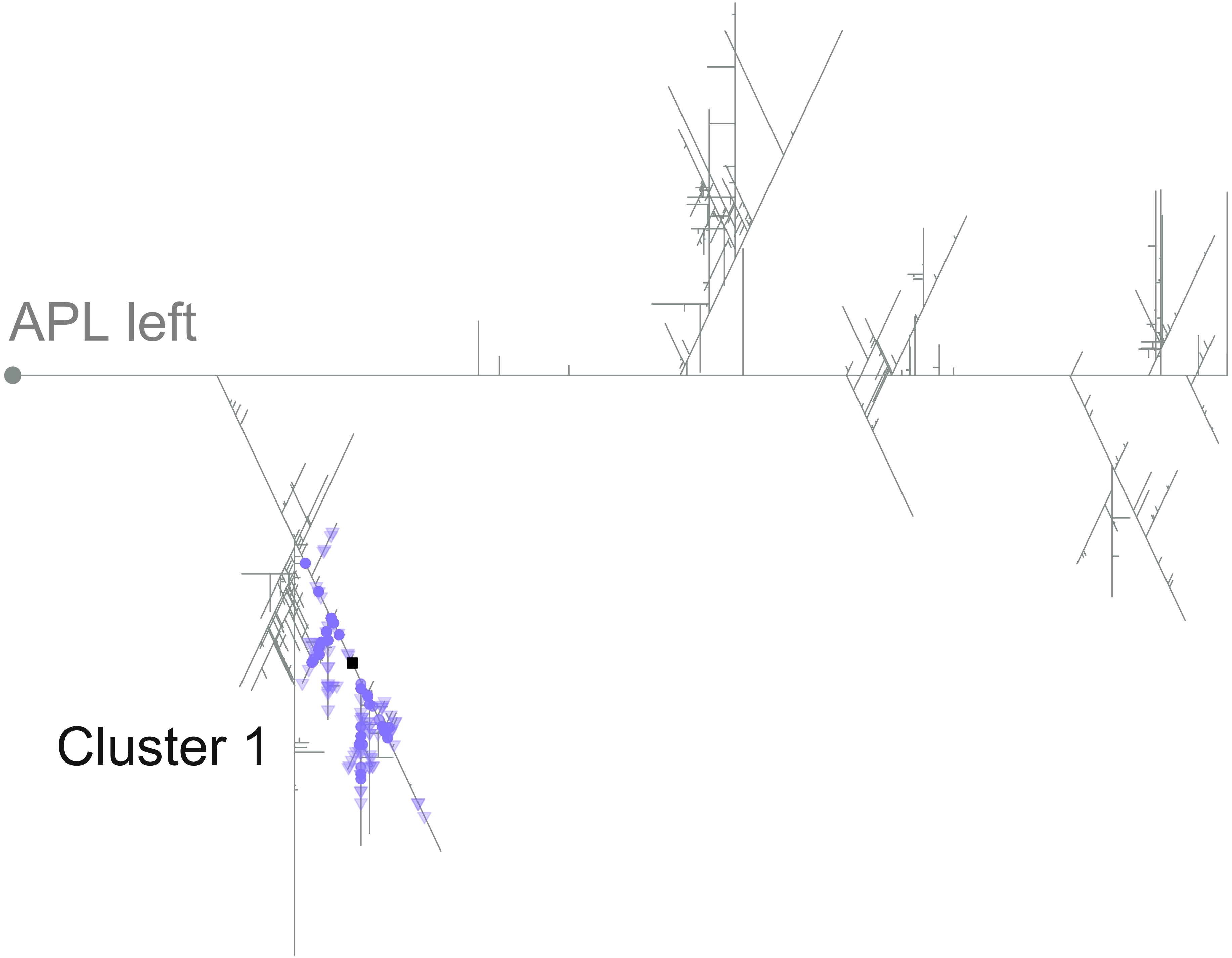
APL left
Cluster 1

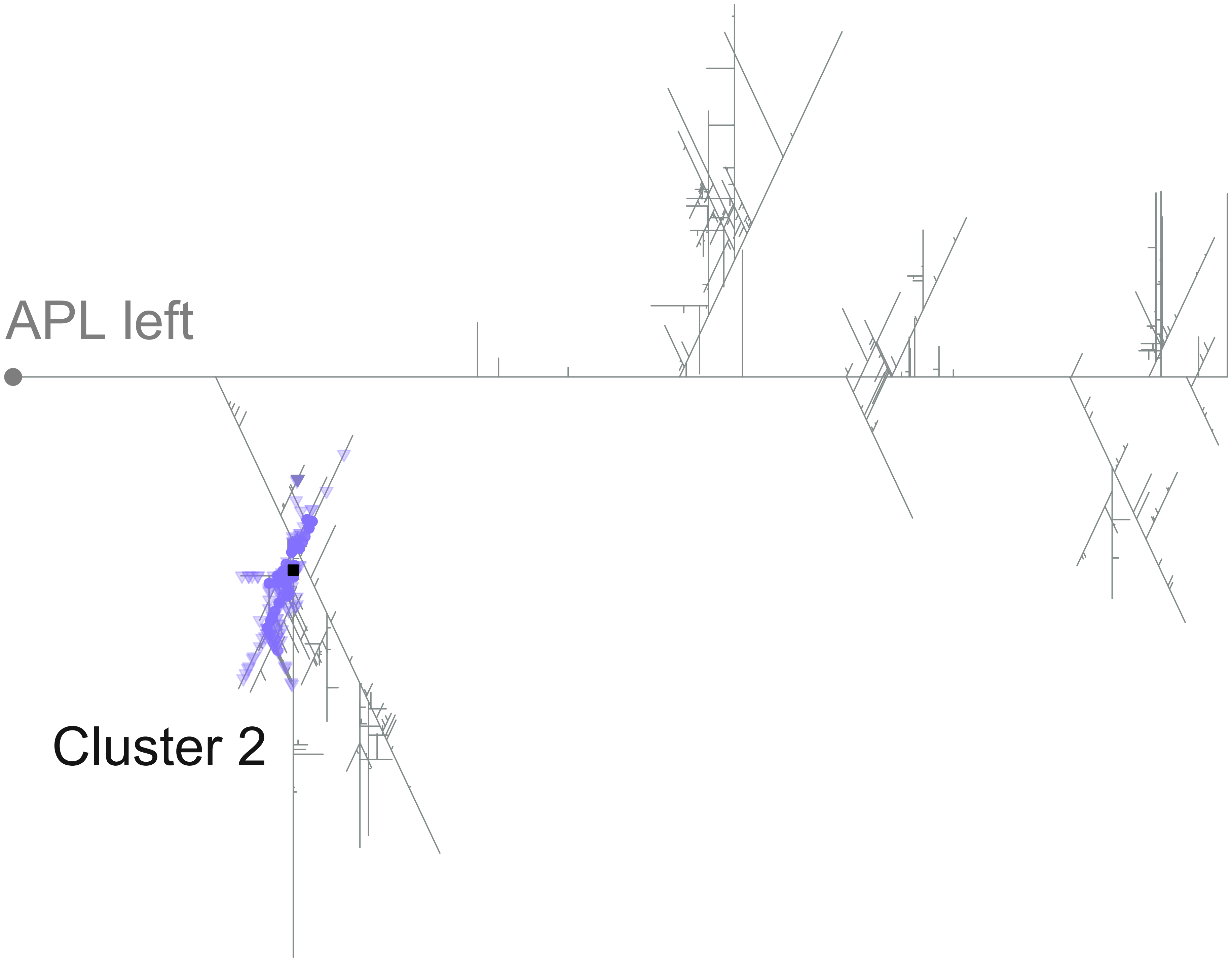
APL left
Cluster 2

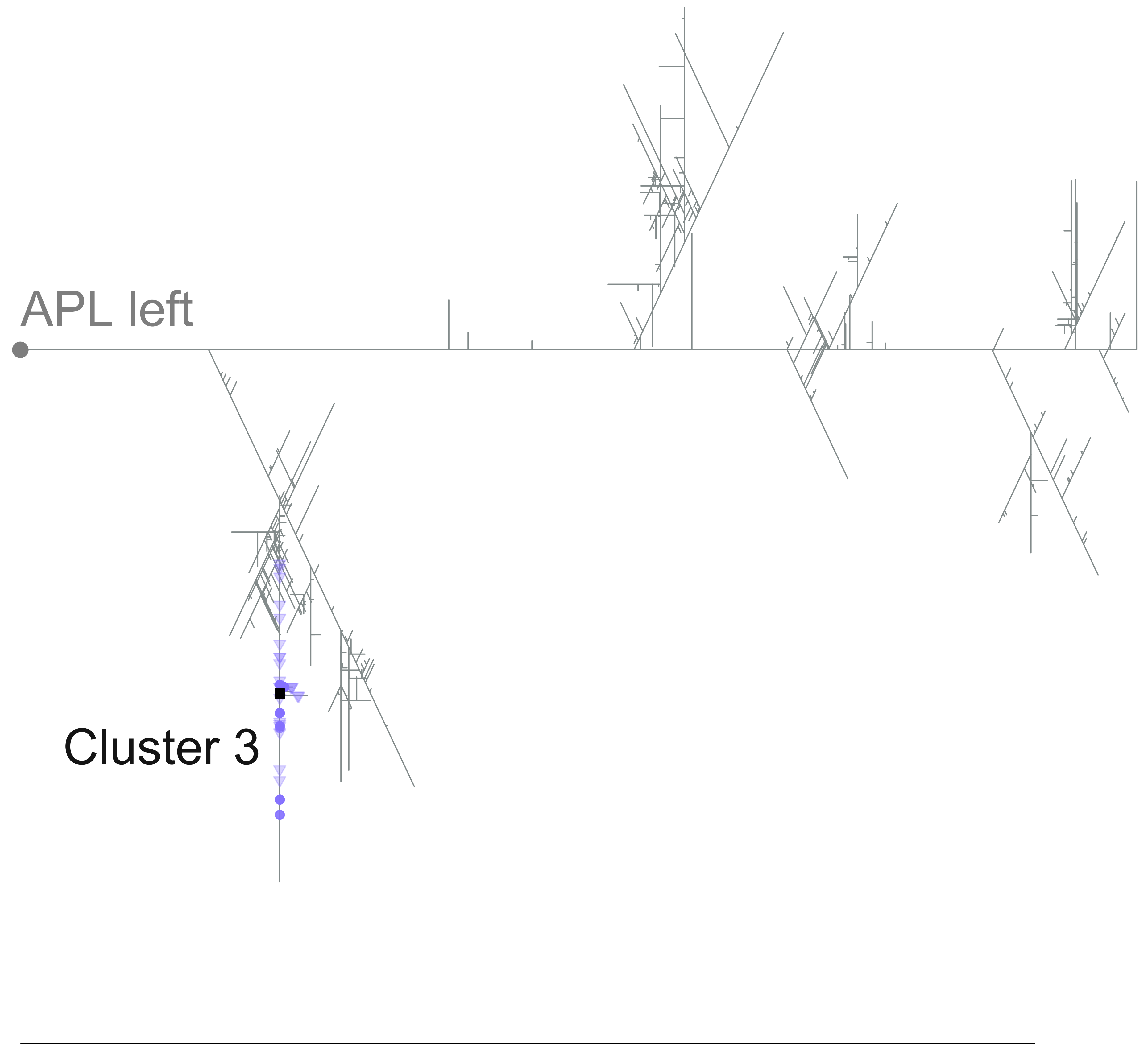
APL left
Cluster 3

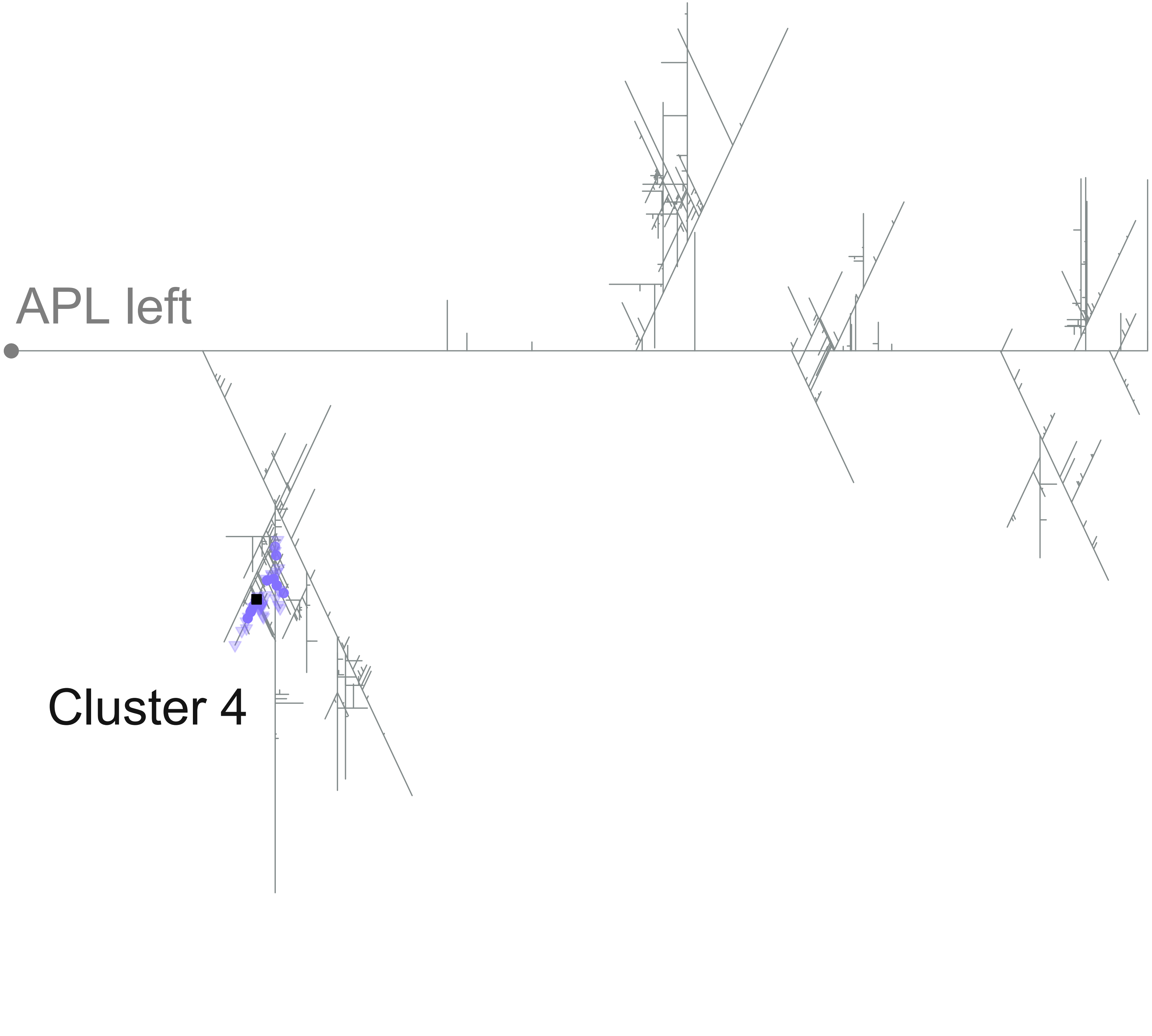
APL left
Cluster 4

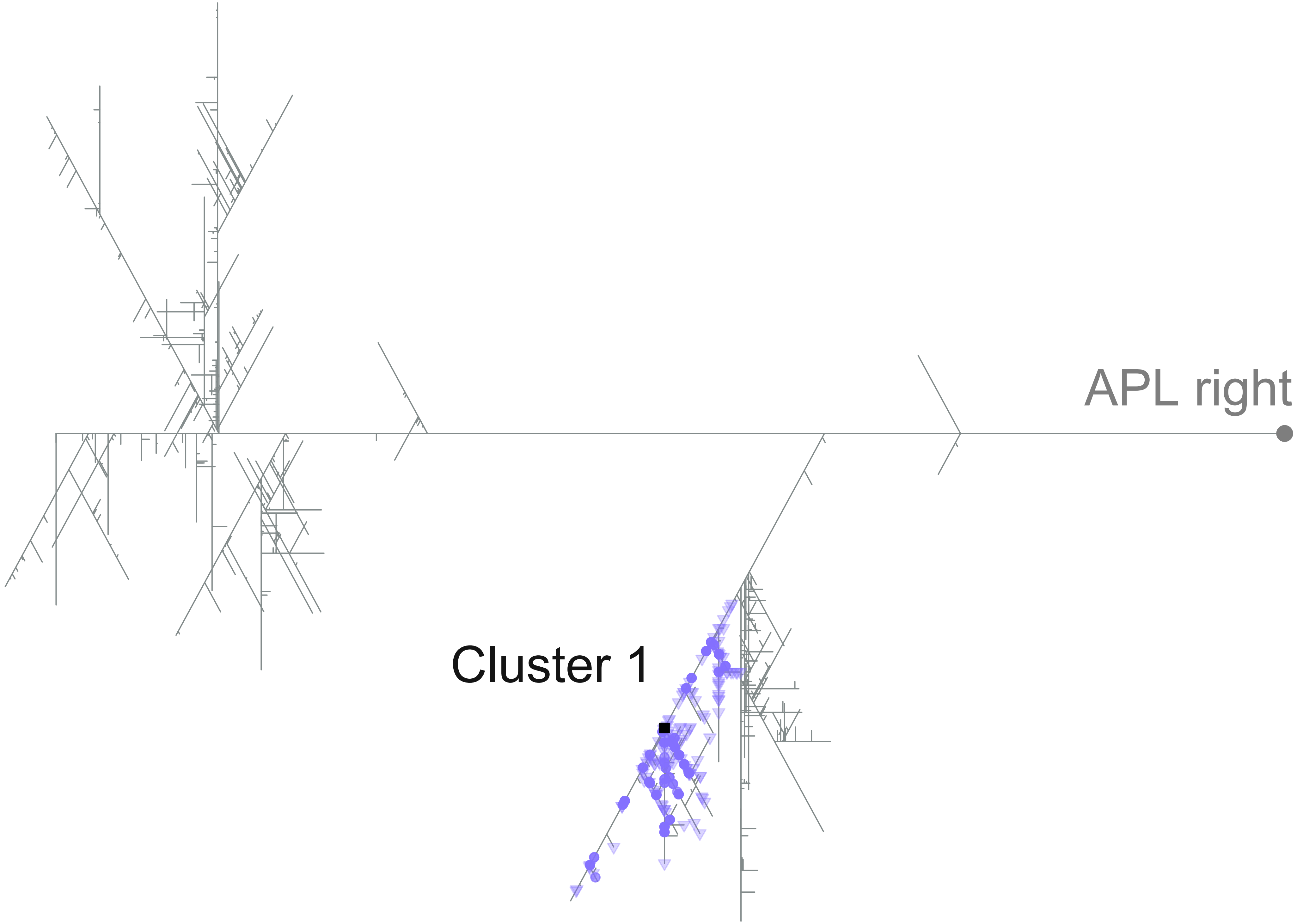
APL right
Cluster 1

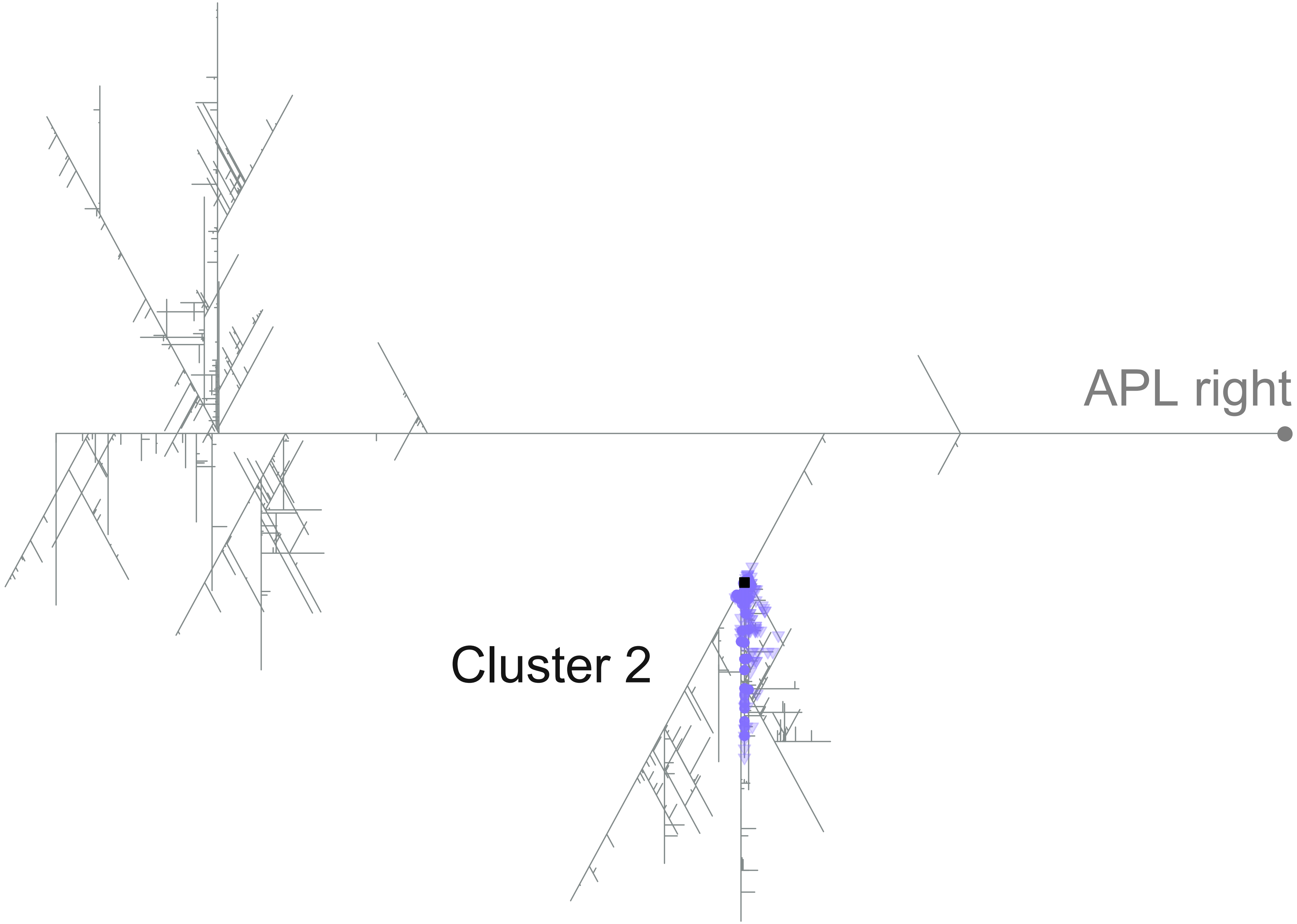
APL right
Cluster 2

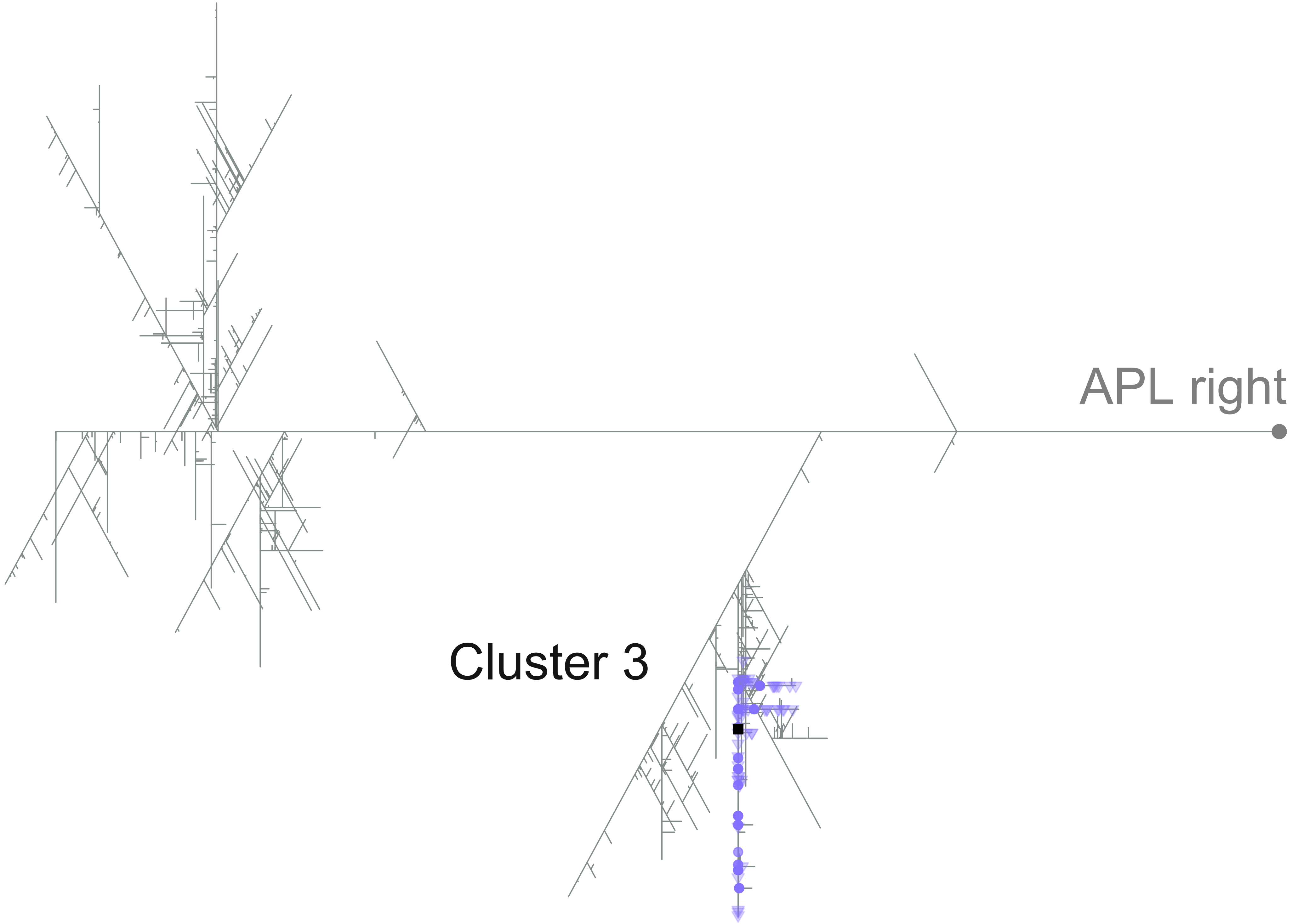
APL right
Cluster 3

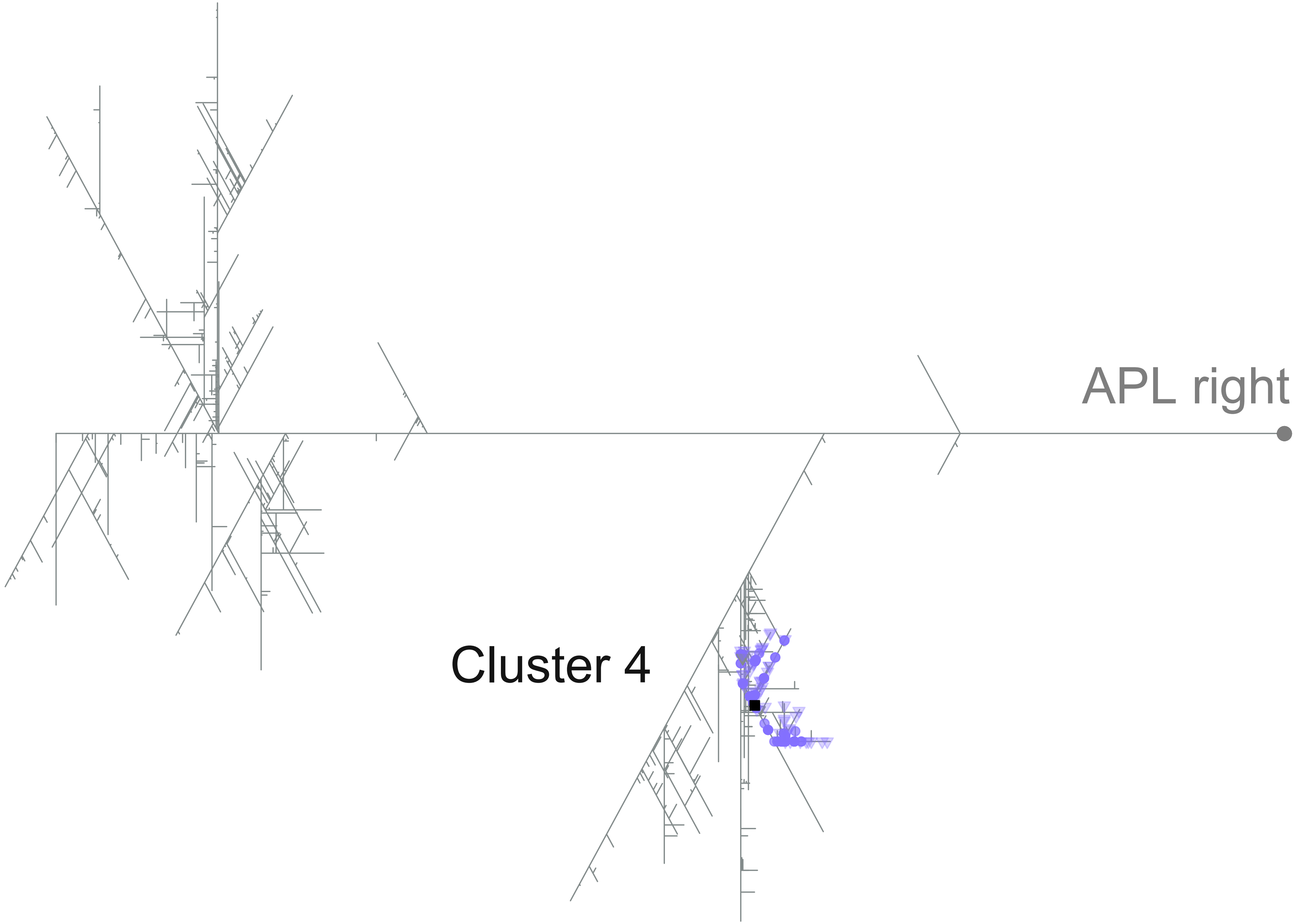
APL right
Cluster 4

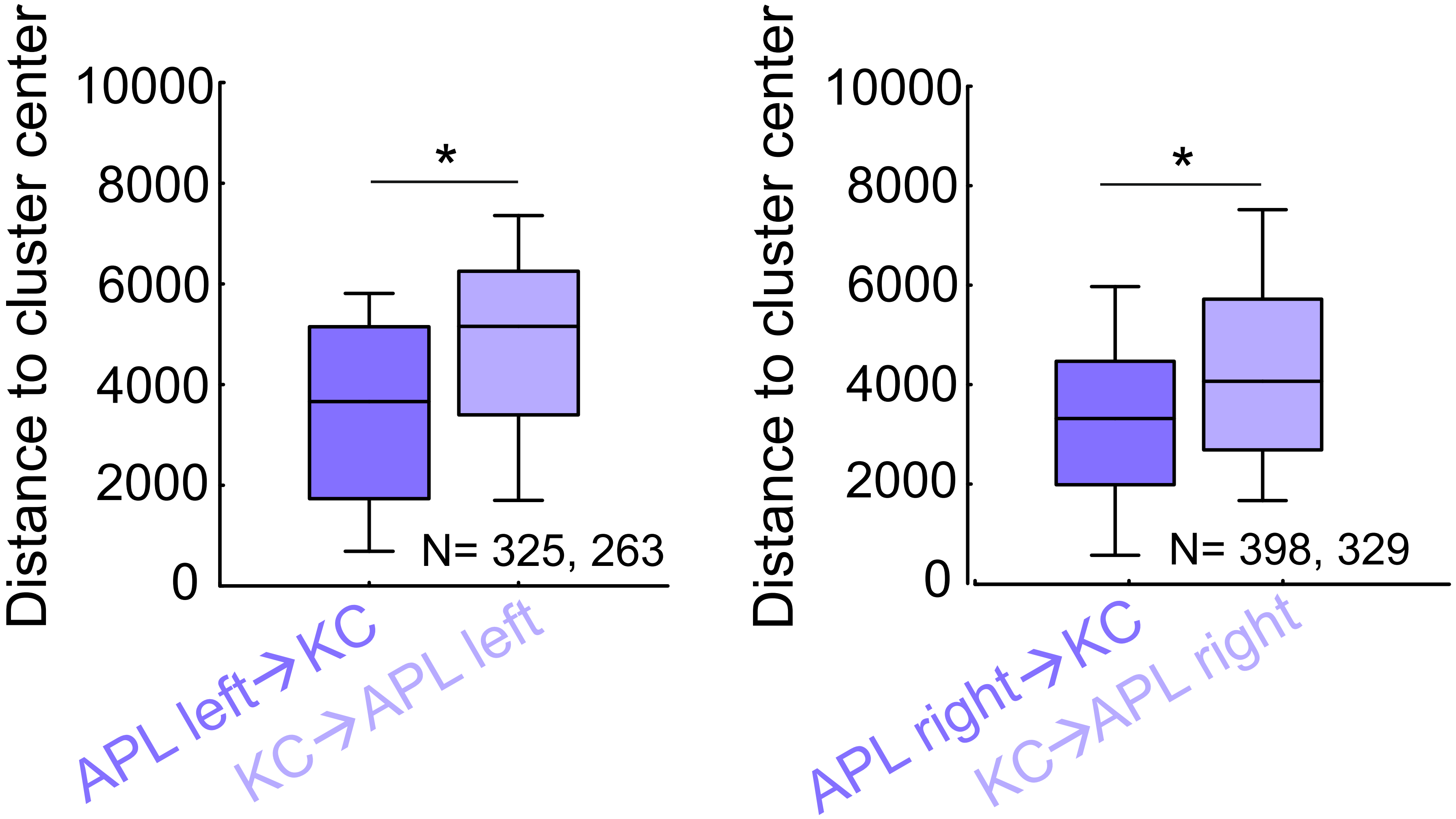

Distance to cluster center
10000
8000
6000
4000
2000
0
*
N= 325, 263
APL left→KC
KC→APL left
Distance to cluster center
10000
8000
6000
4000
2000
0
*
N= 398, 329
APL right→KC
KC→APL right
