## Extended Data Figure 16-1 for "Rewarding capacity of optogenetically activating a giant GABAergic central-brain interneuron in larval *Drosophila*"

Mancini et al.

### Extended Data Figure 16-1

**Expression patterns of the calyx MBON drivers used in Figure 16C-D. (A)** Full projection of the expression pattern from the SS02006-GAL4 driver (MBON-a1) covering only one calyx MBON in each hemisphere (N = 7 brains tested). **(B)** As in (A), but for SS01417-GAL4 (MBON-a2); in two out of N= 5 brains tested this driver covered both calyx MBONs (white arrowheads; left: cell bodies overlap one another). The data were acquired with a 63x glycerol objective; scale bar and grid spacing: 20 μm. Other details as in Figure 9C.

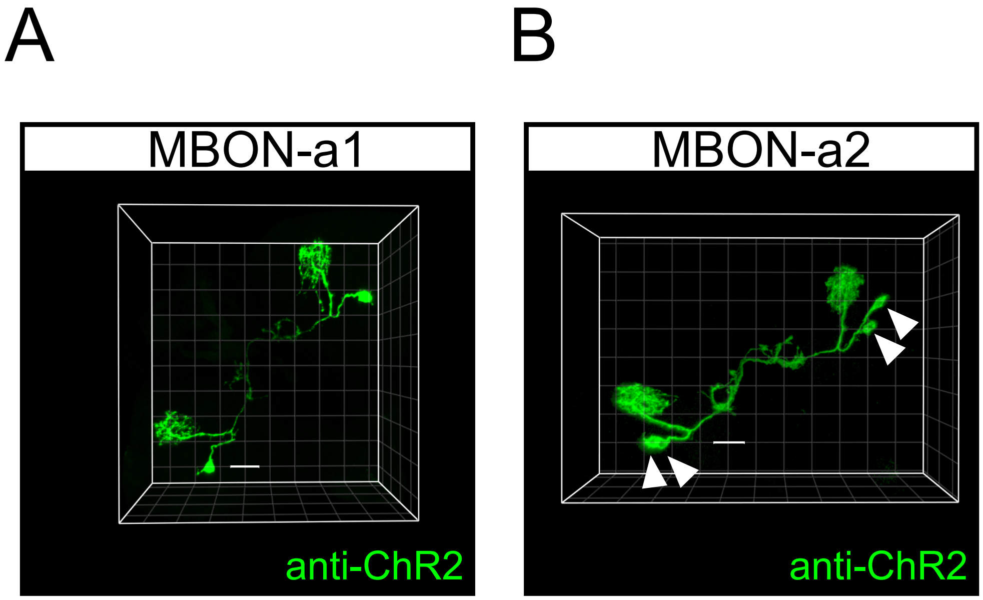
