## Extended Data Table 2-1 for "Rewarding capacity of optogenetically activating a giant GABAergic central-brain interneuron in larval *Drosophila*"

Mancini et al.

**Extended Data Table 2-1**

| REAGENT or RESOURCE | SOURCE or REFERENCE | IDENTIFIERS | ADDITIONAL INFORMATION |
| --- | --- | --- | --- |
| Fly strains | | | |
| SS01671-GAL4 (APL-GAL4; split-GAL4 driver covering specifically APL in larvae) | Saumweber et al., 2018 |  |  |
| APLi-GAL4 (intersectional driver covering specifically APL in larvae and adults) | Lin et al., 2014; Mayseless et al., 2018 |  |  |
| R36G04-GAL4 (MBONa1,a2-GAL4; GAL4 driver covering the calyx MBONs, plus additional neurons in the VNC in larvae) | Saumweber et al., 2018 | BDSC no. 49940 | chrs III |
| SS02006-GAL4 (MBONa1-GAL4; split-GAL4 driver covering specifically one of the two calyx MBONs in larvae) | Eschbach et al., 2020b; kindly provided by M. Zlatic, University of Cambridge |  |  |
| SS01417-GAL4 (MBONa2-GAL4; split-GAL4 driver covering one, or in some cases both calyx MBONs in larvae) | Eschbach et al., 2020b; kindly provided by M. Zlatic, University of Cambridge |  |  |
| R26G02-GAL4 (GAL4 driver covering APL, plus additional neurons in the VNC) | Jenett et al., 2012; Saumweber et al., 2018 | BDSC no. 48065 | chrs III |
| R55D08-GAL4 (GAL4 driver covering APL, plus additional neurons in the VNC) | Jenett et al 2012; Saumweber et al., 2018 | BDSC no. 39115 | chrs III |
| R14H06-LexA (LexA driver covering the KCs in larvae) | Pfeiffer et al., 2013; lexA Driver Collection of Rubin Laboratory at Janelia Farm | BDSC no. 52482 | chrs II |
| R58E02-LexA (LexA driver covering the pPAM neurons in larvae) | Pfeiffer et al., 2013; lexA Driver Collection of Rubin Laboratory at Janelia Farm | BDSC no. 52740 | chrs II |
| UAS-ChR2-XXL (optogenetic effector) | Dawydow et al., 2014 | BDSC no. 58374 | chrs II |
| UAS-ChR2-XXL-td::tomato (reporter/optogenetic effector) | Saumweber et al., 2018 | FlyBase ID: FBtp0131815 | chrs II |
| UAS-mCherry-CAAX (reporter effector) | Sens et al., 2010; Kobler et al., 2020 | BDSC no. 59021 | chrs II |
| UAS-mIFP-T2A-HO1 (reporter effector) | Yu et al., 2015; Kobler et al., 2020 | BDSC no. 64181 | chrs III |
| UAS-mIFP/MB247>mCherry-CAAX (recombined reporter effector/enhancer>effector ) | Kobler et al., 2020 |  |  |
| UAS-mCD8::GFP (reporter effector) | Lee and Luo 1999 | BDSC no. 5137 | chrs II |
| 20xUAS-IVS-CsChrimson::mVenus (reporter/optogenetic effector) | Klapoetke et al., 2014 | BDSC no. 55136 | chrs III |
| UAS-GtACR1::YFP (reporter/optogenetic effector) | Kindly provided by R. Kittel, Würzburg; Koenig et al., 2019 | BDSC no. 9736 | chrs II |
| lexAop-GCaMP6m (reporter effector) | Chen et al., 2013; Lyutova et al., 2019 | BDSC no. 44276 | chrs III |
| lexAop-reaper (pro-apoptotic effector) | Herranz et al., 2014; Lyutova et al., 2019 |  | chrs III |
| UAS-Dsyd-1::GFP (pre-synaptic reporter) | Owald et al., 2015 |  | chrs III |
| UAS-DenMark (post-synaptic reporter) | Nicolai et al., 2010 | BDSC no. 33062 | chrs II |
| UAS-Syt1:SNAP (pre-synaptic TAG reporter) | Kohl et al., 2014 | BDSC no. 58379 | Chrs III |
| UAS-TLN:CLIP (post-synaptic TAG reporter) | Kohl et al., 2014 | BDSC no. 58382 | Chrs III |
| w1118 |  | BDSC no. 3605, 5905, 6326 |  |
| y1w1 |  | BDSC no. 1495 |  |
| attP40/attP2 | Pfeiffer et al., 2010 |  |  |
| Antibodies | | | |
| primary monoclonal mouse anti-FASII | DSHB | 1D4 anti-Fasciclin II; AB_528235 | 1:50 |
| primary monoclonal mouse anti-ChR2 | ProGen Biotechnik | 610180 | 1:100 |
| primary polyclonal rabbit anti-GFP | Life Technologies | A6455 | 1:1000 |
| primary polyclonal FITC-conjugated goat anti-GFP | Abcam | ab 6662 | 1:1000 |
| primary monoclonal mouse 4F3 anti-DLG | Hybridoma | AB_528203 | 1:200 |
| primary polyclonal rabbit anti-DsRed | Clontech | 632496 | 1:200 |
| primary monoclonal rat anti-N-Cadherin | Hybridoma | DN-Ex #8-s | 1:50 |
| primary polyclonal rabbit anti-GABA | Sigma Aldrich | A2052 | 1:500 |
| primary polyclonal chicken anti-GFP | Aves Labs | AB_10000240 | 1:500 |
| secondary polyclonal goat anti-rabbit Alexa Fluor 488 | Life Technologies | A11008 | 1:500 |
| secondary polyclonal goat anti-mouse Alexa Fluor 568 | Life Technologies | A10037 | 1:500 |
| secondary polyclonal goat anti-rat Alexa Fluor 647 | Jackson IR | 712-605-153 | 1:500 |
| secondary polyclonal donkey anti-mouse Cy3 | Jackson IR | 715-165-150 | 1:300 |
| secondary polyclonal goat anti-mouse Alexa Fluor 488 | Invitrogen | A11001 | 1:200 |
| secondary polyclonal goat anti-rabbit Cy5 | Life Technologies | A10523 | 1:200 |
| secondary polyclonal goat anti-rat Cy3 | Life Technologies | A10522 | 1:200 |
| secondary polyclonal FITC-conjugated goat anti-chicken | Invitrogen | A16055 | 1:300 |
| Chemical TAG ligands (chemical substrates) | | | |
| SNAP-tag ligands (SNAP surface 549 - BG 549) | NEB | S9112S |  |
| CLIP-tag ligands (CLIP surface 647 - BC 647) | NEB | S9234S |  |
| Odors | | | |
| n-amyl acetate (AM) | Merck | 628-63-7 |  |
| 1-octanol (OCT) | Sigma Aldrich | 111-87-5 |  |
| Drugs | | | |
| carbamylcholine | Sigma-Aldrich | 51-83-2 |  |
| 3-Iodo-L-tyrosine (3IY) | Sigma–Aldrich | 70-78-0 |  |
| 3,4-dihydroxyphenylalanine (L-DOPA) | Sigma–Aldrich | 59-92-7 |  |
| Software | | | |
| Fiji ImageJ 1.53c | National Institutes of Health | SCR_002285 |  |
| CATMAID | Saalfeld et al., 2009; Schneider-Mizell et al., 2016 |  |  |
| Imaris 9.72, 9.8 | Oxford Instruments | SCR_007370 |  |
| ImSpector 7.1.4 | Miltenyi Biotec |  |  |
| R 3.3.2 | Development Core Team 2016 |  |  |
| Statistika 13 | StatSoft Inc | SCR_014213 |  |
| Corel Draw 2019 | Corel Corporation | SCR_013674 |  |
| GraphPad Prism 6 | GraphPad Software Inc. | SCR_002798 |  |
| Adobe Premiere Pro 2020 14.9.0 | Adobe Inc |  |  |
